# Genome-wide association study of musical beat synchronization demonstrates high polygenicity

**DOI:** 10.1101/836197

**Authors:** Maria Niarchou, Daniel E. Gustavson, J. Fah Sathirapongsasuti, Manuel Anglada-Tort, Else Eising, Eamonn Bell, Evonne McArthur, Peter Straub, The 23andMe Research Team, J. Devin McAuley, John A. Capra, Fredrik Ullén, Nicole Creanza, Miriam A. Mosing, David Hinds, Lea K. Davis, Nori Jacoby, Reyna L. Gordon

**Affiliations:** Vanderbilt Genetics Institute, Vanderbilt University Medical Center, TN, USA; Division of Genetic Medicine, Department of Medicine, Vanderbilt University Medical Center, TN, USA; 23andMe, Inc., Sunnyvale, CA, USA; Computational Auditory Perception Research Group, Max Planck Institute for Empirical Aesthetics, Frankfurt, Germany; Department of Language and Genetics, Max Planck Institute for Psycholinguistics, Nijmegen, Netherlands; Department of Music, Columbia University, New York, NY, USA; Department of Computer Science, Durham University, Durham, UK; Department of Psychology, Michigan State University, MI, USA; Bakar Computational Health Sciences Institute and Department of Epidemiology & Biostatistics, University of California, San Francisco, CA, USA; Department of Neuroscience, Karolinska Institutet, Sweden; Department of Cognitive Neuropsychology, Max Planck Institute for Empirical Aesthetics, Frankfurt, Germany; Department of Biological Sciences, Vanderbilt University, Nashville, TN, USA; Melbourne School of Psychological Sciences, University of Melbourne, Australia; Department of Biomedical Informatics, Vanderbilt University Medical Center, TN, USA; Department of Psychiatry and Behavioral Sciences, Vanderbilt University Medical Center, TN, USA; Department of Molecular Physiology and Biophysics, Vanderbilt University, TN, USA; Department of Otolaryngology – Head & Neck Surgery, Vanderbilt University Medical Center, TN, USA; Department of Psychology, Vanderbilt University, TN, USA; Vanderbilt Brain Institute, Vanderbilt University, Nashville, TN, USA

## Abstract

Moving in synchrony to the beat is a fundamental component of musicality. Here, we conducted a genome-wide association study (GWAS) to identify common genetic variants associated with beat synchronization in 606,825 individuals. Beat synchronization exhibited a highly polygenic architecture, with sixty-nine loci reaching genome-wide significance (p<5×10^−8^) and SNP-based heritability (on the liability scale) of 13%-16%. Heritability was enriched for genes expressed in brain tissues, and for fetal and adult brain-specific gene regulatory elements, underscoring the role of central nervous system-expressed genes linked to the genetic basis of the trait. We performed validations of the self-report phenotype (through internet-based experiments) and of the GWAS (polygenic scores for beat synchronization were associated with patients algorithmically classified as musicians in medical records of a separate biobank). Genetic correlations with breathing function, motor function, processing speed, and chronotype suggest shared genetic architecture with beat synchronization and provide avenues for new phenotypic and genetic explorations.

## Introduction

Our tendency to perceive, create, and appreciate rhythms in a variety of contexts (e.g., speech, music, movement) is a key feature of the human experience^1–3^. Rhythmic patterns provide predictable and robust sensorimotor structure to every-day interactions^4,5^, helping guide our attention to communicatively important moments in time^6,7^. Even young children are sensitive to the social and linguistic signals carried by rhythm^8–10^ and parents use rhythmic vocalizations and synchronous movement (e.g., lullabies and rocking) to interact with their infants from birth^11,12^. Rhythmic musical interactions are structured around the percept of a stable periodic pulse (termed the “beat” in Western music and present in music of most cultures^1,13^, though its precise instantiation in musical structure varies cross-culturally^14,15^). While music in general and rhythmic structures in particular vary globally^15–17^, there is evidence that hierarchical beat structure of most music is robust to cultural transmission^2^ and indeed common in many types of music^1^.

*Beat perception and synchronization* (i.e. perceiving, predicting, and moving predictively in synchrony to a musical beat^18^) is an important feature of musical experiences across many human cultures and musical genres^1,19^. The predictive temporal mechanisms afforded by beat structure enhance general perceptual and learning processes in music, including melody perception and production, singing, and joint music-making^3,6^. While some features of rhythm perception and production vary across listeners from different cultures^13,19–21^, the same studies showed considerable consistencies across cultures for other features (e.g., preference for beat-based isochrony). Musicality (broadly encompassing musical behavior, music engagement and musical skill^22^) impacts society by supporting pro-social behavior^11,23^ and well-being^24^. Many have proposed that beat perception and synchronization evolved in humans to support communication and group cohesion^18,22,25,26^. In modern humans, beat perception and synchronization are predictive of language and literacy skills^27,28^ and are related to cognition, motor function, and social coordination^29^. Thus, the biology of beat synchronization has general importance for understanding human ability to perceive and predict natural rhythms, may have relevance for characterizing phenotypes such as developmental speech-language disorders which demonstrate associations with atypical rhythm^30^, and may further elucidate mechanisms of rhythm-based rehabilitation (e.g., for stroke and Parkinson’s^31^).

Neuroimaging findings highlight auditory-motor networks in the brain underlying beat perception and production^32^, during which there is precise entrainment of neural oscillatory activity to musical signals, primarily involving motor planning areas and auditory regions of the brain, even during passive listening to music^33,34^. Neural mechanisms of entrainment, prediction, and reward work in concert to coordinate the timing of beat-related expectancies to musical signals during listening, playing, singing, and dance^26,34^. The significant inter-individual variation of beat synchronization^35^ are thought to be influenced, in part, by common genetic variation; thus genetic approaches can be used to gain a foothold on the biological basis of musicality and human rhythm traits.

Indeed, twin-modelling and other family-based studies point to moderate heritability of rhythm-related traits such as duration discrimination^36,37^, rhythm discrimination^38^, isochronous sensori-motor production^39^, and off-beat detection^40^. Much less is known at the molecular level about human genome variation underlying rhythm, and more generally musicality^41^ which to date has been investigated in relatively small samples^37^, due to the challenge of assessing such phenotypes in samples large enough to provide sufficient power to detect common variants with small effects (as expected for complex traits^42^). Large-scale genome-wide association studies (GWASs) of rhythm-based traits (e.g. beat synchronization) are thus needed to advance this field. Our understanding of the biological underpinnings of beat synchronization, from its genetic architecture to its neural instantiation, behavioral manifestation, and relationship to health, requires mechanistic multi-methodological approaches. Post-GWAS approaches (i.e., enrichment of gene expression in central nervous system tissues and genetic correlations) can be deployed to illuminate the relationship between the genetic and neural architecture of music-related traits, and shared underlying biology with other health traits.

Here, we report a large-scale genome-wide interrogation of beat synchronization. Our approach was as follows (Supplementary Figure 1): 1) We validated a subjective self-reported beat synchronization item (“Can you clap in time with a musical beat?”, referred to in this paper as the “target question”), in relation to measured beat synchronization and rhythm perception task performance. 2) We performed a GWAS in N=606,825 to identify genomic loci associated with beat synchronization. 3) We further investigated the genetic architecture of beat synchronization by estimating SNP-based heritability, partitioned heritability, and conducting gene property and gene set enrichment analyses. Lastly, we evaluated the contribution of genomic regions that have experienced significant human-specific evolutionary shifts. 4) We then validated GWAS results by testing whether a cumulative sum of the genetic effects for beat synchronization detected in our GWAS (i.e., polygenic score or PGS) was significantly associated with algorithmically identified musical engagement in a separate sample. 5) We explored shared genetic effects (pleiotropy) on beat synchronization and other traits through genetic correlation and genomic structural equation analyses.

## Results

### Overview. Validating the self-reported beat synchronization phenotype

In light of prior work suggesting that musicality and rhythm skills are complex traits that can be quantified with both objective (experiment-derived) assessment and subjective self-reported data^43,44^, we performed a series of validations of the GWAS target question (i.e., the self-report “Can you clap in time with a musical beat?”), in relation to rhythm perception and beat production tasks. Both studies were administered in English for consistency. We also explored the relationship between task-based beat synchronization ability, a self-reported rhythm scale, and musicality. Study overviews and key results are summarized in Figure 1.

**Figure 1.**
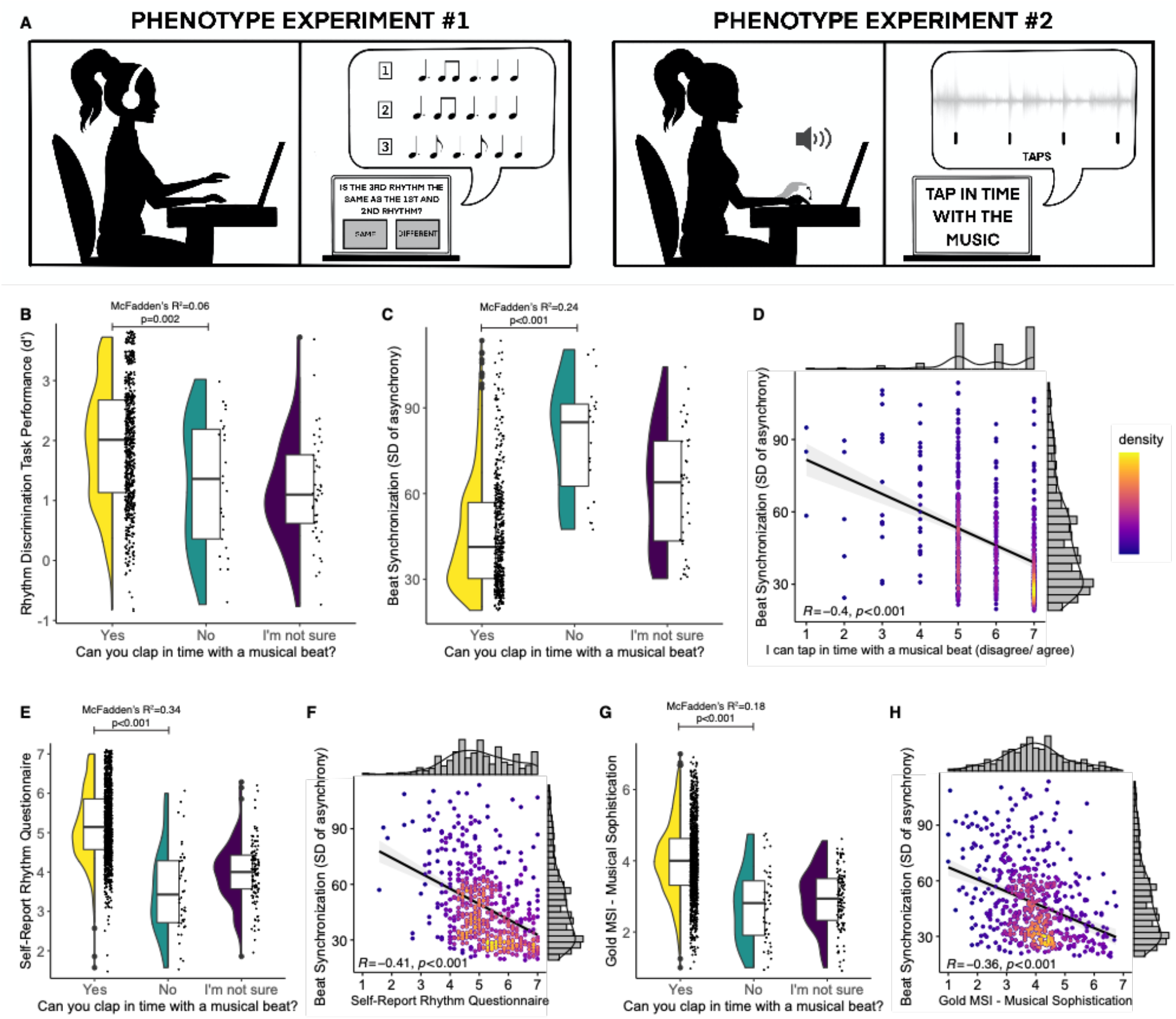
Phenotype validation studies overview and results. <u>A) Schema of internet-based phenotype validation studies</u>. In phenotype experiment #1, participants performed a musical rhythm perception test and provided self-report of the same target question in the GWAS study (“Can you clap in time with a musical beat?”). In phenotype experiment #2, participants performed beat synchronization tasks (which involved tapping to the beat of musical excerpts) as well as responding to the same target question, in addition to a series of other questionnaires about their musical engagement/ability and health traits. B) <u>Phenotype Experiment 1 results</u> in N=724 show rhythm perception task performance is correlated with Yes vs. No responses to GWAS target question, Odds Ratio (OR)=1.92, McFadden’s R^2^ = 0.06, *p*=0.002. <u>C-H): Phenotype Experiment 2 results</u>. C) Beat synchronization task performance (n=542) is highly correlated with Yes vs. No responses to the target question in OR=0.28, McFadden’s R^2^ =0.24, p<0.001; note that lower values of SD of the asynchrony correspond to more accurate tapping in time to the musical beat. D) Beat synchronization task performance is correlated with responses to a similar self-report question asked on a Likert scale, in n=542, *r*=–0.40, p<0.001. E) Self-reported rhythm questionnaire (seven-item scale in N=1,412) is correlated with responses to the target question, McFadden’s R^2^ =0.34, p<0.001. F) Beat synchronization task performance is correlated with Self-reported rhythm questionnaire in n=542, *r*=0.41, p<0.001. G) Gold-MSI (self-reported musical sophistication questionnaire) is correlated with responses to the target question in N=1,412, OR=4.16, McFadden’s R^2^ = 0.18, p<0.001. H) Beat synchronization task performance is correlated with Gold-MSI in n=542, r=-0.36, p<0.001. Within each plot for panels B,C,E and G, distributions are displayed using violin plots (mirrored density plot showing probability density on the left), jittered individual data plots (right), and box plots in the center (horizontal line at median, lower and upper hinges correspond to the first and third quartiles. The upper and lower whisker extends from the hinges to the value no further than 1.5 * interquartile range from the hinge). Data beyond the end of the whiskers are called “outlying” points and are plotted individually. Panels D, F, and H scatterplots are shown with dots colored by density to illustrate distribution. Taken together, these results show that self-reported beat synchronization is a reasonable proxy of the trait.

#### Phenotype Experiment 1: Rhythm perception task performance

In this experiment N=724 (see Table 1 for demographics) were asked the target question and performed a musical rhythm perception test (Supplementary Figure 2). In each of the 32 trials of the task, participants judged whether two rhythms were the same or different (see Figure 1A), following a standard procedure for assessing musical perception ability^45^ and utilizing rhythm sequences with simple (highly metrical) and complex (syncopated) rhythms^46^. The rhythm perception task yielded quantitative scores (*d’*). Individuals with better performance in the rhythm perception test (higher total *d’*) were more likely to answer Yes (vs. No) to the target question (OR=1.92, p=0.002, McFadden’s R^2^=0.06, 95% CI=1.27,2.95; Figure 1B). All tests in both phenotype experiments were two-tailed.

**Table 1.** Sample demographics for each of the three study samples (GWAS and Phenotype Experiments 1 and 2).

**GWAS Sample by Phenotype group (response to Clap-to-beat question)**
|  | *Cases (Yes)* | *Controls (No)* |
| --- | --- | --- |
| Total | 555660 | 51165 |
| Males | 226188 | 23998 |
| Females | 329472 | 27167 |
| 18 to 30 years old | 57898 | 5186 |
| 30 to 45 years old | 135168 | 12909 |
| 45 to 60 years old | 150939 | 13312 |
| 60 years old and over | 211655 | 19758 |

**Phenotype Validation Experiment 1 (rhythm perception)**
| *Full sample who provided demographics* | *N* | *Mean Age in years (for N=722 who reported demographics)* | *SD Age* |
| --- | --- | --- | --- |
| Total | 722 | 36.08 | 10.90 |
| Males | 386 | 34.95 | 10.60 |
| Females | 332 | 37.49 | 11.07 |

**Phenotype Validation Experiment 2 (beat synchronization and cross-trait phenotypic replication)**
| *Full sample (questionnaires)* | *N* | *Mean Age in years* | *SD Age* |
| --- | --- | --- | --- |
| Total | 1412 | 36.34 | 11.93 |
| Males | 678 | 35.53 | 11.12 |
| Females | 728 | 37.15 | 12.61 |
| *Subset with valid tapping data* | *n* | *Mean Age in years* | *SD Age* |
| Total | 542 | 35.24 | 11.39 |
| Males | 241 | 35.02 | 10.62 |
| Females | 300 | 35.43 | 12.00 |

#### Phenotype Experiment 2: Beat synchronization task performance

We then validated self-reported beat synchronization phenotype (N=1,412) as a proxy for directly-measured beat synchronization ability. Participants (Table 1) completed a questionnaire on musicality, health, and personality, and were asked to tap in real time to the beat of 4 different musical excerpts (Supplementary Figure 3). Beat synchronization tapping accuracy was assessed similarly to lab-based studies^35^, but with a recently developed online-based technology that precisely measures asynchrony of participants’ taps along to music clips - i.e., REPP (Rhythm ExPeriment Platform^47^) for additional details and pre-registered hypotheses (H1-H6), see Methods and Supplementary Notes. Key results of this study are summarized in Figure 1 and Supplementary Table 1. Note that more accurate tapping is reflected in lower tapping asynchrony scores, i.e., more accurate timing of taps in relation to the beat.

As predicted (OSF pre-registered H1), individuals who responded Yes to the target question (“Can you clap in time with a musical beat”) had lower tapping asynchrony, OR=0.28, p<.001, McFadden’s R^2^=.24, 95% CI=0.18,0.42 (Figure 1C). Tapping asynchrony was also negatively correlated with responses to a highly similar item (“I can tap in time to a musical beat”) when asked on a seven-point Likert agreement scale (1= disagree; 7 = agree), *r*= -.40, p<.001, 95% CI=-0.47,-0.33] (H1a; Figure 1D). Similarly, individuals with higher self-reported rhythmic ability (from another multi-item questionnaire) were much more likely to respond “Yes” to the target question, OR=7.34, p<.001, McFadden’s R^2^=.34, 95% CI=4.90,11.52], (Figure 1E), and demonstrate lower tapping asynchrony, r = -.41, p <.001, 95% CI=-0.47,-0.33] (Figure 1F) (H2). Controlling for confidence judgments or confidence as a personality trait did not diminish the associations between self-report and tapping asynchrony (H3; Supplementary Notes). Musical Sophistication^43^ was positively associated with the target question, OR=4.16, p <.001, McFadden’s R^2^=.18, 95% CI=2.90,6.12 (Figure 1G) and negatively correlated with tapping asynchrony r= -.36, p <.001, 95% CI =-0.43,-0.28 (Figure 1H; H5). There was no credible evidence that Musical Sophistication or prior/current musician status interacted with the tapping asynchrony to predict responses to the target question (H6). All associations reported here were maintained when controlling for age, sex, and education (Supplementary Table 1). Key analyses were repeated using vector length (variability of relative phase of participants’ tapping) as an outcome, and showed the same pattern of results as SD of the asynchrony (Methods and Supplementary Notes). Taken together, results show that the self-reported target question is a valid phenotype, and that other similar self-reported rhythm measures are also valid proxies of beat synchronization.

### Beat Synchronization GWAS

#### Genomic study population

The discovery GWAS sample consisted of N=606,825 unrelated participants of European ancestry (see Table 1 for demographics), who consented to participate in research with 23andMe, Inc. and answered Yes (91.57%) or No (8.43%) to the target question “Can you clap in time with a musical beat?”

#### GWAS results and SNP-based heritability estimation

GWAS was conducted using logistic regression under an additive genetic model, while adjusting for age, sex, the first five principal components from genetic data, and genotype platforms (Methods). Seventy “sentinel” SNPs (after two rounds of LD pruning, first at R^2^=0.6 and then at R^2^=0.1, kb = 250) at 69 genomic loci reached genome-wide significance (p<5×10^−8^; two-tailed; Figure 2, Table 2, and Supplementary Table 2), with a total of 6,160 SNPs passing the genome-wide significance threshold. Sixty-seven loci were autosomal and two were on the X chromosome; locus 28 contains two independent sentinel SNPs. QQ-plot is provided in Supplementary Figure 4, and local association plots at each locus are in the Regional Plots Supplemental document. The LD score regression intercept was 1.02 (se=0.01) the ratio was 0.03, indicating that the majority of inflation in test statistics was due to true polygenicity instead of population stratification.

**Table 2.**
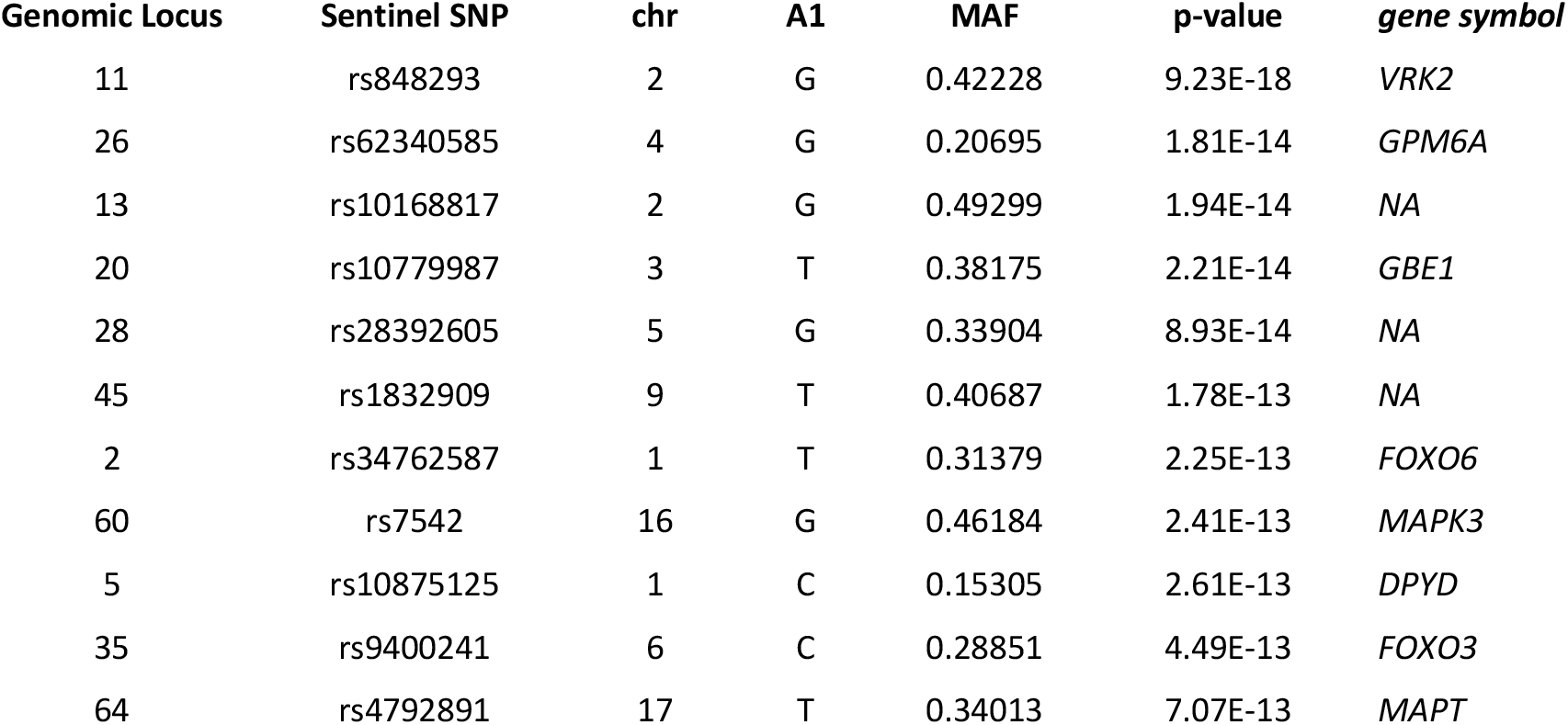
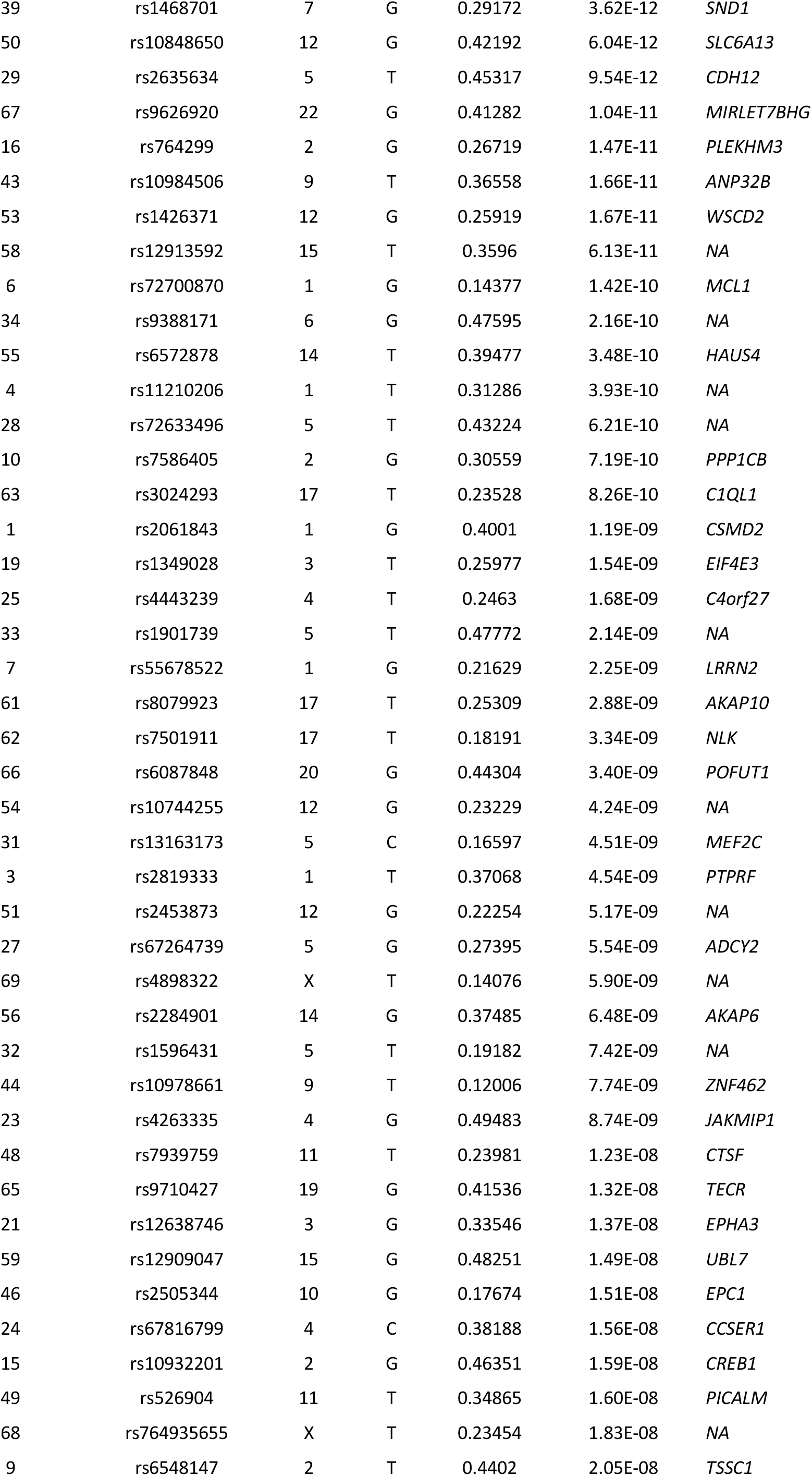
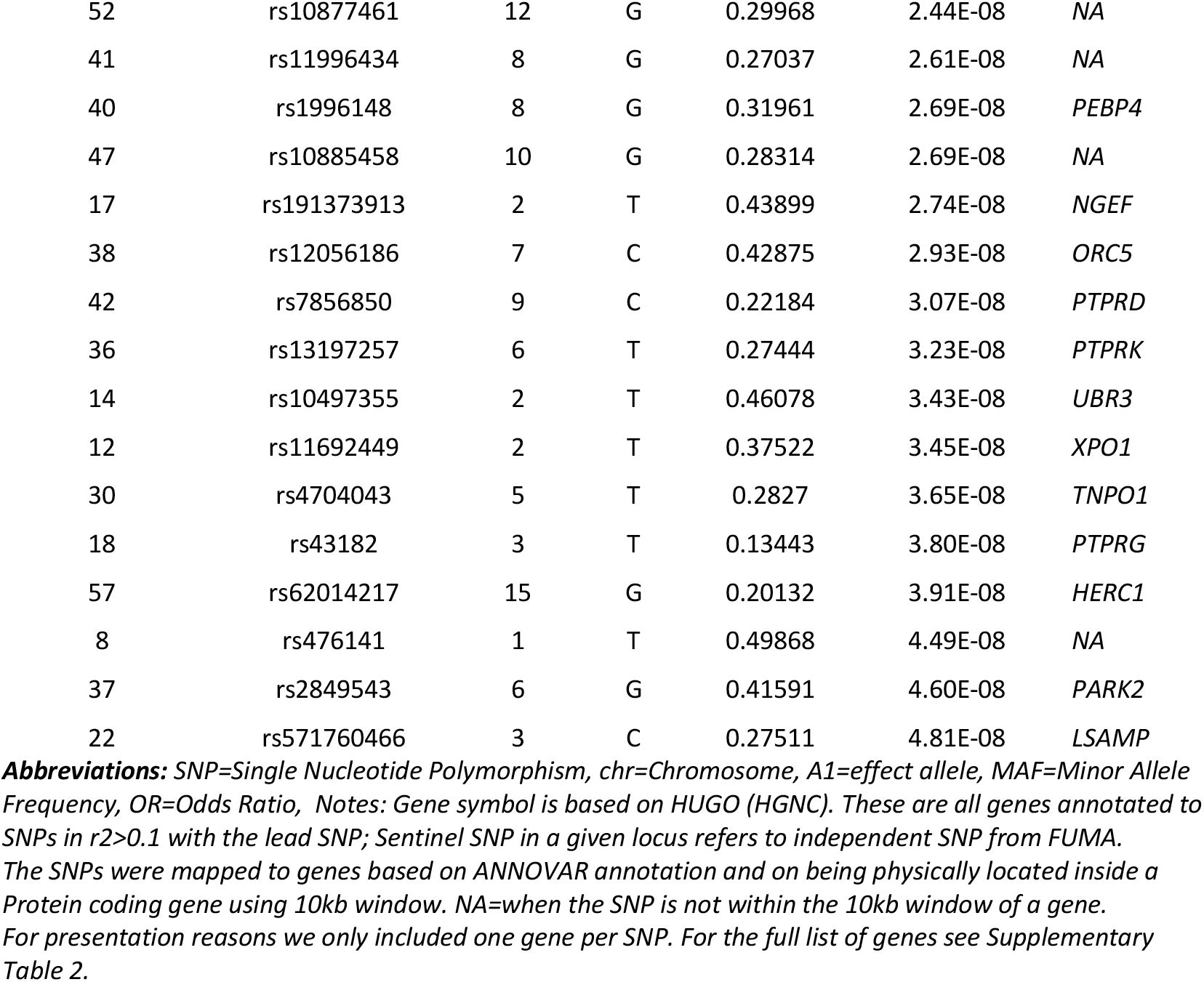
Genomic loci and sentinel SNPs significantly associated with beat synchronization in the primary GWAS. Further details (e.g., chromosomal location) are provided in Supplementary Table 2.

| **Genomic Locus** | **Sentinel SNP** | **chr** | **A1** | **MAF** | **p-value** | **gene symbol** |
| --- | --- | --- | --- | --- | --- | --- |
| 11 | rs848293 | 2 | G | 0.42228 | 9.23E-18 | VRK2 |
| 26 | rs62340585 | 4 | G | 0.20695 | 1.81E-14 | GPM6A |
| 13 | rs10168817 | 2 | G | 0.49299 | 1.94E-14 | NA |
| 20 | rs10779987 | 3 | T | 0.38175 | 2.21E-14 | GBE1 |
| 28 | rs28392605 | 5 | G | 0.33904 | 8.93E-14 | NA |
| 45 | rs1832909 | 9 | T | 0.40687 | 1.78E-13 | NA |
| 2 | rs34762587 | 1 | T | 0.31379 | 2.25E-13 | FOXO6 |
| 60 | rs7542 | 16 | G | 0.46184 | 2.41E-13 | MAPK3 |
| 5 | rs10875125 | 1 | C | 0.15305 | 2.61E-13 | DPYD |
| 35 | rs9400241 | 6 | C | 0.28851 | 4.49E-13 | FOXO3 |
| 64 | rs4792891 | 17 | T | 0.34013 | 7.07E-13 | MAPT |
| 39 | rs1468701 | 7 | G | 0.29172 | 3.62E-12 | *SND1* |
| 50 | rs10848650 | 12 | G | 0.42192 | 6.04E-12 | *SLC6A13* |
| 29 | rs2635634 | 5 | T | 0.45317 | 9.54E-12 | *CDH12* |
| 67 | rs9626920 | 22 | G | 0.41282 | 1.04E-11 | *MIRLET7BHG* |
| 16 | rs764299 | 2 | G | 0.26719 | 1.47E-11 | *PLEKHM3* |
| 43 | rs10984506 | 9 | T | 0.36558 | 1.66E-11 | *ANP32B* |
| 53 | rs1426371 | 12 | G | 0.25919 | 1.67E-11 | *WSCD2* |
| 58 | rs12913592 | 15 | T | 0.3596 | 6.13E-11 | *NA* |
| 6 | rs72700870 | 1 | G | 0.14377 | 1.42E-10 | *MCL1* |
| 34 | rs9388171 | 6 | G | 0.47595 | 2.16E-10 | *NA* |
| 55 | rs6572878 | 14 | T | 0.39477 | 3.48E-10 | *HAUS4* |
| 4 | rs11210206 | 1 | T | 0.31286 | 3.93E-10 | *NA* |
| 28 | rs72633496 | 5 | T | 0.43224 | 6.21E-10 | *NA* |
| 10 | rs7586405 | 2 | G | 0.30559 | 7.19E-10 | *PPP1CB* |
| 63 | rs3024293 | 17 | T | 0.23528 | 8.26E-10 | *C1QL1* |
| 1 | rs2061843 | 1 | G | 0.4001 | 1.19E-09 | *CSMD2* |
| 19 | rs1349028 | 3 | T | 0.25977 | 1.54E-09 | *EIF4E3* |
| 25 | rs4443239 | 4 | T | 0.2463 | 1.68E-09 | *C4orf27* |
| 33 | rs1901739 | 5 | T | 0.47772 | 2.14E-09 | *NA* |
| 7 | rs55678522 | 1 | G | 0.21629 | 2.25E-09 | *LRRN2* |
| 61 | rs8079923 | 17 | T | 0.25309 | 2.88E-09 | *AKAP10* |
| 62 | rs7501911 | 17 | T | 0.18191 | 3.34E-09 | *NLK* |
| 66 | rs6087848 | 20 | G | 0.44304 | 3.40E-09 | *POFUT1* |
| 54 | rs10744255 | 12 | G | 0.23229 | 4.24E-09 | *NA* |
| 31 | rs13163173 | 5 | C | 0.16597 | 4.51E-09 | *MEF2C* |
| 3 | rs2819333 | 1 | T | 0.37068 | 4.54E-09 | *PTPRF* |
| 51 | rs2453873 | 12 | G | 0.22254 | 5.17E-09 | *NA* |
| 27 | rs67264739 | 5 | G | 0.27395 | 5.54E-09 | *ADCY2* |
| 69 | rs4898322 | X | T | 0.14076 | 5.90E-09 | *NA* |
| 56 | rs2284901 | 14 | G | 0.37485 | 6.48E-09 | *AKAP6* |
| 32 | rs1596431 | 5 | T | 0.19182 | 7.42E-09 | *NA* |
| 44 | rs10978661 | 9 | T | 0.12006 | 7.74E-09 | *ZNF462* |
| 23 | rs4263335 | 4 | G | 0.49483 | 8.74E-09 | *JAKMIP1* |
| 48 | rs7939759 | 11 | T | 0.23981 | 1.23E-08 | *CTSF* |
| 65 | rs9710427 | 19 | G | 0.41536 | 1.32E-08 | *TECR* |
| 21 | rs12638746 | 3 | G | 0.33546 | 1.37E-08 | *EPHA3* |
| 59 | rs12909047 | 15 | G | 0.48251 | 1.49E-08 | *UBL7* |
| 46 | rs2505344 | 10 | G | 0.17674 | 1.51E-08 | *EPC1* |
| 24 | rs67816799 | 4 | C | 0.38188 | 1.56E-08 | *CCSER1* |
| 15 | rs10932201 | 2 | G | 0.46351 | 1.59E-08 | *CREB1* |
| 49 | rs526904 | 11 | T | 0.34865 | 1.60E-08 | *PICALM* |
| 68 | rs764935655 | X | T | 0.23454 | 1.83E-08 | *NA* |
| 9 | rs6548147 | 2 | T | 0.4402 | 2.05E-08 | *TSSC1* |
| 52 | rs10877461 | 12 | G | 0.29968 | 2.44E-08 | NA |
| 41 | rs11996434 | 8 | G | 0.27037 | 2.61E-08 | NA |
| 40 | rs1996148 | 8 | G | 0.31961 | 2.69E-08 | PEBP4 |
| 47 | rs10885458 | 10 | G | 0.28314 | 2.69E-08 | NA |
| 17 | rs191373913 | 2 | T | 0.43899 | 2.74E-08 | NGEF |
| 38 | rs12056186 | 7 | C | 0.42875 | 2.93E-08 | ORC5 |
| 42 | rs7856850 | 9 | C | 0.22184 | 3.07E-08 | PTPRD |
| 36 | rs13197257 | 6 | T | 0.27444 | 3.23E-08 | PTPRK |
| 14 | rs10497355 | 2 | T | 0.46078 | 3.43E-08 | UBR3 |
| 12 | rs11692449 | 2 | T | 0.37522 | 3.45E-08 | XPO1 |
| 30 | rs4704043 | 5 | T | 0.2827 | 3.65E-08 | TNPO1 |
| 18 | rs43182 | 3 | T | 0.13443 | 3.80E-08 | PTPRG |
| 57 | rs62014217 | 15 | G | 0.20132 | 3.91E-08 | HERC1 |
| 8 | rs476141 | 1 | T | 0.49868 | 4.49E-08 | NA |
| 37 | rs2849543 | 6 | G | 0.41591 | 4.60E-08 | PARK2 |
| 22 | rs571760466 | 3 | C | 0.27511 | 4.81E-08 | LSAMP |

**Abbreviations:** SNP=Single Nucleotide Polymorphism, chr=Chromosome, A1=effect allele, MAF=Minor Allele Frequency, OR=Odds Ratio, Notes: Gene symbol is based on HUGO (HGNC). These are all genes annotated to SNPs in $r^2 > 0.1$ with the lead SNP; Sentinel SNP in a given locus refers to independent SNP from FUMA. The SNPs were mapped to genes based on ANNOVAR annotation and on being physically located inside a Protein coding gene using 10kb window. NA=when the SNP is not within the 10kb window of a gene. For presentation reasons we only included one gene per SNP. For the full list of genes see Supplementary Table 2.

**Figure 2.**
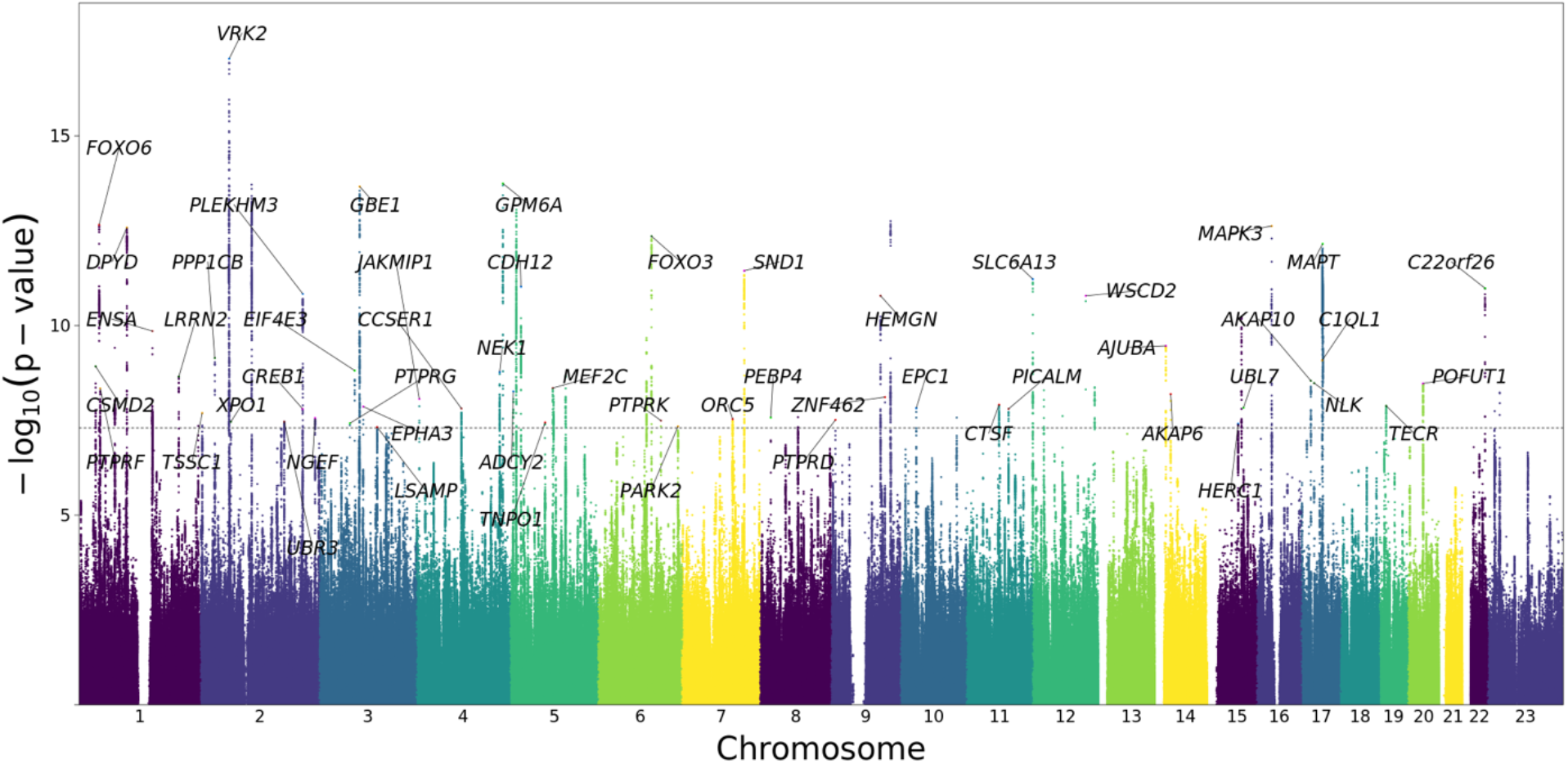
Manhattan plot of GWAS results of beat synchronization. Results of GWAS in N=606,825 with 23andMe. The GWAS phenotype is participants’ responses of Yes (N=555,660) vs. No (N=51,165) to the question “Can you clap in time with a musical beat?”. The GWAS controlled for age, sex, top 5 PC’s for ancestry, and genotype platform. The x-axis shows chromosomal position and the y-axis shows -log10 p-values). Sixty-nine loci (70 sentinel SNPs, with one locus containing two independent sentinel SNPs) surpassed the threshold for genome-wide significance of p<5×10^−8^ (dotted horizontal line). For illustration purposes, only 500,000 SNPs with p<0.1 are shown; gene symbols for sentinel SNPs are notated when FUMA provided a gene mapped to nearest sentinel SNP.

The top-associated locus (rs848293) was mapped at chromosome 2 close to *VRK2 (*Vaccinia Serine/Threonine Kinase 2 which codes for a protein kinase with multiple spliced isoforms expressed in the brain) and *FANCL*, within a region previously linked to multiple neurological phenotypes^48,49^. Another strongly associated locus at chromosome 17 (rs4792891) included the Microtubule Associated Protein Tau (*MAPT*) gene, a Parkinson’s disease^50^ associated locus. The Mitogen-Activated Protein Kinase 3 (*MAPK3*) gene at 16p11.2, a region known to harbor rare variants which influence neurodevelopmental disorders ^51^ and language-related phenotypes^52^, was also strongly implicated. We also identified a locus at Glycoprotein M6A (*GPM6A*), whose gene promoter contains a transcription factor binding site for *GATA2*, a gene previously related to music phenotypes^37^.

SNP-based heritability estimates on the liability scale^53^ ranged from 13% to 16% when adjusted for a range of estimated population prevalence for atypical beat synchronization (3.0% to 6.5%; Supplementary Table 3; see Supplementary Notes for explanation of prevalence estimates). The observed (unadjusted) genetic variance explained 5% (se=0.002) of the phenotypic variance.

#### Gene based GWAS

Gene-based genome-wide association analyses performed with MAGMA yielded 129 genes surpassing the threshold of p<2.56×10^−6^ (two-tailed; Supplementary Table 4), with top two hits at: *CCSER1*, in the 4q22 region in proximity to genes previously associated with multiple musicality phenotypes^54^, and *VRK2* (converging with the top locus in our SNP-based GWAS). Within these associations, we examined potential replication of 29 genetic associations with musicality in humans from prior reports^37,54,55^; none reached significance after genome-wide correction (Supplementary Table 5, Supplementary Notes), neither independently, nor as a gene-set (p=0.297).

### In silico functional analyses

#### Gene property and gene set enrichment analyses

To understand the biological functions and gene expression associations of beat synchronization, we performed gene set analysis (GSA) and gene property enrichment analyses^56^ on the gene-based p-values, using MAGMA^57^ implemented in FUMA^58^. Results of conditional gene property analysis (based on GTEx data tissue types^59^ and controlling for average expression) demonstrated that the genetic architecture of beat synchronization was significantly enriched in genes expressed in brain tissues (Figure 3A), including cortex, cerebellum, and basal ganglia (putamen, caudate and nucleus accumbens), converging with subcortical and cortical regions supporting beat perception and synchronization^34^.

**Figure 3.**
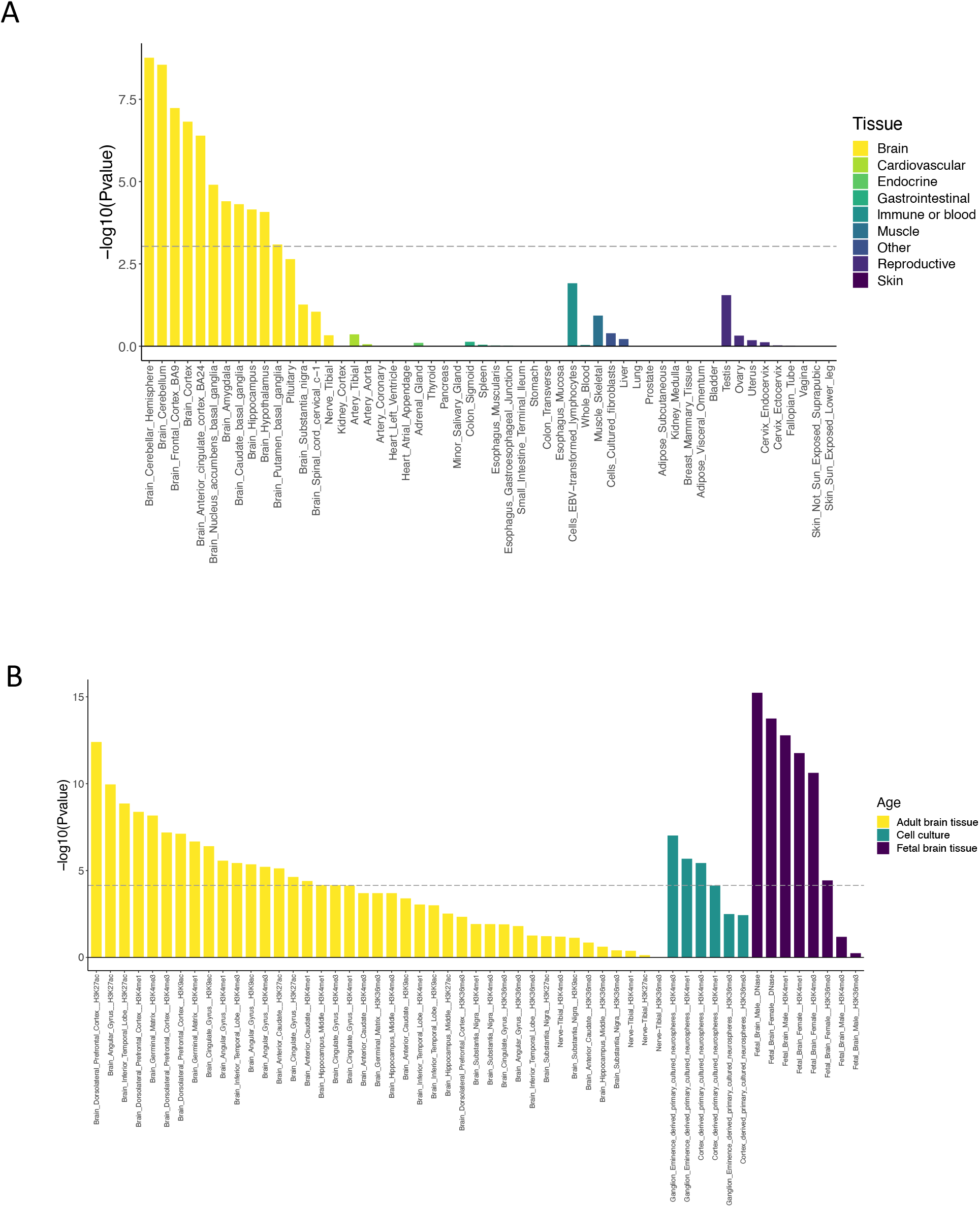
Genetic architecture of beat synchronization is enriched for brain-related expression. <u>A. Genes associated with beat synchronization are enriched for expression in brain tissue</u>. Results of MAGMA gene-property analysis are based on gene expression levels from GTEx v8, in 54 tissues, conditioned on average expression across tissues. Associations with beat synchronization were significantly enriched in brain-expressed genes (-log-10 p-values are on the y-axis, with tissue type on the x-axis). Dotted line shows p-value threshold for significant enrichment after Bonferroni correction. <u>B. Partitioned heritability shows enrichment in brain-specific regulatory regions of the genome.</u> Partitioned heritability analysis was performed with LDSC-SEG. Tissue-specific regulatory elements are marked by histone 3 acetylation or DNase hypersensitivity (for open chromatin) and H3K4me1 (for enhancers). Regulatory regions in adult brain tissues are shown in yellow, with regulatory elements in in cell cultures in teal, and in fetal brain tissue shown in dark purple. The graph shows -log-10 p-values are on y-axis, with tissue and marker type on x-axis. The dotted line shows p-value threshold for significant enrichment after Bonferroni correction for number of gene sets tested.

To further examine potential biological functions associated with beat synchronization, we performed exploratory GSA^57^ (Supplementary Table 6). The genetic architecture of beat synchronization was enriched for two gene sets associated with nervous system function: gene sets for synaptic membrane adhesion (p=1.01 × 10^−7^) and synaptic adhesion-like molecules (p=8.35×10^−7^).

#### Partitioned Heritability

Complementing these gene-based enrichment analyses, we also performed stratified LDSC^60^ on the GWAS results to partition heritability according to genomic properties, using specific functional categories to gain insight into the types of variation that contribute most to beat synchronization. Among broad SNP annotation categories^61^ (Supplementary Table 7), we found enrichment (all p<9.6×10^−4^) of: regions conserved in mammals (considered under purifying selection^62^), regulatory regions marked by acetylation of histone H3 at lysine 9 (H3K9ac; a marker for active chromatin, and monomethylation of histone H3 at lysine 4 (H3K4me1; a marker for enhancers), supporting the hypothesis that identified associations may affect gene regulation. We next used LDSC-Specifically Enriched Genes (LDSC-SEG^63^) to determine whether genes expressed in specific cell- or tissue-types (conditional to the other annotations) are enriched for beat synchronization-associated variants. For tissue-specific annotations of active chromatin and enhancers (marked by H3K9ac, H3K27ac, DNase hypersensitivity sites and H3K4me1), heritability was enriched in central-nervous-system- and skeletal muscle-specific regulatory regions (Supplementary Table 8). Cell-type specific, multi-tissue chromatin, and multi-tissue gene expression results are shown in Supplementary Figures 5, 6 and 7 respectively. Enrichment in brain-specific regulatory elements, in multiple fetal and adult tissue-specific elements as well as CNS-specific cell cultures, are shown in Figure 3B.

### Evolutionary Analyses

Given evolutionary hypotheses about the origins of rhythm^4,18,64^, we evaluated the contribution of regions of the human genome that have experienced significant human-specific shifts in evolutionary pressure, using stratified LDSC^53,60^. In particular, we analyzed the contribution to beat synchronization heritability from variants in genomic loci that are conserved across non-human species, but have elevated substitution rate on the human lineage^65^. Many of these human accelerated regions (HARs) play roles in human-enriched traits^66^, including cognition^67^. Two variants significantly (*p* < 5× 10^−8^) associated with beat synchronization (rs14316 at locus 66, rs1464791 at locus 20) fall within HARs. This is 11.2 times more overlap than expected by chance (µ=0.178 overlaps; p=0.017, based on 10,000 permutations). The rs1464791 variant is near *GBE1*, a gene associated with neuromuscular disease^68^, reaction time^69^ and cognitive impairments^70^. Applying LDSC to consider the full set of association statistics, we find that genetic variants in HARs contribute 2.26 times more to the observed heritability of beat synchronization than would be expected if heritability were distributed uniformly across variants (*p* = 0.14). Given the small number of common variants within HARs, this stratified heritability analysis is substantially underpowered (0.17% of variants considered are in HARs). The general agreement of these two approaches supports the enrichment of functional variation relevant to beat synchronization in HARs. We also evaluated the contribution of genetic variants detected in the Neanderthal genome to the heritability of beat synchronization (Supplementary Notes and Supplementary Table 9).

### Polygenic scores (PGS) for beat synchronization are related to musicality reported in a health care context

We investigated whether the polygenic model of beat synchronization from the GWAS would differentiate self-identified musicians from non-musicians in a separate sample. Musicians (n=1,259) and matched controls (n=4,893) were drawn from a study^71^ that algorithmically identified musically active patients (Methods and Supplementary Notes). PGS for beat synchronization were significantly associated with musical engagement (OR=1.33 per SD increase in PGS, p<2 × 10^−16^, Nagelkerke’s R^2^=2%, 95% CI=1.25,1.42) consistent with beat synchronization capturing a dimension of musicality.

### Cross-trait analyses

#### Genetic correlations

To determine if beat synchronization shares genetic architecture with other traits, we tested genetic correlations^72^ between beat synchronization GWAS and available GWAS of 64 traits classified into seven domains (Supplementary Table 10 and Supplementary Notes for details). There were 15 statistically significant genetic correlations (p<7.8×10^−4^) (Figure 4, Supplementary Table 11). Results included positive correlations with motor function (grip strength and usual walking pace) and heaviness of smoking, and negative correlations with risk-taking and smoking initiation. There were two correlations with hearing traits (positive correlation with tinnitus and negative correlation with hearing difficulty). From the cognitive traits, processing speed (faster perceptual motor speed) was genetically correlated with beat synchronization, in addition to executive function - shifting (from a GWAS of trail-making, a task that involves complex processing speed). There were multiple associations with other biological rhythms: breathing function traits (positive associations with peak expiratory flow, forced expiratory volume, forced vital capacity, and a negative correlation with shortness of breath) and negative associations with sleep-related traits (insomnia and morning chronotype).

**Figure 4.**
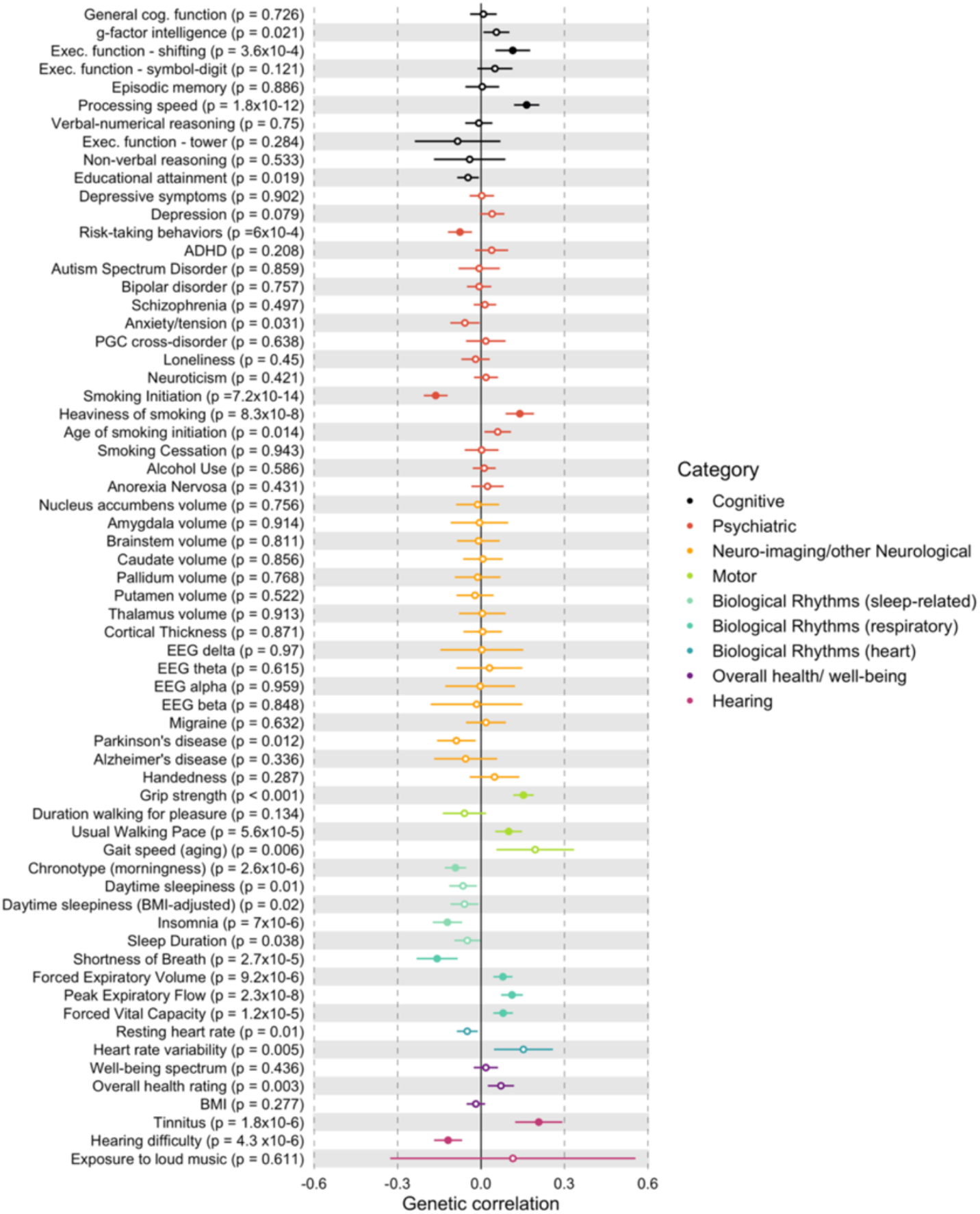
Cross-trait genetic correlations with beat synchronization. Results of exploratory genetic correlation analyses between beat synchronization and 64 traits from eight domains, conducted with LDSC. The x-axis is magnitude of genetic correlation (*rg*) with standard error visualized, and the (uncorrected) p-values for each trait’s correlation with beat synchronization are shown next to each trait label. Significant genetic correlations (after adjusting for multiple comparisons with a threshold of p<7.8×10^−4^) are shown with filled-in circles; empty circles are results that did not pass this threshold. See Supplementary Notes for detail on source studies. There are significant positive associations between beat synchronization and two of the cognitive domain GWASs; associations with smoking and risk-taking; two associations with hearing traits; two positive associations with motor function; and multiple associations with other biological rhythms (morning/evening chronotype, insomnia, and four breathing-related traits).

#### Genomic Structural Equation Modeling (SEM)

We conducted Genomic SEM^73^ to examine whether specific associations between beat synchronization and a subset of associated traits (e.g., musculoskeletal strength, walking pace, breathing function, and processing speed^74–76^) that are known to be related among each other in prior research^74–76^ represent distinct genetic relationships or share a common set of genetic influences with beat synchronization. The best fitting model, displayed in Figure 5, included a common genetic factor that accounted for genetic correlations among beat synchronization, grip strength, processing speed, usual walking pace, and expiratory flow. This common factor explained 11.6% of total variance in the beat synchronization GWAS and 9.6-25.0% of the variance in the other GWASs (see Supplementary Notes).

**Figure 5.**
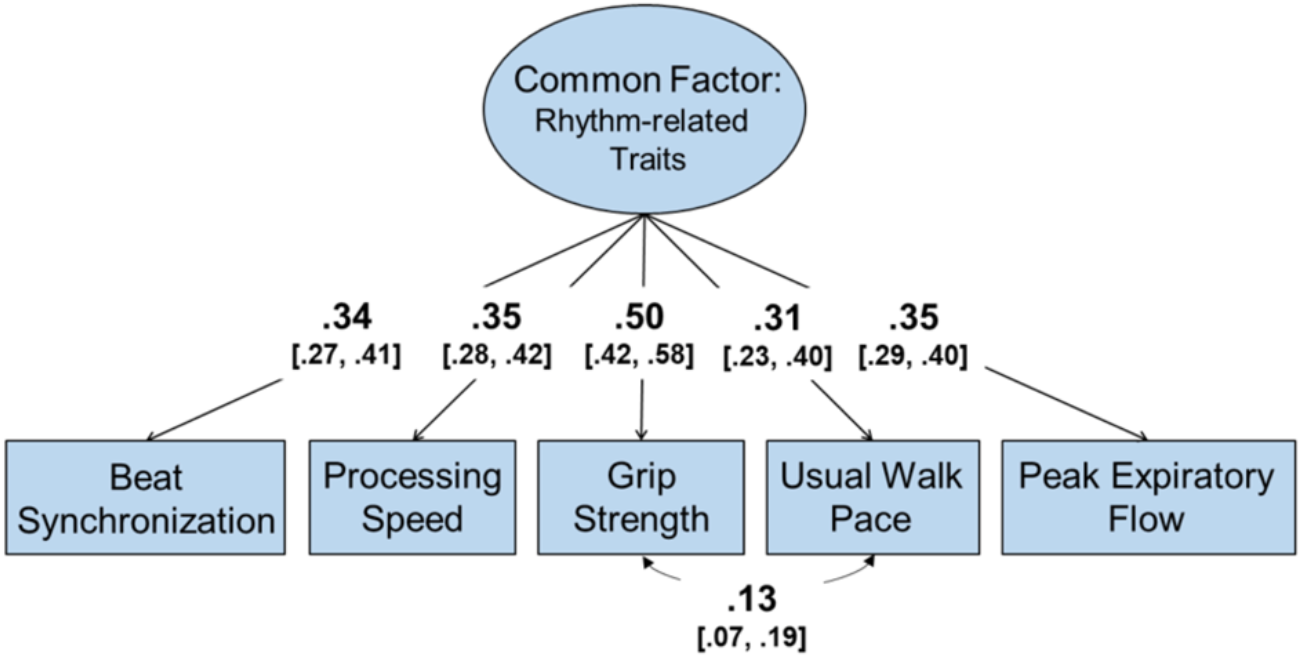
Genomic SEM model of beat synchronization and rhythm-related traits. The best-fitting genomic structural equation model of beat synchronization with GWASs of processing speed, grip strength, usual walking pace, and peak expiratory flow. 95% confidence intervals of factor loadings and correlations are displayed in brackets. Results suggest that beat synchronization was associated with the other traits through a set of common genetic influences. Model fit: *χ*^2^ (4) = 10.85, *p* = .028, CFI = .983, SRMR = .017.

#### Common Factor GWAS: Rhythm-Related Traits

Using genomic SEM, we conducted a multivariate GWAS (Supplementary Notes) on the latent genetic factor from the model presented above and portrayed in Figure 5. The heritability of this latent genetic factor was 7.27% (s.e.=0.25%) and there were 130 genome-wide significant loci (Supplementary Table 12; Supplementary Figure 8). Heritability was enriched for genes expressed in cerebellum (Supplementary Figure 9).

#### Cross-trait phenotypic extension of genetic correlations

Data from Phenotype Experiment 2 was analyzed to examine whether a subset of genetic correlations would be reflected in true phenotypic associations (pre-registered H4). Less accurate beat synchronization was weakly associated with a morningness preference (r=-.10, p=.015), more shortness of breath (r=-.16, p<.001), and smoking 20 or more (lifetime) cigarettes (r=-.11, p<.001) (Supplementary Table 13). In other words, accuracy in beat synchronization was correlated with eveningness chronotype, reduced shortness of breath when walking on level ground, and smoking abstinence (these associations go in the same direction as the genetic correlations; moreover, these associations remained significant after controlling for age, sex and education, and/or removing professional musicians from the sample.

#### Additional sensitivity analyses

Our results are robust to several potential biases (Supplementary Notes): the GWAS beat synchronization results are not driven by 1) shared genetic effects with general cognitive ability or educational attainment or 2) subtle residual population substructure, and 3) the *MAPT* association is not confounded with Parkinson’s Disease.

## Discussion

Our GWAS revealed highly polygenic architecture of the human capacity to synchronize to a musical beat, representing a significant advancement of our understanding of the genomic basis of a musicality phenotype. Heritability of beat synchronization is enriched for functions of the central nervous system on a number of dimensions: SNPs involved in neural development and brain-specific regulatory regions of the genome; genes involved in synaptic function; and gene expression in cortical and subcortical brain tissues aligned with auditory-motor regions previously linked beat perception and synchronization^34^.

Polygenic scores for beat synchronization were associated with self-identified musicians in a separate cohort, showing that the GWAS taps into the larger construct of musicality. Genetic correlations pointed to pleiotropy between beat synchronization and biological rhythms (including breathing function, walking pace, and chronotype), paving the way to a better understanding of the biological underpinnings of musicality and its health relevance.

In a series of phenotypic experiments, we also demonstrate that self-reported beat synchronization/rhythm measures can be used in large-scale population-based studies as suitable proxies for measuring individual differences in beat synchronization. Our findings indicate that the GWAS phenotype beat synchronization question was highly related to beat synchronization task performance (i.e., accuracy in tapping along to musical excerpts). Clearly the self-report is an imperfect correlate of beat synchronization; nevertheless, we demonstrate that it is a suitable proxy for very large-scale studies in which task administration is impractical. Furthermore, the GWAS phenotype is also significantly associated with: rhythm perception task performance^46^, a multi-item Rhythm questionnaire, and a well-established assessment of musical sophistication^43^. These results also converge with earlier work showing shared variance among task performance of beat synchronization, rhythm perception, and musical engagement/training^44,77–80^. The phenotypic associations were robust to demographic factors and self-confidence, and were not driven by the presence of professional musicians in the sample. These phenotype validation studies represent critical groundwork (see^81^) enabling brief rhythm self-report questionnaires to be deployed online in large-scale population genetic cohorts.

With sixty-nine loci surpassing the threshold for genome-wide significance, the polygenic architecture of the beat synchronization GWAS aligns with expectations for complex traits^82,83^. The top-associated locus mapped to *VRK2*, a gene previously associated with behavioral and psychiatric phenotypes (i.e. depression^84^, schizophrenia^85^ and developmental delay^86^), suggesting a biological connection between beat synchronization and neurodevelopment. The SNP-based heritability of beat synchronization on the liability scale was moderate, ranging from 13 to 16%, similar to heritability estimates of other complex traits (e.g., chronotype GWAS^87^) and consistent with moderate heritability estimates of musical rhythm abilities reported in twin studies^38–40^. Still, the limitation of the heritability adjustment on the liability scale is that the exact population prevalence of atypical beat synchronization is unknown and had to be estimated based on other indices of rhythm (see Supplementary Notes); this limitation should be addressed in future work.

We examined potential mechanisms linking genetic variation to neural architecture of the beat synchronization trait using multiple in-silico enrichment methods. Results showed enrichment of the heritability of beat synchronization in many brain tissues including cerebellum, dorso-lateral prefrontal cortex, inferior temporal lobe, and basal ganglia nuclei (i.e., putamen, caudate, and nucleus accumbens). This pattern of results likely reflects a genetic contribution to subcortical-cortical networks underlying musical rhythm perception and production^32,34^; furthermore, enrichment of brain-tissue-specific enhancers and active-regulatory regions in tandem with gene expression enrichments in brain tissue suggest that regions of the genome involved in regulation of gene expression within the beat perception and synchronization network contribute to phenotypic variance. Moreover, the partitioning heritability chromatin results showed an enrichment in both fetal and adult brain tissues, suggesting that beat synchronization may be the result of neurodevelopmental or basic brain processes. Gene set enrichments were also observed for synaptic function in the nervous system. Taken together, these results are a building block towards understanding how genes influence neural processes during beat perception and production, complementing results obtained with neuroimaging methods^88–93^.

Insights about the evolution of rhythm traits are suggested by the occurrence of two of the beat-synchronization-associated loci in human-accelerated regions (HARS) of the genome. In particular, rs1464791 is an expression quantitative trait locus (eQTL) that regulates expression of *GBE1* in multiple tissues, including muscle^59^; *GBE1* is also linked to neuromuscular disease^68^ and reaction time^69^. HARs are involved in many functions, so it is difficult to explicitly link their accelerated evolution to beat synchronization. It is too early to tell whether the overlap between beat synchronization-associated loci and those two HARS supports evolutionary theories about music (e.g., joint synchronous music-making has been posited to exert selective pressures in early humans by enhancing group social cohesion and family bonding^26,94^). The contribution of the genetic architecture of motor function to beat synchronization is further suggested by enriched heritability of SNPs that are enhancers located in musculoskeletal-tissue-specific regulatory regions of the genome, as well as our findings of genetic correlations between walking pace, musculoskeletal strength, and beat synchronization.

Moreover, our findings are promising for future large-scale genomic interrogations using comprehensive music phenotyping yielding continuous musicality variables (whether questionnaire-based^43,95^ or measured aptitude-based variables^38^), providing a path to examine potential genetic correlations between beat synchronization and other musical traits, such as music training or pitch discrimination, in line with family-based findings^36,37,41,96^. While the current data show a clear connection between the beat synchronization and broader musicality at the phenotypic and genetic levels, further genomic investigation in well-powered samples is needed to disentangle the *specificity* of genetic influences on beat synchronization from other genetic influences on musical traits, or motor or auditory function.

Finally, although our GWAS was based on self-report, the magnitude of the sample size bolstered statistical power. This is important because previous GWAS of other health traits based on self-report have effectively replicated associations from other GWAS of deeper phenotypes^83^, and it is generally acknowledged that powerful sample size can overcome some of the limitations arising from modest measurement error^97^.

Moving in synchrony to a musical beat encompasses beat perception and extraction, motor periodicity, meter perception, and auditory-motor entrainment (see ^4,32,98^ and Glossary in Supplementary Notes). Despite this complexity, beat is a highly frequent feature of many musical systems^1,3,26^. Indeed, we found that the heritability of beat synchronization is enriched in auditory-motor regions known to be active during rhythm tasks.^34^ It should be noted that beat perception and production does not depend on musical training or music genre, and atypical beat synchronization is not linked to lack of music exposure^99^. A limitation of the current work is the restriction of the genetic sample to a European ancestry (due to GWAS methodology constraints); investigating beat synchronization, musicality, and cross-trait correlations in populations of non-European ancestry should be a future priority for capturing the spectra of musicality traits in a wider range of ethnic, cultural and socio-economic contexts (see^100^). Regrettably, early research on individual differences in musicality in the early 1900’s was pursued not only using what we now recognize as highly culturally biased assessments, but also explicitly through the lens of eugenics (see^101^), similar to early research on individual differences in cognition. We strongly condemn the intent and design of those studies, and emphasize that the value of this work arises not from the hypothetical ranking of interindividual differences in beat synchronization (indeed, genomic associations with beat synchronization cannot be used to make deterministic predictions about individual abilities or aptitudes^102, 103^). Rather the value arises from discovering that the shared experience of rhythm, different though it is across cultures, is, in part, hardwired into our human genome. Furthermore, new knowledge on the genetic basis of musicality must be used ethically and fairly for research discovery and never for harm (e.g., discouraging a child from accessing musical activities).

We replicated previous findings implicating location 4q22.1 in musicality-related traits^36,55^ (*CCSER1* was the top-associated gene in our MAGMA analysis), but did not find support for previous gene associations from a set of genes that was drawn from prior candidate-gene, linkage, and GWAS studies with relatively small samples^54^. This is potentially due to well-known methodological problems with these methods particularly when applied to complex traits in small samples^104^. Without a second comparably sized GWAS available within which to conduct replication of the loci discovered in the primary GWAS, we were still able to demonstrate generalizability of these results by showing that PGS for beat synchronization predicts a musical trait in a separate biobank sample. The GWAS results of beat synchronization were nearly identical even after conditioning the results on GWASs of educational attainment and general cognition (g-factor); these results align with twin findings of specific genetic effects of rhythmic aptitude over and above any common genetic influences between rhythm and non-verbal cognitive variables^39,105^. Moreover, given both the likely capturing of genetic variation related to SES^106^ by the educational attainment GWAS summary stats, and the observation that our beat synchronization GWAS loci are robust to educational attainment, SES is unlikely to play a major role in our findings.

Our cross-trait explorations revealed pleiotropic effects between beat synchronization and several breathing-related phenotypes (peak expiratory flow, forced vital capacity, forced expiratory volume, and shortness of breath). We demonstrated phenotypically that more accurate beat synchronization task performance was related to lower likelihood of shortness of breath, mirroring the genetic correlations between beat synchronization and breathing function. In light of our genetic correlation between beat synchronization and three categories of traits (breathing, motor, and cognitive functions) previously shown to be genetically interrelated during the aging process^74,75^, we used genomic SEM to uncover shared genetic variance among beat synchronization and enhanced breathing function, greater grip strength, faster walking pace, and faster processing speed. Poor beat synchronization could be tied to certain health risks during aging, in light of other genetic and epidemiological work showing that lung function decline predicts later declines in motor function and psychomotor speed in older adults^107–110^.

The genetic correlation results suggest that beat synchronization shares common biology with a constellation of health traits, converging with the growing literature on the overlapping biomechanical and perceptual mechanisms of rhythms harnessed during synchronization, communication, muscle tensioning, and breathing; these relationships start very early in development^111,112^. The cerebellum in particular plays important roles in the control of coordinated movement, balance, respiration, dance, and even rhythm perception during passive listening to music^33^. Indeed, our rhythm-related traits multi-variate GWAS demonstrated enriched heritability of genes expressed in Cerebellar tissue, potentially of note in relation to experimental findings of functional synchronization of respiratory and upper limb movements during vocalization^5^. Moreover, “beat gestures” in speech involve the cerebellum^113^ and are inextricably linked to respiration, upper limb movement, and postural control, all of which may be biomechanically related to tapping or clapping to music.

Another dimension of biological chronometry was captured in the genetic correlation between chronotype and beat synchronization, which we replicated phenotypically (individuals who self-identified as ‘evening people’ tended to tap more accurately to music, even after removing professional musicians from the analysis). These results complement recent evidence of the increased prevalence of eveningness in musicians^114^, indicating that the relationship between chronotype and musicianship cannot solely be explained by environment (i.e., nocturnal job demands of professional musicians), but that also other shared biological factors may play a role. Given the genetic correlation between beat synchronization and lowered incidence of insomnia, the relationship between regulation of sleep, musicality, and rhythm represents an area for further exploration.

Our GWAS effectively identified alleles at 69 separate loci differentially associated with typical vs. atypical beat synchronization, complementing existing evidence of underlying neural mechanisms^77,79,80,99^. Future genetic studies could study beat synchronization as a continuous trait, either through self-report or internet-based task paradigms (i.e., REPP^47^). Prior literature on liability threshold models has shown that case-control GWAS of complex traits yield similar results to those obtained through continuous phenotypic measures (e.g., the genetic architecture of continuous measures of psychiatric symptoms is highly similar to the genetic architecture of cases versus controls^115^). Moreover, the use here of a population-based control group that is not “super-normal” increases the likelihood that the genetic correlations that we uncovered are reliable and not biased upward^116^.

Taken together, our results advance knowledge of the biological basis of beat synchronization by identifying genomic regions associated with individual differences in beat synchronization, estimating its cumulative SNP-based heritability, successfully applying a polygenic score model in a separate genetic sample, and exploring the enrichment of heritability in genes tied to central nervous system function. Movement in synchrony with a musical beat is a fundamental feature of music, and sensitivity to the beat emerges early in development, supporting childhood development in numerous ways^3,11,27,30^ and with importance over the lifespan^117^. We have elucidated the genetic architecture of beat synchronization and revealed its health relevance through cross-trait analyses. This study also provides a solid foundation for future exploration of how specific genetic variants contribute to neural mechanisms of entrainment, prediction, and reward harnessed during musical interactions^118^.

## Methods

### Phenotype validation studies

#### Phenotype Validation Experiment 1

##### Overview

Phenotype Validation Experiment 1 was designed to determine if self-reported rhythm abilities measured with the question used in the GWAS (i.e., ‘Can you clap in time with a musical beat?’) would be associated with task-based rhythm perception performance. The study was conducted in Amazon’s Mechanical Turk and received ethical approval from the Columbia University Institutional Review Board; participants gave their written informed consent, and the research complied with all relevant ethical regulations. We selected the Beat-based Advantage paradigm as a rhythm discrimination (perception) test due to its design of stimuli with simple and complex meter^119^ and prior history investigating individual differences in rhythm perception in a variety of brain and behavioural studies in adults and children with typical and atypical development^46,120–122^ as well as feasibility for internet-based adaptation. A questionnaire (self-report questions) was administered prior to the perception task, to avoid biasing participant self-report responses by how they perceived their own task performance. See Supplementary Notes for additional details on procedure, compensation, and self-report questionnaire.

##### Participants

The study sample was N=724 participants recruited anonymously in Amazon’s Mechanical Turk. All consented and passed a common headphone check^123^ that guarantees good listening conditions and the ability to follow basic instructions; this test also effectively filters out bots.

Participants (333 females; 387 males; 4 self-reported “other”) were 18-73 years old (mean = 36.1 years, SD=10.9) with 0-45 years of self-reported musical experience (mean 3.7 years, SD=5.7), representing an average degree of musical experience (see norms in^43^); demographics are reported in Table 1 (note that n=3 did not report their age).

##### Rhythm Perception Task

Stimuli for the rhythm perception task consisted of 32 rhythms drawn from prior work^46,119^. For each participant, we randomized with probability of one half the occurrence of “simple” rhythms (strong beat-based metrical structure and generally easier to discriminate) and “complex” rhythms (weaker metrical structure due to syncopation and generally more challenging to discriminate). Each rhythm was presented using pure tone stimuli in one of 6 frequencies (294, 353, 411, 470, 528, and 587 Hz, selected at random), and one of 4 durations (ISI of 220, 230, 240, and 250 ms). Each trial consisted of 3 rhythms separated by 1500 ms of silence; there were 32 trials of the task. The two first presentations were always identical, and in half of the trials (counterbalanced) the third rhythm was also identical (standard condition); in the other half of the trials, the rhythm differed by having one interval swapped (deviant condition). The pairings and structure of standard and deviant trials were taken from^46^. Participants were instructed that in each trial, they would listen to the series of three rhythms (the first two were always identical, and the third could be the same or different), and they had to indicate if the third rhythm was the same or different (see Supplementary Figure 2). Additional technical details are provided in the Supplementary Notes.

##### Data analysis

###### Self-report

Responses to the target question were as follows: n=654 (90.3%) participants answered ‘Yes’, n=25 (3.5%) answered ‘No’ and n=45 (6.2%) answered “I’m not sure.” Regarding an additional self-report question ‘Do you have a good sense of rhythm?’, n=503(67%) answered ‘Yes’, 102(14%) answered ‘No’ and n=117(16%) answered ‘I don’t know’. n=488 answered ‘Yes’ to both questions; the tetrachoric correlation between these two self-report questions was *r*=0.73.

###### Rhythm perception test

Responses to the rhythm perception test were analysed using signal detection theory^46,124^; this method is appropriate for discrimination tasks where the participant has to categorize stimuli along some dimension with the resulting *d’* values the strength of detection of the signal relative to noise. *d’* values were calculated on the 32 test trials. As expected from prior work^46,125^, individuals performed better at discriminating simple rhythms (mean d’= 1.98, SD =0.91) than complex rhythms (mean d’=1.43, SD =0.97) (t(724)=11.11, *p*<0.001, Cohen’s d=0.58).

To examine whether the target question was related to the objective (experimenter-measured) performance on the rhythm perception test, we performed a logistic regression analysis in which the clap-beat target question (Yes vs. No) was the outcome and quantitative scores on the rhythm perception test (*d’* scores) were the predictor. Covariates included age, education, and sex. McFadden’s R^2^ was also computed. We did not include ‘I’m not sure’ in the regressions, because this response was not available for data analysis in the GWAS. Given that the simple rhythms have a strong metrical structure that is known to facilitate detection and synchronization of the beat^46^, we also tested whether performance on the simple rhythm trials predicted self-reported beat synchronization (i.e., those who responded Yes to the clap-beat question). See Supplementary Notes for additional analyses.

#### Phenotype Experiment 2

##### Overview

The aims of Phenotype Experiment 2 were two-fold: 1) to validate self-reported beat synchronization phenotype as a proxy for objectively measured beat synchronization ability, and 2) to explore phenotypic associations between rhythm/beat synchronization and assorted traits found to be genetically correlated with beat synchronization. Phenotype Experiment 2 was pre-registered with Open Science framework (https://osf.io/exr2t) on July 8, 2020, prior to data collection. This internet-based study consisted of a beat synchronization task to assess the accuracy of participants’ tapping in time with musical excerpts, and a series of questionnaires assessing self-reported rhythm, musicality/music engagement, selected health traits, confidence as a personality trait, and demographics. We used REPP^47^ to measure participants’ tapping responses online with high temporal fidelity. The item from the GWAS study, “Can you clap in time with a musical beat?” with possible responses: Yes/No/I’m not sure, is referred to as the “target question.”

We tested the following hypotheses: *H1*: Self-report responses to the target question will be correlated with beat synchronization task performance (i.e., accuracy of tapping to the beat of music), such that individuals who respond Yes to the “target question” are predicted to tap more accurately to the beat of musical excerpts (i.e., they will have lower standard deviation of asynchrony than individuals who respond No to the target question). *H1*a: Self-report on a highly similar self-report question (“I can tap in time with a musical beat”) with responses on a 7-point agreement Likert scale are predicted to be correlated with tapping accuracy. *H2a:* The target question will be associated with broader rhythm ability/engagement (measured with a rhythm scale from seven other self-report questions). *H2b:* Beat synchronization task performance reflects broader self-reported rhythm ability/engagement. *H3:* To examine whether confidence (either as a personality trait or sureness in one’s own task performance) affects the reliability of self-reported beat synchronization. *H4:* Selected traits found to be genetically correlated with beat synchronization in the GWAS will be phenotypically correlated with beat synchronization task performance and the Rhythm Scale. Specifically: better beat/rhythm is correlated with evening chronotype (H4a), less shortness of breath (H4b), more tinnitus and loud music exposure (H4c), and more smoking (H4d); and that these associations would survive controlling for age, sex, and education (H4e). *H5*. Responses to the target question will be positively correlated with musical engagement measured with the Gold-MSI. *H6*. The associations in H4 would interact with being a musician, or more generally, with musical engagement.

##### Participants

A total of N=1,412 individuals met participation criteria outlined in the pre-registration (including passing the attention check item and not abandoning the study before completion). The study took place in Amazon Mechanical Turk and all participants provided informed consent in accordance with the Max Planck Society Ethics Council’s approved protocol; the research complied with all relevant ethical regulations. Participants (728 females; 678 males; 6 prefer not answer) were 18-77 years old (mean=36.3 years, SD=11.9) and had of 1-2 years of self-reported musical experience (Table 1). To ensure that the tapping technology measured beat synchronization with high temporal fidelity, it was crucial that participants complied with instructions to perform the tapping task (e.g., using the laptop speakers instead of headphones, with minimal background noise, etc.), and also used hardware and software without any technical issues that would preclude the recording signal (e.g., malfunctioning speakers or microphones, or the use of strong noise cancellation technology; see^47^). Thus, several precautions, including calibration tests and practice trials, were taken to make sure the tapping technology would work effectively, excluding cases that did not meet the requirements (see Supplementary Notes for details). A subset of n=542 had appropriate hardware to complete all parts of the study (including the tapping tests). Questionnaires were administered in the full sample of participants. Sample demographics are reported in Table 1. Demographics of the participants that completed the tapping experiment was highly similar to the full sample, as shown in the table; furthermore, 65.3% of the full sample and 64.9% of tapping sample had a Bachelor’s degree or higher.

#### Data collection for Phenotype Experiment 2

The first questionnaire included self-report items, including the “target question,” and also covering a variety of musical, health, and interest phenotypes. The health phenotype questions were chosen from phenotypes (chronotype, smoking, shortness of breath, and tinnitus) found to be genetically correlated with beat synchronization in our genetic analyses. Rhythm questions were selected for their particular relevance to various aspects of interacting/engaging with musical rhythm. The order of the questions was fixed for all participants. In addition, we used an attention check item^126^ between item 10 and 11, in order to exclude fraudulent responders, such as computer bots or disengaged participants responding randomly to the experiments. The end-questionnaire consisted of items covering the following additional self-report topics: another question about being a musician, a task confidence rating question, a Confidence scale, a 16-item short version of the Gold-MSI^43^ (items were chosen due to their high reliability scores: reliability omega = 0.92), and a Demographic questionnaire. Questionnaire items for Phenotype Experiment 2 are listed in the Appendix of the Supplementary Notes.

##### Tapping technology

Beat synchronization is particularly challenging to study with online research, where variability in participants’ hardware and software can introduce delay in latency and jitter into the recorded time stamps^127,128^. Here we used REPP (see^47^ for full details and a validation study of the technology), a robust cross-platform solution for measuring sensorimotor synchronization in online experiments that has high temporal fidelity and can work efficiently using hardware and software available to most participants online. To address core issues related to latency and jitter, REPP uses a free-field recording approach: specifically, the audio stimulus is played through the laptop speakers and the original signal is simultaneously recorded with participants’ tapping responses using the built-in microphone. The resulting recording is then analyzed using signal processing techniques to extract and align timing cues with high temporal accuracy.

##### Beat synchronization task

The beat synchronization task procedure consisted of three parts: calibration tests, practice phase, and main tapping phase. Participants started with the calibration tests, including a volume test to calibrate the volume of the laptop speakers to a level sufficient for detection by the microphone, a background noise test to make sure participants were in a quiet environment, and a tapping test to help participants practice how to tap on the surface of their laptop in the right level and location to be detected by the microphone. Participants were then presented with the practice phase, which consisted of four 15-second trials of isochronous tapping to a metronome beat (two with inter-onset interval of 500 msec and two with inter-onset interval of 600 msec). Following the practice phase, participants were presented with the main tapping task consisting of eight trials (4 musical excerpts, each played twice), with each trial 30 seconds long. The order of presentation of the practice trials and test trials was randomized for each participant.

The musical excerpts were drawn from the MIREX 2006 Audio Beat Tracking database in which musical excerpts had been annotated for beat locations by 30 listeners who tapped along to the music^129^. We chose these four MIREX clips that represent different music genres with different tempos and tapping difficulty: track 1 (“You’re the First, the Last, My Everything” by Barry White), track 3 (“El Contrapunto” by Los Mensajeros de La Libertad), track 7 (“Le Sacre du Printemps” by Stravinsky), and track 19 (“Possessed to Skate” by Suicidal Tendencies) of the MIREX training set (respectively). Based on the annotations in^129^, we identified the target beat locations from those consistently produced by the annotators. Additional technical details are provided in the Supplementary Notes, and Supplementary Figure 2 illustrates the instructions for participants.

#### Data Analysis

##### Beat synchronization task performance: Tapping accuracy analysis

Let S_t_ and R_t_ be the stimulus and response onsets, respectively. In case of the metronome S_t_ are the metronome onset (practice phase) and for music clips S_t_ is the target beat location based on the annotations. We define the asynchrony as a_t_=R_t_ -R_t_. Based on prior work^130^, we chose the standard deviation of the asynchrony (std(a_t_)) as our main target interest variable, as this appears to be a robust measure of individual performance and tightly linked to musical abilities^131^. We used metronome onsets to mark the beat metric level in an unambiguous way^132^. We emphasize that the metronome onsets were only physically present during the beginning and end of each clip. We used only the participant-produced asynchronies during the epoch at beats in which the guiding metronome was *not* present, in order to test the ability of the participants to synchronize to music without the metronome sounds (results were nearly identical when we included all onsets including the ones where physical metronome onsets were present). For the main test scores, we used the asynchronies computed relative to the virtual beat locations computed from prior human annotators in MIREX. We also computed vector length in order to confirm key associations of interest between the target question and beat synchronization accuracy (See Supplementary Notes).

##### Regression analyses

In accordance with the OSF preregistration, we examined whether responses to self-reported beat synchronization phenotype were associated with objectively-measured tapping accuracy, other self-reported measures of rhythm ability, confidence, and/or musical sophistication using logistic regression and McFadden’s R^2^ (for H1, H2a, H3, and H5) and linear regression (for H1a and H2b). Likewise, we used linear regression to examine potential replication of cross-trait associations uncovered by genetic analyses (H4a-d), to examine whether musical background interacted with the above associations (H6). Analyses were conducted in R version 3.5.1^133^. As described in our preregistration, individuals were recruited using MTurk and were included unless they failed and attention check item or abandoned the experiment before completing the study (N=1,412). Usable tapping data was available for n=542 individuals. The majority of exclusions were due to technical reasons detected by REPP’s signal processing pipeline during the practice trials (e.g., poor signal, noisy environment, wearing headphone, issues with laptop microphone, or people not tapping at all), but some additional subjects (n=19) were excluded for not having enough usable trials during data analysis. Missing covariates were handled using pair-wise deletion. Exclusion criteria are detailed in the Supplementary Notes.

##### GWAS of beat synchronization

Genome-wide association study summary statistics were generated from data acquired by personal genetics company 23andMe, Inc. Phenotypic status was based on responses to an English-language online questionnaire in which individuals self-reported “Yes” (cases) or “No” (controls) to the question ‘Can you clap in time with a musical beat?”. Individuals who responded “I’m not sure” were excluded from the genomic dataset as their data was not available. The GWAS included a total of 555,660 cases and 51,165 controls (total N=606,825, mean age(SD)=52.09(18.5), prevalence=92%), unrelated individuals of European ancestry; age range breakdown is provided in Table 1. All individuals provided informed consent according to 23andMe’s human subject protocol, which is reviewed and approved by Ethical & Independent Review Services, a private institutional review board (http://www.eandireview.com); the study complied with all relevant ethical regulations.

GWAS was conducted using logistic regression under an additive genetic model, while adjusting for age, sex, the top five principal components estimated from genetic data in order to control for population stratification, and indicators for genotype platforms to account for batch effects. We excluded SNPs with Minor Allele Frequency (MAF) <0.01, low imputation quality (R^2^<0.3) and indels, resulting in 8,288,850 SNPs in the GWAS summary statistics. Genotyping and QC details are provided in the Supplementary Notes.

##### Post GWAS enrichment analyses

###### FUMA-based analyses

The FUMA^58^ web application was used on the Genome-Wide Association summary statistics to identify genomic loci along with the “sentinel” SNPs that were independent in our analysis with a genome-wide significant P-value (<5 × 10^−8^) that are in approximate linkage disequilibrium (LD) with each other at R^2^<0.1 and to generate Manhattan plots and Quantile-Quantile plots. GWAS Catalogue associations for top loci were performed in FUMA (Supplementary Table 16).

Next, using the GWAS summary statistics as input for MAGMA (v1.08), we conducted a gene-based test of association, a gene property enrichment test, and a gene-set enrichment analysis. Gene property analysis^56^ utilized GTEx v8 data integrated in FUMA, with gene expression values log2 transformed average TPM per tissue type after winsorization at 50 based on GTEx RNA-seq data; this analysis was performed for 54 tissue types where the result of gene analysis was tested for one side while conditioning on average expression across all tissue types. We also performed exploratory GSA^57^ in FUMA using 15,556 Gene Ontology gene sets from the MsigDB database^134,135^; a Bonferroni threshold of 3.2×10^−6^ was used.

###### SNP-based heritability and partitioned heritability

SNP-heritability was computed with LD Score regression software^60^, and heritability estimates were adjusted to the liability scale based on population prevalence of atypical beat synchronization of 3.0%-6.5% (Supplementary Table 3, Supplementary Notes). We partitioned heritability of beat synchronization by 52 broad functional categories (Supplementary Table 7), using stratified LD score regression^60,63^ (Bonferroni-corrected significance level of p=9.6×10^−4^). We hypothesized that SNPs falling into open chromatin regulatory regions (i.e., accessible to transcriptional machinery), and regions with human-specific variation, would be enriched for beat synchronization-associated variation.

We further investigated (SNP-based) cell-type-specific and tissue-specific enrichments with LDSC-SEG (LDSC Specifically Expressed Genes)^67^, using a total of 697 gene sets (3 Cahoy gene sets, 205 Multi-tissue gene expression sets and 489 Multi-tissue chromatin sets from the RoadMap Epigenomics and ENCODE datasets); the Bonferroni-corrected significance level for this analysis was 7.1×10^−5^ (Supplementary Table 8). The X chromosome was not included in these analyses or any subsequent analyses using LDSC, given that the file that is required for LDSC analysis (w_hm3_snplist) does not include chromosome X SNPs.

###### Evolutionary analyses

The set of human accelerated regions (HARs) was taken from^65^. All variants in perfect LD (R^2^ = 1.0 in 1000 Genomes European participants) with variants in HARs were considered in the analysis. Similarly, variants tagging Neanderthal introgressed haplotypes were defined as in^136^. All variants in perfect LD with a Neanderthal tag SNP were considered Neanderthal variants. For each set, we performed stratified LDSC (v1.0.0) with European LD scores and the baseline LD-score annotations v2.1. The heritability enrichment is defined as the proportion of heritability explained by SNPs in the annotation divided by the proportion of SNPs in the annotation. Standard effect size (), which quantifies the effects unique to the annotation, is the proportionate change in per-SNP heritability associated with a one standard deviation increase in the value of the annotation, conditional on other annotations in the baseline v2.1 model^62^. To determine the expected number of overlaps between the N loci significantly associated with beat synchronization and HARs, we computed all overlaps between these sets of genomic regions (in hg19 coordinates) using bedtools2^137^. We then randomly shuffled the locations of HARs around the genome choosing segments of equal lengths and avoiding gaps in the genome assembly. We repeated this process 10,000 times and for each iteration computed the number of overlaps observed with the significantly associated loci. Based on this empirical distribution created with no association between the region sets, we computed the enrichment and p-value for the observed number of overlaps.

###### Genetic correlations

The genetic correlation method is designed to show whether there is shared genetic variation linked to a particular trait (here, our beat synchronization trait) and traits measured in other GWAS studies. We curated GWAS summary statistics for 64 complex traits representing a broad range of phenotypic categories: cognitive, psychiatric, neuro-imaging/other neurological, motor, other biological rhythms (circadian, heart, and breathing), overall health/well-being, and hearing (see Supplementary Table 10 and Supplementary Notes for details of the source studies). We estimated genetic correlations between beat synchronization and each of these traits using LDSC^72^, with a Bonferroni threshold of 7.5 × 10^−4^ (Supplementary Table 11).

###### Overview

We examined whether beat synchronization polygenic scores (PGS) would be associated with music engagement reported in health records. Individuals who disclosed music engagement to their care providers (which was subsequently recorded by their provider) were drawn from a recent phenome-wide study of 9,803 musicians^71^ identified from keyword searches of patient electronic health records (EHRs) in Vanderbilt University Medical Center’s de-identified research database (Synthetic Derivative). The phenotyping method was based on mining of clinical notes, utilizing 4 keywords and 449 regular expressions (i.e., “musician”, “plays the piano”); see Supplementary Notes and^71^ for details. The method was then validated with manually conducted chart review, with a positive predictive value (PPV) of 93%. Here we accessed the subset of n=1,259 musicians and 4,893 controls (matched for sex, median age (across the patients’ medical record), ethnicity, race, and length of record) that were also part of the BioVU database and had genotyped data on file, to test the hypothesis that higher PGS for beat synchronization would be associated with musical engagement operationalized as a having musician-related keywords/regular expressions recorded in an individual’s electronic health record.

We only selected individuals of European ancestry with genetic data that met standard quality control thresholds due to the poor performance of PGS trained in individuals of one ancestry and applied to individuals of another. This resulted in n=1,259 individuals (553 (44%) females, mean median age of record (SD)=53.1(16.5)) as musician “cases” and 4,893 controls (1,963(40%) females, mean median age of record (SD)=53.2(16.3)). See Supplementary Notes for details on the phenotyping, the samples, genotyping, and QC.

###### Polygenic scores

We used an IBD filter of 0.2 in order to include only unrelated European samples of BioVU. PGS were generated using the beat synchronization GWAS summary statistics, using software PRS_CS^138^. Briefly, this method uses a Bayesian regression framework and places continuous shrinkage (CS) prior on SNP effect sizes; this method outperforms previous methods in terms of prediction accuracy especially when the training sample size is large^138^, as is the case with the beat synchronization GWAS. The 1000genomes European reference set was used. The PGS was standardized to have a mean of 0 and SD of 1. Chromosome X was not included in the BioVU sample.

###### Data analysis

We conducted a logistic multivariable regression where the outcome variable was musician vs. control, the predictor variable was PGS for beat synchronization, and covariates included median age across the patients’ medical record, sex, top 10 principal components estimated from BioVU genetic data.

### Study FAQ

A live FAQ for the study is at: https://www.vumc.org/music-cognition-lab/news/faq-about-beat-synchronization-gwas-study

## Supporting information

Regional association plots

## Acknowledgements

We are grateful to 23andMe participants and employees for their contribution to the study, Nancy Cox and Simon Fisher for suggestions and insight throughout the process, Navya Thakkar for assistance with graphics, and Miriam Lense, Matthew D. Morrison, Aniruddh Patel, and Wim Pouw for thoughtful discussions. We would also like to thank Michaela Novakovic, Yune Lee, and Duane Watson for input during previous stages of the project, and Vanderbilt Trans-Institutional Programs for sparking the initial collaboration. RLG would like to thank Brandy Clark and P!nk for ongoing musical inspiration. Members of the 23andMe Research Team are: Michelle Agee, Stella Aslibekyan, Adam Auton, Robert K. Bell, Katarzyna Bryc, Sarah K. Clark, Sarah L. Elson, Kipper Fletez-Brant, Pierre Fontanillas, Nicholas A. Furlotte, Pooja M. Gandhi, Karl Heilbron, Barry Hicks, Karen E. Huber, Ethan M. Jewett, Yunxuan Jiang, Aaron Kleinman, Keng-Han Lin, Nadia K. Litterman, Jennifer C. McCreight, Matthew H. McIntyre, Kimberly F. McManus, Joanna L. Mountain, Sahar V. Mozaffari, Priyanka Nandakumar, Elizabeth S. Noblin, Carrie A.M. Northover, Jared O’Connell, Steven J. Pitts, G. David Poznik, Anjali J. Shastri, Janie F. Shelton, Suyash Shringarpure, Chao Tian, Joyce Y. Tung, Robert J. Tunney, Vladimir Vacic, and Xin Wang.

Research reported in this publication was supported by the National Institutes of Health Common Fund through the Office of the NIH Director and the National Institute on Deafness And Other Communication Disorders under awards DP2HD098859, K18DC017383, and R01DC016977. JAC was supported by the National Institutes of Health (R35GM127087). The content is solely the responsibility of the authors and does not necessarily represent the official views of the NIH. The dataset used for the analyses described were obtained from Vanderbilt University Medical Center’s BioVU which is supported by numerous sources: institutional funding, private agencies, and federal grants. These include the NIH funded Shared Instrumentation Grant S10RR025141; and CTSA grants UL1TR002243, UL1TR000445, and UL1RR024975. Genomic data are also supported by investigator-led projects that include U01HG004798, R01NS032830, RC2GM092618, P50GM115305, U01HG006378, U19HL065962, R01HD074711; and additional funding sources listed at https://victr.vumc.org/biovu-funding/. Also, The Genotype-Tissue Expression (GTEx) Project was supported by the Common Fund of the NIH, and by NCI, NHGRI, NHLBI, NIDA, NIMH, and NINDS. The funders had no role in study design, data collection and analysis, decision to publish or preparation of the manuscript.

## Author contributions

*Conceptualization of study:* RLG, LKD; *GWAS data acquisition and GWAS analyses: JFS*, 23andMe Research Team, DH, PS, MN; *GWAS and post-GWAS study design: LKD, RLG, JFS, MN, AJC, DH; Data visualization:* PS, MN, MAT, NJ, RLG, EE, DEG; *Post-GWAS analyses and interpretation: MN, DEG, PS, RLG, EE, EM, JAC, JFS, MM, FU, NC*; *Phenotype studies design: RLG, JDM, NJ, DEG, MAT; Phenotype studies data collection and analysis: MAT, NJ, DEG, EB, PS, MN, RLG, MM; Interpretation of results, writing, editing, and reviewing drafts:* All authors; *Project Supervision:* RLG, LKD, NJ, DH.

