## Supplementary notes for "Genome-wide association study of musical beat synchronization demonstrates high polygenicity"

### Glossary

**Music cognition terminology**

**beat perception:** perceptual inference of a pulse given a rhythmic pattern (source: Kotz et al, 2018 https://www.cell.com/trends/cognitive-sciences/fulltext/S1364-6613(18)30191-8)

**beat synchronization:** production of periodic motor actions synchronized to a perceptual beat, inferred from quasi-periodic auditory stimulus (source: Kotz et al, 2018).

**meter:** "hierarchical structuring of a series of events (which may or may not be strictly isochronous) into higher-order groupings." (source: Kotz et al, 2018)

**motor periodicity:** repetitive action with an identifiable frequency and phase; motor periodicity is "ubiquitous in biology, including heartbeat, breathing, running, swimming, chewing, wake/sleep cycle". (source: Kotz et al, 2018)

**musicality**: set of traits allowing for the perception and production of music, constrained by our cognitive and biological systems (Honing, 2018; https://nyaspubs.onlinelibrary.wiley.com/doi/full/10.1111/nyas.13638)

**rhythm discrimination task:** experimental paradigm to test perception/differentiation of musical rhythms; usually administered as a same-different task.

**rhythm:** "systematic pattern of events in terms of timing, accent, and grouping" (Patel 2008, Chap. 3 https://psycnet.apa.org/record/2008-04843-000

**Computational genetics terminology**

**chromosomal inversion:** change in orientation of a segment of DNA within a chromosome. source: Puig et al, 2015 https://www.ncbi.nlm.nih.gov/pmc/articles/PMC4576756/

**chronotype.** Individual preference for sleep patterns, i.e. behavioral manifestation of circadian rhythms resulting in morning types and evening types. (source: https://en.wikipedia.org/wiki/Chronotype)

**complex trait** "A trait that does not follow Mendelian Inheritance patterns, is likely derived from multiple genes, and exhibits a large variety of phenotypes. (source: https://www.nature.com/scitable/definition/complex-trait-82/ Nature Education)"

**electronic health record (EHR):** electronically stored and managed medical chart.

**gene regulation**: mechanisms that act to induce or repress the expression of a gene **(source:** https://www.nature.com/subjects/gene-regulation)

**Generalized Summary-data-based Mendelian Randomization (GSMR):** statistical method for testing for causal influence between phenotypes, using GWAS data. (Source: Zhu et al, 2018, Nature communications).

**Genetic correlations.** Genetic relationship between two traits, related to the concept of **pleiotropy** (a genetic locus that affects more than one trait). With LDscore regression software, it is possible to test genetic correlations between complex traits measured in separate samples. (sources: https://www.nature.com/articles/ng.3406 and https://www.nature.com/articles/s41576-019-0137-z)

**genome-wide association study (GWAS):** an approach that scans markers across genomes of many people to find common genetic variants associated with diseases or complex traits (source: https://www.genome.gov/about-genomics/fact-sheets/Genome-Wide-Association-Studies-Fact-Sheet)

**genomic locus:** location of a gene or DNA sequence. source: https://www.cancer.gov/publications/dictionaries/genetics-dictionary/def/locus and https://doi.org/10.1016/j.mehy.2011.01.019. In the present study, each locus is defined using FUMA's mapping of independent SNPs.

**handgrip strength.** A proxy for muscular fitness; handgrip strength is also predictive of other health and fitness traits (sources: https://www.nature.com/articles/ncomms16015 and https://www.nature.com/articles/s41598-018-24735-y)

**heritability on the liability scale:** method of adjusting heritability estimates in the research study sample to account for population prevalence of a given trait (source: https://www.cell.com/ajhg/fulltext/S0002-9297(11)00020-6)

**heritability**: the estimate of how much of the variation in a given trait can be attributed to genetic variation. source: https://ghr.nlm.nih.gov/primer/inheritance/heritability)

**Human Accelerated Regions (HARS):** "DNA sequences that changed very little throughout mammalian evolution, but then experienced a burst of changes in humans since

divergence from chimpanzees"
source: https://www.ncbi.nlm.nih.gov/pubmed/25156517

**linkage disequilibrium (LD): "**correlation between nearby variants such that the alleles at neighboring polymorphisms (observed on the same chromosome) are associated within a population more often than if they were unlinked."
source: https://www.sciencedirect.com/topics/neuroscience/linkage-disequilibrium

**Partitioned heritability: "**the proportion of genome-wide SNP heritability attributable to various functional categories" source: Finucane et al 2015 https://www.nature.com/articles/ng.3404

**polygenic risk scores** (also called polygenic scores or genetic risk scores or genetic risk profile scores): "The cumulative risk derived from aggregating contributions of the many DNA variants associated with a complex trait or disease" (source: (https://jamanetwork.com/journals/jama/fullarticle/2730627)

**single nucleotide polymorphism (SNP):** type of common genetic variation representing differences in building blocks of DNA (nucleotides). Source: https://ghr.nlm.nih.gov/primer/genomicresearch/snp

**SNP-based heritability:** total phenotypic variance explained by the aggregate of SNPs in a GWAS. source: https://www.nature.com/articles/ng.3941

**UK Biobank:** a large-scale community-based data repository, which houses many genetic, phenotypic, and other variables. https://www.ukbiobank.ac.uk/

### Additional details on Phenotype Validation Experiments

#### Additional Details of Phenotype Experiment 1

*Procedure.* This experiment was conducted in Amazon's Mechanical Turk (MTurk), an online service for crowd-sourcing workers for online tasks. Participants were invited to participate in an experiment where they would "listen to sounds and answer questions". All participants provided informed consent in accordance with the ethical protocol approved by the Columbia University IRB.

*Questionnaire.* To simulate the user environment within 23andMe where research participants answer a series of unrelated questions about health and other traits, we asked participants to provide answers for a series of randomly presented questions on a variety of other topics, such as "Do you have wisdom teeth?". Among these questions we embedded two rhythm-related questions: the target question: "Can you clap in time with a musical beat?" and an additional question that was intended to provide additional validation to the target question by using different wording, "Do you have a good sense of rhythm?". The questionnaire was administered first, to reduce the risk that another component of the session would prime participants or bias their responses. Participants then passed a headphone usage test[1] which guarantees good listening conditions as well as the ability to follow instructions. This test also filters bots, as it is unlikely that a bot could pass the headphone task without listening to the stimuli[1]. Participants that passed the headphone test were invited to perform the rhythm perception task.

*Rhythm perception test.* Participants received 8 training trials that were selected from rhythms that were not part of the test set, and then performed 32 rhythm perception task trials (see Methods for details of stimuli and task design and Supplementary Figure 2A for example). In all trials (practice and task) participants received feedback regarding their performance ("correct" and "incorrect"), with each correct trial resulted in a small monetary bonus (participants were paid about $1.60-$2.00 for participation). Participants who did not pass the headphone test received $0.20 for about one minute of answering the initial questions and performing the headphone test. Participant demographic data was collected after the rhythm test. The duration of the complete study was ~20 minutes.
Data analysis is described in the Methods section.

*Additional analyses for Phenotype Experiment 1 data.* Primary results are reported in the main manuscript. In addition, higher scores on the subset of "simple" (i.e., more strongly beat-based) rhythm trials were associated with Yes vs. No (OR(95%CI)=1.99[1.36-2.91], p<.001, McFadden's $R^2$ = .06. Similarly, higher scores on the subset of "complex" (syncopated, less strongly beat-based) rhythm trials were associated with Yes vs. No (OR(95%CI)=1.66 [1.10, 2.52], p=.017, McFadden's $R^2$ =.03.

#### Additional details of Phenotype Experiment 2

*Procedure and Questionnaire.* This experiment was also conducted in MTurk and was pre-registered with Open Science framework (Gordon et al.[2]; project at: https://osf.io/rp7bg/ ; pre-registration at: https://osf.io/exr2t). All participants provided informed consent in accordance with the Max Planck Society Ethics Council approved protocol. The complete experiment took 15-20 minutes. The experimental design took into account a number of considerations to ensure that the tapping technology would work efficiently, primarily: (a) we wanted to guarantee that the participant understood and complied with instructions to meet the necessary requirements for the tapping task (e.g., performing the task without headphones, minimal background noise, and calibrating the volume of the speakers to the right level); and (b) we wanted to make sure that the tapping technology allowed for reliable extraction of tapping onsets so that the measured variability in tapping response represents the participants' rhythmic ability, and not measurement noise.

There were five requirements for study participation: (i) participants must be at least 18 years old, (ii) participants must be fluent English speakers, (iii) participants must use a laptop to complete the full experiment (due to the technological constraints of the tapping tasks), (iv) participants must use an updated Google Chrome browser, and (iv) participants must be sitting in a quiet environment. In addition, as specified in the OSF pre-registration, to maximize reliability, we only recruited participants with a 95% or higher approval rate on previous tasks on MTurk. Participants were paid at a US $9/hour rate according to how much of the experiment they completed (with a base payment of $0.10 and a bonus of up to $2.87 upon completion). In those cases where participants terminated the experiment early (e.g., due to technical issues or failing the attention check), they were paid a proportional amount for the time spent on the experiment.

After providing informed consent, participants completed the first part of the study which consisted of 17 self-report items (fixed order) and the attention check, to make sure participants were reading the instructions and paying attention to the experiment (see Appendix for the full questionnaires. Participants that failed the attention check were excluded from the rest of the experiment, but they were paid proportionately for their time.

*Tapping Technology.* Measuring beat synchronization in online settings is challenging because it relies on experimental setups with high temporal fidelity to precisely align tapping onsets with the corresponding cue events. However, the high variability in participants' hardware and software typically observed in online experiments can introduce uncontrolled latency and jitter to the recorded time stamps[3,4]. To address this, we used REPP (Rhythm ExPeriment Platform), a recently developed technology for measuring sensorimotor synchronization in online experiments (see[5] for full details and a validation study of the technology). To address core issues related to latency and jitter, REPP uses a free-field recording approach: the audio stimulus is played through the laptop speakers and the original signal is simultaneously recorded with participants' tapping responses using the built-in microphone. Thus, by using a single audio recording to simultaneously capture the stimulus and tapping onsets, we can remove the most significant sources of delay in both response and presentation latencies. REPP then applies audio filtering and other signal processing techniques to the resulting audio recording to split the different components of the recording into separate channels and extract the stimulus and tapping onsets with reliable timing. Finally, REPP uses custom markers with known temporal locations to unambiguously identify the position of the tapping and stimulus onsets in the audio recording, allowing a precise alignment to measure participants' asynchronies.

*Beat Synchronization Task.* Before the tapping tasks, several precautions were taken to make sure that REPP was recording participants' tapping signals precisely. First, participants were instructed that the experiment can only be performed using the laptop speakers and they should unplug any headphones/ earplugs or disconnect any wireless devices. Second, participants were asked to calibrate the volume of their speakers to a level that was sufficiently good to be detected by the microphone. A sound meter was used to visually indicate when the level was appropriate. Third, the level of background noise was measured through the laptop's microphone, and visual and verbal feedback through the sound meter were provided to indicate whether the background noise was too noisy or just right. In cases where it was too noisy, participants were asked to either reduce the noise or move to a quiet environment. Finally, participants were asked to tap in the surface of their laptops with their index finger to test if the microphone could detect their signal; in cases where the signal was too low, participants were asked to tap in different locations of the laptop or try to tap louder. Screenshots of all instructions and feedback were included in the pre-registered protocol.

The practice phase consisted of four trials of isochronous tapping following a metronome sound (two with inter-onset interval of 500 msec and two with inter-onset interval of 600 msec), each 15 seconds long. In each trial, participants were asked to start tapping when the metronome started and stop tapping when the metronome stopped. After the first trial, the audio recording of the tapping was analysed using REPP's signal processing pipeline and feedback was provided based on the quality of the signal. If the signal was not sufficiently good to be detected by REPP, participants were reminded of the instructions outlined above and then were able continue with the practice phase. Moreover, after completing the four practice trials of the practice phase, the four corresponding audio recordings were analysed using the same procedure to determine whether the quality of the signal could be detected by REPP. Note that participants were never excluded based on their tapping accuracy but only based on technological constraints, i.e., whether the tapping signal was sufficiently good to be processed by the signal processing analysis performed by REPP or not. A threshold was used where participants with 3 or more usable practice trials were allowed to continue with the experiment, whereas participants with less than 3 usable trials were excluded from the tapping task but redirected to the final questionnaire portion of the experiment. Pilot testing indicated that about 30-50% of participants would have the appropriate technology to complete the tapping tasks. Our power estimates and the stopping rule in the OSF pre-registration took this constraint into account.

After the practice phase, participants were presented with the main tapping task. Participants were given the instructions on how to tap, and told that this time, they would tap to the beat of musical excerpts (see Supplementary Figure 2B). The tapping task consisted of 8 trials (4 musical excerpts of 30 seconds each, and each excerpt occurring twice). The order of trials was randomized for each participant. See Methods for description of the musical stimuli. To identify the beat of the musical excerpts in relation to annotations, we performed kernel density estimation with a kernel width of 20 msec; this provided an estimate of the probability of producing a response in any given time. The peaks of the probability density were located using Matlab's findpeaks function with the following parameters: 'MinPeakHeight', 0.11/ts, 'MinPeakProminence',0.11/ts, 'MinPeakDistance', 100 msec, where ts is the number of responses in the clip. Beat locations were extracted from the entire 30 seconds of the clip and used as the reference location for computing the asynchrony. To help participants find the beat and eliminate potential ambiguity of tapping at half- or double-time the tempo, a metronome marking the beats in the first 11 seconds of the clip were added to the stimulus (as commonly used in this type of tapping paradigm).

The final section of the experiment consisted of an additional self-report questionnaire (including two music-related questions about how the participant perceived their own task performance and their musical background; a Confidence scale[6]; a brief version of the Gold-MSI[7]; and a brief set of demographic questions on age, sex, and education). See Appendix for full questionnaire.

*Additional data analyses.*

*Accounting for self-confidence.* Results following the pre-registered analytic plan are reported in the main manuscript. In addition, we also tested *H3:* To examine whether confidence (either as a personality trait or sureness in one's own task performance) affects the reliability of self-reported beat synchronization (pre-registered H3). Results demonstrated that although responses to the target question were associated with confidence judgements of one's own tapping performance assessed either immediately after the tapping trials, OR = 1.72, p<0.026, 95% CI [1.05, 2.73], or confidence assessed as a personality trait, OR = 1.75, 95% CI [1.04, 3.06], p=0.041, controlling for these confidence measures had no credible impact on the association between the target item and tapping asynchrony, OR = 0.28, p<0.001, 95% CI [0.17, 0.43], McFadden $R^2$=.29 (H3). These findings suggest that while the target question may encompass some self-reporting bias, the bias does not diminish its strong association with true beat synchronization ability.

*Vector length of tapping as an outcome.* We also confirmed that key associations between self-report and the beat synchronization task were similar when using vector length of tapping to the musical excerpts as an outcome rather than SD of the asynchrony. This was done by computing the vector length to measure participant synchronization using circular statistics[8]. For each response $R_i$ we first identified the stimulus $S_j$ that immediately precedes the response $R_i$. We next computed the phase associated with the response $R_i$ using the following formula: $\boldsymbol{\phi_i = \frac{R_i}{S_{j+1} - S_j}}$

We then computed the average vector length defined as:

$$\boldsymbol{\bar{\phi} = |\Sigma_{i=1,\ldots,N} \exp(-2pi * \Phi_i)\,|/N}$$

Where we performed this computation on the complex plane.
This number lies between 0 and 1, where 1 indicates perfect synchrony and 0 indicates random phase.

Results show that individuals who respond Yes vs. No to the target question had longer tapping vector lengths, OR= 3.07 [2.10, 4.65], p < 0.001, McFadden's $R^2$ =0.21. Tapping accuracy with vector length was correlated with responses to a highly similar item ("I can tap in time to a musical beat") when asked on a seven-point Likert agreement scale, *r*=.37 [.30, .44], p<0.001. Vector length was positively associated with scores on the self-reported rhythm scale (responses to a seven-item questionnaire), *r*=.37, [.29, .44], p<0.001, and Musical Sophistication (Gold-MSI), *r* = .31 [.23, .38], p<0.001. Results are plotted in Supplementary Figure 3.

### Genotypes and QC of beat synchronization GWAS

The National Genetics Institute (NGI) performed the DNA extraction and genotyping on saliva samples for the 23andMe GWAS. Overall, there were five genotyping platforms and subjects were genotyped on only one of them. The v1 and v2 platforms had variants of the Illumina HumanHap550+ BeadChip, including approximately 25,000 custom SNPs selected by 23andMe, with a total of about 560,000 SNPs. The v3 platform had variants of the Illumina OmniExpress+ BeadChip, with custom content to improve the overlap with the v2 array, with a total of about 950,000 SNPs. The v4 platform covered about 570,000 SNPs, providing extra coverage of lower-frequency coding variation. The v5 platform, in current use, is based on an Illumina Infinium Global Screening Array (~640,000 SNPs) supplemented with ~50,000 SNPs of custom content. In cases where samples did not reach the 98.5% call rate, the sample was re-genotyped. When analyses failed repeatedly, then customers were re-contacted by 23andMe customer service to provide additional samples.

23andMe restricted participants to a set of unrelated individuals of European ancestry, determined through an analysis of local ancestry[9]. Relatedness was defined using a segmental identity-by-descent (IBD) estimation algorithm[10]. Imputation was conducted by combining the May 2015 release of 1000 Genomes Phase 3 haplotypes[11] with the UK10K imputation reference panel[12] to create a single unified imputation reference panel. Phasing was conducted using an internally-developed tool, Finch, which uses the Beagle graph-based haplotype phasing algorithm[13] for platforms V1 to V4 while for the V5 platform a similar approach was used with a new phasing algorithm, Eagle2[14]. SNPs with a Hardy-Weinberg $p<10^{-20}$, or a call rate of <90% were flagged. SNPs were also flagged if they were only genotyped on their 'V1' and/or 'V2' platforms due to small sample size and also if SNPs had genotype date effects. Finally, SNPs were also flagged if they had probes matching multiple genomic positions in the reference genome[10–14].

### Estimation of population prevalence of atypical beat synchronization.

In order to adjust SNP-based heritability on the liability scale, we estimated the population prevalence of atypical beat synchronization using three different sources of data available data for atypical rhythm. It is important to note that to our knowledge, there is not sufficient data in the literature from a large sample to estimate population prevalence of measured beat synchronization ability; thus instead, we used three different sources of data representing the prevalence of atypical rhythm in the population and to thus infer the prevalence of atypical rhythm.

First, out of a total sample of N=1,412 in our phenotype validation experiment 2 (see Methods and Results), n=43 answered 'No' to the question 'Can you clap in time with a musical beat?', yielding a prevalence of 3.04%.

Second, we consulted a large-scale study published by Peretz & Vuvan[15]. They report n=457 "time-based amusics" (i.e., they failed the Off-beat test but had normal performance on the pitch-based tasks) and n=569 individuals with an "uncategorized" musical deficit (for which failing the Off-beat Test was a criteria but they also had poor performance on at least one pitch-based task). Out of a total N=16,625, the combined n=1,026 with atypical rhythm yields a prevalence of 6.17%.

Third, we consulted data from N=6,881 Swedish twins who performed a rhythm aptitude test, reported in Ullén et al.[16]. The rhythm perception task was the Rhythm scale of the Swedish Music Discrimination Test. In each of 18 items, participants were instructed to indicate whether two consecutively presented rhythmic sequences are the same or different (binary forced choice). The binomial probability of performing significantly above chance on this test is a raw score of 12 items or more. The twin data showed that 6.44% of the study sample performed at chance or below, suggesting atypical rhythm ability.

Thus, considering evidence on atypical rhythm prevalence from these three data sources, we chose to use a range of values (3.0%, 3.5%, 4%, 4.5%, 5%, 5.5%, 6%, 6.5%) in our study for the heritability adjustment, because the prevalence of atypical beat synchronization in the GWAS sample appears to be slightly inflated/over-ascertained. Note that the differences in population prevalence do not impact the primary results from our GWAS or modify any of the post-GWAS analyses; the only impact this adjustment has on the results is for the liability adjustment of the total SNP-based heritability estimation (we also report the unadjusted heritability rate of 5%; see Results).

#### Genes previously implicated in musicality studies

Taking a list of 29 genes previously implicated in musicality studies[17] as well as *GATA2* and *PCDH7* from[18] and *UGT8* from[19], we examined our current data for replications of these associations by looking up the p-values of each gene from the MAGMA gene-based analysis (top associations are reported in Supplementary Table 4), as well as by including them in a gene-set and testing their associations en-masse. When examined independently, none of the genes reached the statistical significance threshold used for gene-based analysis ($p<2.56x10^{-6}$). Similarly, when examined as a gene set using Gene Set Analysis implemented in MAGMA, the association was not significant (b=0.11, s.e.=0.20, p=0.297). However, several of those genes are located nearby our top gene in the MAGMA analysis, *CCSER1* in the 4q22-24 region. The list of genes and their p-values in our analysis are reported in Supplementary Table 5.

#### Evolution of beat synchronization: Neanderthal Introgression Stratified Heritability Analyses

We evaluated the contribution of genetic variants detected in the Neanderthal genome present in modern Eurasians due to interbreeding (hereafter "Neanderthal variants") to the heritability of beat synchronization (see Methods). Eurasian genomes contain ~1.5-4% of DNA as a result from interbreeding with Neanderthals around 50,000 years ago. Heritability of beat synchronization was significantly depleted among Neanderthal variants (1.97-fold depletion, $p$ = 0.001). However, Neanderthal ancestry is significantly depleted in functional genomic regions overall[20]; therefore, the depletion of beat synchronization heritability in these regions is likely the result of the overall depletion for Neanderthal ancestry in functional regions of the genome. This is supported by a non-significant $\tau_c^*$, illustrating that Neanderthal vs. human variants do not provide unique heritability when conditioned on a broad set of regulatory elements (Supplementary Table 9).

#### Beat synchronization polygenic score approach: genetics of musicality in a health care context

We examined whether common alleles associated with rhythm en masse (also known as genetic profile risk scores or polygenic risk scores: PGS) predict musical engagement in a health care context. Musicians cases were drawn from a recently generated dataset[21] in which the authors accessed data from Electronic Health Records (EHRs) in the Vanderbilt University Medical Center's de-identified research database (Synthetic Derivative) and mined clinical notes for a collection of keywords and regular expressions that were indicative that the patient was a musician, utilizing 4 keywords and 449 regular expressions (e.g., "musician", "vocalist", "songwriter", "drummer", "plays the piano", "playing the guitar", "played violin", "player of the cello", "plays saxophone", "flutist", "plays the flute", "player of oboe", "accordion player", etc.). For more details on the phenotype parameters and methodology including chart review validation see: Niarchou et al.[21]. The full list of keywords and regular expressions are reported in Supplementary Table 2 of their study.

Controls were drawn from the control sample in Niarchou et al.[21] (matched for sex, median age (across the medical record), ethnicity, race, and length of record and also did not have any of the music-related keywords/regular expressions in their record. Although it is certainly possible that there are professional musicians within the Control group that do not have any indication of their musician profession associated with our particular keyword search in their EHR, the presence of such individuals within the control sample would only reduce power and increase the false negative rate of the analyses[22,23]. Our hypothesis was that higher PGS for beat synchronization would be associated with higher likelihood of having the musician-related keywords/regular expressions recorded in an individual's electronic health record.

Genotyping and QC of musicians and controls in the BioVU sample

The VUMC BioVU MEGA$^{EX}$ project genotyped ~100,000 samples over a period of 2.5 years. DNA was obtained from blood samples from the BioVU Biobank participants and were assayed using Illumina bead arrays (MEGA$^{EX}$) containing more than 2 million markers. The BioVU resource meets the criteria for non-human-subjects data, and IRB exemption was obtained to access this data.
We only selected individuals of European ancestry with genetic data that met standard quality control thresholds. Genotyped data was available from n=1,259 individuals (553 (44%) females, mean median age of record (SD)=53.1(16.5)) as musician "cases" that we compared with 4,893 controls (1,963(40%) females, mean median age of record (SD)=53.2(16.3)).

The pre-imputation procedures followed standard quality control procedures including SNP pre-cleaning, filtering at an individual call rate <0.98, clarifying sex discrepancies, |Fhet| >0.2. Data was imputed to the Haplotype Reference Consortium panel using the Michigan Imputation Server and converted from dosage to hard calls using PLINK's default settings. It was then filtered to include only biallelic SNPs, Minor Allele Frequency (MAF)<=0.005, $R^2$>=0.3 and call rates <0.98. SNPs were filtered for batch effects using logistic regression of paired imputation batches. SNPs were also filtered when MAFs within ancestry >0.1 difference from MAFs of corresponding 1000genomes MAFs. Finally, SNPs were also filtered when Hardy Weinberg Equilibrium p-value $<10^{-10}$ within ancestry.

#### Curation of GWAS summary statistics for genetic correlation analyses

We curated GWAS summary statistics for 64 complex traits representing a broad range of phenotypic domains, with details about the source studies in Supplementary Table 10.

*Cognitive* GWAS's included: general cognitive function from Davies et al.[24]; eight other cognitive traits from de la Fuente et al.[25] (g-factor, executive function- shifting; executive function – symbol digit; executive function – tower; episodic memory, processing speed, verbal-numerical reasoning), and educational attainment from Lee et al.[26].

*Psychiatric* GWAS's included: Depressive symptoms from Baselmans et al.[27]; Depression from Howard et al.[28]; Risk-taking behavior from Karlsson Linner et al.[29]; Attention Deficit Hyperactivity Disorder from Demontis et al.[30]; Autism Spectrum Disorder from Grove et al.[31]; Bipolar Disorder from Mullins et al.[32]; Schizophrenia from Pardinas et al.[33]; Anxiety from Hill et al.[34]; PGC Cross-Disorder from Lee et al.[35]; Loneliness from Day et al.[36]; Neuroticism from Baselmans et al.[27]; four Smoking-related traits and Alcohol Use from Liu et al.[37]; and Anorexia Nervosa from Watson et al[38].

*Neuro-imaging/other neurological* GWAS's included: seven subcortical volume traits from Satizabal et al.[39]; cortical thickness from Grasby et al.[40]; four EEG traits from Smit et al.[41] (delta power, theta power, alpha power, and beta power); migraine from Watanabe et al.[42]; Parkinson's Disease from Nalls et al.[43]; Alzheimer's Disease from Kunkle et al.[44]; and handedness from Watanabe et al.[45].

*Motor function* GWAS's included: Grip strength (right hand) from Zhu et al./LDHub[46], duration of walking for pleasure and usual walking pace from Watanabe et al.[42]; and gait speed in old age from Avraham et al.[47].

GWAS's of *Other biological rhythms* included: circadian/sleep-related traits including chronotype (morningness) from Jones et al.[48]; daytime sleepiness (adjusted and non-adjusted for Body Mass index; BMI) from Wang et al.[49]; insomnia from Jansen et al.[50]; and sleep duration from Dashti et al.[51]. Three breathing function traits were used from Shrine et al.[52] (forced expiratory volume, peak expiratory flow, and forced vital capacity), in addition to shortness of breath from Watanabe et al.[42]. Two heart-rhythm traits were resting heart rate from Zhu et al.[53] and heart rate variability from Nolte et al.[54]

*Hearing-related* GWAS's included hearing difficulty from Wells et al.[55], and tinnitus and exposure to loud music from Zhu et al./LDHub[46].

GWAS's of Overall health/well-being included well-being spectrum from Baselmans et al.[27]; overall health rating from Watanabe et al.[42]; and BMI from Pulit et al.[56].

#### Genomic Structural Equation Modeling

Analyses were conducted using the genomic structural equation modeling (SEM) R package[57], an extension of LD score regression[58] which calculates genetic correlations between any two traits with summary statistics available, provided the samples were drawn from the same ancestral background. Using LD score regression, genomic SEM computes a full genetic correlation matrix across the set of traits for which GWAS summary statistics are provided, and then estimates the model using this correlation matrix using the lavaan package[59] in R. Model fit was determined based on chi-square tests ($\chi^2$), the Comparative Fit Index (CFI), and the standardized root mean squared residual (SRMR), with good fitting models expected to have CFI > .95 and SRMR < .08. Good-fitting models also traditionally have nonsignificant $\chi^2$ statistics, but because GWAS sample sizes are extremely large, and this statistic is sensitive to sample size, we focused on other fit indices. Significance of individual parameter estimates were established with 95% confidence intervals (95% CIs).

Aside from beat synchronization, Grip strength (right hand) GWAS were from GWAS Analyses from UK Biobank (housed at http://www.nealelab.is/uk-biobank)[46]; Usual Walking pace GWAS was from[42], and Peak Expiratory Flow was from[52]. Additionally, processing speed summary statistics were from[25] de la Fuente et al. (2021), but only for the initial SEM model (not enough information was provided for the subsequent multivariate GWAS, so we used the UK Biobank GWAS version for multivariate GWAS. The two versions of this GWAS were genetically correlated at 1.0, suggesting they are nearly identical). SNPs on X chromosome were included in where available (i.e., for beat synchronization, processing speed, and grip strength only). Additional analyses evaluated whether smoking initiation GWAS (based on[37]) could also be included in this common factor. However, this model was not supported, likely because smoking initiation was genetically correlated with beat synchronization ($r_g$= -.16) and walking pace ($r_g$= .21) in opposite directions, and was uncorrelated with the other traits ($r_g$s = -.07 to .01).

The initial common factor model of beat synchronization, grip strength, processing speed, walking pace, and expiratory flow fit the data well, $\chi^2$ (5) = 23.93, $p$ < .001, CFI = .954, SRMR = .025. However, we noticed that walking pace and grip strength appeared more correlated with one another than any of the other genetic correlations ($r$=.28), and model fit was improved by including a residual correlation between these two GWAS, $\chi^2$ (4) = 10.85, $p$ = .028, CFI = .983, SRMR = .017. No additional adjustments were made to this final model, displayed in Figure 5 of the main manuscript.

Next, using genomic SEM, we also conducted a multivariate GWAS on the latent genetic factor, which we termed *rhythm-related traits*. Individual SNP effects were estimated for the common factor if they were available in all summary statistics files, had a minor allele frequency > 1.0%, and were present in the 1000 Genomes Phase 3 v5 reference panel (N=6,546,496 SNPs). We then used FUMA v1.3.6 to identify independent loci and estimate gene expression using the same parameters as in the primary GWAS (see Supplementary Figures 8 and 9). Genome-wide significant loci are reported in Supplementary Table 12.

### Cross-trait phenotypic correlations

Analyses of data from phenotype experiment 2 revealed that some of the cross-trait phenotype relations mirrored the genetic correlations, while others did not (Supplementary Table 13). Primary results are reported in the main manuscript. In addition, individuals who had better beat synchronization were less likely to report ever smoking, and less likely to report tinnitus (these associations were opposite of what was found in the genetic study); moreover, the association with loud music exposure was nonsignificant. These associations with chronotype, shortness of breath, and smoking remained significant after controlling for age, sex and education, and/or removing professional musicians from the sample. Self-reported rhythm (assessed using the seven-item Rhythm scale) was only associated with smoking status ($r$=-.08) and loud music exposure ($r$=-.13), even when controlling for covariates or focusing on non-musicians; however, these associations appeared in the opposite direction of the corresponding genetic associations. There was no evidence of interactions with musical sophistication or prior/current musician status for the H4 constructs, except that the association between loud music exposure and self-reported rhythm was weaker in individuals with who more actively performed music ($p$=.022), though this effect would not survive a strict multiple test correction.

### Beat synchronization beyond the contribution of cognition and educational attainment

In light of previous work linking rhythm and IQ[60,61], we used multi-trait conditional joint analysis[62] (mtcojo v.1.91.7 beta) to remove potentially shared genetic effects between beat synchronization and general cognition, and also between beat synchronization and educational attainment. These analyses generated two new sets of summary statistics of beat synchronization in which the betas, standard errors and p-values were adjusted based on the summary statistics of the GWAS of general cognition and then separately, on the GWAS of educational attainment.

*Conditioning based on general cognition.* We selected the *g*-factor from de la Fuente et al.[25], with N= 331,000 for general cognition. In their study, a *g*-factor was obtained from the following tests: reaction time, matrix pattern recognition, verbal numerical reasoning, symbol digit substitution, memory pairs-matching test, tower rearranging, trail making test-B. Higher scores indicate more optimal performance. Only autosomal chromosomes were tested, as the publicly available data did not contain X chromosome SNPs. Default parameters were used in the mtcojo software, and the Mtcojo output constituted conditioned beat synchronization summary statistics, upon which we used FUMA as described in the Methods section for the primary GWAS. FUMA identified 62 genome-wide significant loci (with 63 sentinel SNPs) which were very similar to the 67 autosomal loci of the original beat GWAS, albeit some slight attenuation of p-values. Each of these 62 loci was within 5kb of the loci of the unadjusted beat synchronization GWAS. The adjusted genome-wide significant loci are reported in Supplementary Table 14, and Supplementary Figure 10 displays the Miami plot of the unadjusted and adjusted results.

*Conditioning based on educational attainment*. GWAS summary statistics for educational attainment from Lee et al.[26], N= 766,345, were used for a similar analysis. FUMA results of the beat GWAS summary statistics adjusted for educational attainment demonstrated 65 genome-wide significant autosomal loci (Supplementary Table 15 and Supplementary Figure 10; there were 68 sentinel SNPs), also highly similar to loci of the unadjusted summary statistics (within 5kb of the loci of the unadjusted GWAS). Only autosomal chromosomes were tested, as the publicly available data did not contain X chromosome SNPs.

In addition, the estimates of SNP-based heritability (5%) and SNP-based heritability on the liability scale (13% to 16%) remained similar in both sets of adjusted summary statistics (Supplementary Table 3). Taken together, the conditional analyses suggest that the genetic architecture of beat synchronization is largely not confounded by general cognitive skills or educational attainment.

### Population substructure adjustment

Recent studies illustrate the potential for very subtle residual population substructure to influence some polygenic analyses[63] including genetic correlations. Therefore, we also adjusted the beat synchronization associations for SNP-loadings on the first principal component of ancestry estimated from 1KG European populations. Data was accessed from a recent study[64] in which PC loadings had been generated from the 1000 Genomes Project[11] because individual-level genotype data was unavailable on the analysed sample, following[63]. Specifically, the authors[64] selected all unrelated subjects from 1000 Genomes Phase 3 data. After applying a number of excluding criteria (e.g., removing SNPs with MAF<-5%). 264,339 SNPs remained and PCA was performed. Further, the PC scores were regressed on the genotype allele count of each SNP (adjusting for sex), and the regression coefficients were used as SNP PC loading estimates, following[65]. For the first 20 PCs, each PC weight was tested for association with each subject's genotype in PLINK, and the degree of association of that SNP to population frequency differences was identified (Beta_PCs). Only autosomal chromosomes were tested, as the data generated from the 1000 Genomes Project did not contain X chromosome SNPs.

We then merged the beat-synchronization GWAS summary statistics with the Beta_PC values, and applied an ancestry regression analyses following[63], which provided us with ancestry-corrected effect sizes. We then used these SNP estimates of ancestry to adjust the beat synchronization GWAS results, upon which we recalculated the SNP-based heritability, lambda GC intercept, ratio, and calculated the heritability estimates in the liability scale (using LDSC software); results of these analyses are presented in Supplementary Table 3, and show no notable changes from the original GWAS. We also used FUMA (with same parameters as in the primary analysis) to estimate independent sentinel SNPs and genomic loci on the ancestry-adjusted GWAS (Supplementary Table 17), which were highly similar to the loci in the unadjusted GWAS.

### Sensitivity analyses of ‘clap to beat’ phenotype and Parkinson’s Disease

One of the associated loci in the GWAS study is the *MAPT* locus (17q21 locus), known to be associated with Parkinson’s disease (PD)[66]; in particular, our independent SNP in the locus, rs4792891, is in mild LD with the independent SNP associated with PD (rs365825; r2=0.55). In light of prior research in PD patients showing lower rhythm perception task performance[67], they may have also have difficulty clapping in time with a musical beat. Here we found that PD status is significantly associated with difficulty in clapping to the beat (OR=0.5996, p=7.93e-35, adjusting for age, sex, 5 PCs). Due to the 23andMe-Michael J Fox Foundation collaboration, PD patients are over-represented in the 23andMe database. We tested for the possibility that the presence of PD patients in our study sample (less than 1% of the total) could account for the *MAPT* associations by removing PD cases (N=5,644) and fitting the same association model adjusting for age, sex, five principal components, and genotyping platforms. The result shows the variant rs4792891 is still associated with the clap-to-beat phenotype ($p=1.61x10^{-13}$), thus showing that the *MAPT* association with the clap-to-beat phenotype is not driven by the presence of PD cases in our sample (Supplementary Table 18).

### Appendix. Self-report questionnaires used in Phenotype validation experiment #2.

**Part I: self-report questionnaire**

1. Do you consider yourself to be:
    - Definitely a “morning” person
    - More a “morning” than “evening” person
    - Neither a “morning” person or an “evening” person
    - More an “evening” than a “morning” person
    - Definitely an “evening” person
2. [Collected for a separate study] In general, how satisfied are you with your friendships?
    - Extremely happy
    - Moderately happy
    - Moderately unhappy
    - Very unhappy
    - Extremely unhappy
    - Do not know
    - Prefer not to answer
3. Can you clap in time with a musical beat?
    - Yes
    - No
    - I’m not sure
4. Do you get short of breath when walking with people of your own age on level ground?
    - Yes
    - No
    - Do not know
5. I have smoked 20 or more cigarettes in my lifetime (include cigars, pipe tobacco, and chewing tobacco).
    - Yes
    - No
6. How many days have you smoked in the past 180 days?
    - Open text
7. Do you get, or have you had, noises (such as ringing or buzzing) in your head, or in one or both ears, that lasts for more than five minutes at a time?
    - Yes, now most or all of the time.
    - Yes, now a lot of the time.
    - Yes, now some of the time.
    - Yes, but not now, but have in the past.
    - No, never
    - I don’t know
8. Have you ever listened to music for more than 3 hours per week at a volume which you would need to shout to be heard, or, if wearing headphones, someone else would need to shout for you to hear them?
    - Yes, for more than 5 years
    - Yes, for around 1-5 years
    - Yes, for less than a year
    - No
    - I don’t know
9. [Collected for a separate study] Have you ever been diagnosed with dyslexia?
    - Yes

- No
- I don’t know

10. [Collected for a separate study] Did you get speech-language therapy as a child?
    - yes
    - No
    - I don’t know

11. I can tell when people sing or play out of time with the beat.
    - 1. Completely disagree
    - 2. Strongly disagree
    - 3. Disagree
    - 4. Neither agree nor disagree
    - 5. Agree
    - 6. Strongly agree
    - 7. Completely agree
12. I can tap in time with a musical beat.
    - 1. Completely disagree
    - 2. Strongly disagree
    - 3. Disagree
    - 4. Neither agree nor disagree
    - 5. Agree
    - 6. Strongly agree
    - 7. Completely agree
13. My rhythmic ability is important to my identity.
    - 1. Completely disagree
    - 2. Strongly disagree
    - 3. Disagree
    - 4. Neither agree nor disagree
    - 5. Agree
    - 6. Strongly agree
    - 7. Completely agree
14. I struggle to feel the rhythm when listening to, playing, or dancing with music.
    - 1. Completely disagree
    - 2. Strongly disagree
    - 3. Disagree
    - 4. Neither agree nor disagree
    - 5. Agree
    - 6. Strongly agree
    - 7. Completely agree
15. When I hear a tune that I like a lot, I can't help tapping or moving to its beat.
    - 1. Completely disagree
    - 2. Strongly disagree
    - 3. Disagree
    - 4. Neither agree nor disagree
    - 5. Agree
    - 6. Strongly agree
    - 7. Completely agree
16. My musical ability is important to my identity.
    - 1. Completely disagree
    - 2. Strongly disagree
    - 3. Disagree
    - 4. Neither agree nor disagree

- 5. Agree
- 6. Strongly agree
- 7. Completely agree

17. I am confident that I can clap accurately in time to a musical beat.
    - 1. Completely disagree
    - 2. Strongly disagree
    - 3. Disagree
    - 4. Neither agree nor disagree
    - 5. Agree
    - 6. Strongly agree
    - 7. Completely agree

**Sources of questions drawn from extant questionnaires:**

*Q1, Q2, Q4, Q5, Q6, Q7, Q8: UKBioBank. http://biobank.ndph.ox.ac.uk/showcase/*

*Q15. Barcelona Music Reward Questionnaire. Mas-Herrero, E., Marco-Pallares, J., Lorenzo-Seva, U., Zatorre, R. J., & Rodriguez-Fornells, A. (2013). Barcelona Music Reward Questionnaire (BMRQ) [Database record]. APA PsycTests. https://doi.org/10.1037/t31533-000*

*Q11. Goldsmith Musical Sophistication Index. Müllensiefen, D., Gingras, B., Musil, J., & Stewart, L. (2014). The musicality of non-musicians: an index for assessing musical sophistication in the general population. PloS one, 9(2), e89642.*

*Q19. (see Part III below; this item is from an Adaptation of Creative Achievement in Music Questionnaire). Mosing, M. A., Verweij, K. J., Abé, C., de Manzano, Ö., & Ullén, F. (2016). On the relationship between domain-specific creative achievement and sexual orientation in Swedish twins. Archives of sexual behavior, 45(7), 1799-1806.*

**Part II. Beat synchronization task.**

See Anglada Tort et al., 2021[5] for visual examples of the utilization of the REPP method.

**Part III: post-test questionnaires**

**Post-test confidence**

Q18. I am confident that I tapped accurately in time to the musical beat in the excerpts.

- 1. Completely disagree
- 2. Strongly disagree
- 3. Disagree
- 4. Neither agree nor disagree
- 5. Agree
- 6. Strongly agree
- 7. Completely agree

**Short version of Gold-MSI**

| ID* | Item | Response |
|---|---|---|
| AE_01 | I spend a lot of my free time doing music-related activities. | Agreement scale |
| AE_02 | I enjoy writing about music, for example on blogs and forums. | Agreement scale |
| EM_04 | I am able to identify what is special about a given musical piece. | Agreement scale |
| MT_02 | At the peak of my interest, I practised my primary instrument for _ hours per day. | 0 / 0.5 / 1 / 1.5 / 2 / 3-4 / 5 or more |
| MT_03 | I have never been complimented for my talents as a musical performer. | Agreement scale |
| MT_06 | I can play _ musical instruments. | 0 / 1 / 2 / 3 / 4 / 5 / 6 or more |
| MT_07 | I would not consider myself a musician. | Agreement scale |
| PA_04 | I can compare and discuss differences between two performances or versions of the same piece of music. | Agreement scale |
| PA_08 | When I sing, I have no idea whether I'm in tune or not. | Agreement scale |
| SA_01 | If somebody starts singing a song I don't know, I can usually join in. | Agreement scale |
| SA_02 | I can sing or play music from memory. | Agreement scale |
| SA_03 | I am able to hit the right notes when I sing along with a recording. | Agreement scale |
| SA_04 | I am not able to sing in harmony when somebody is singing a familiar tune. | Agreement scale |
| SA_05 | I don't like singing in public because I'm afraid that I would sing wrong notes. | Agreement scale |
| SA_06 | After hearing a new song two or three times, I can usually sing it by myself. | Agreement scale |
| MT_05 | I have had _________ years of formal training on a musical instrument (including voice) during my lifetime. | 0/ 0.5/ 1/ 2/ 3/ 4-6 / 7 or more years |

**Information about the questionnaire (Gold-msi):*

Müllensiefen, D., Gingras, B., Musil, J., & Stewart, L. (2014). The musicality of non-musicians: an index for assessing musical sophistication in the general population. *PloS one*, *9*(2), e89642.

Metrics of the 15 items selected:

| Item N | IRT reliability | IRT error | Reliability alpha | Reliability omega | Validity MT | Validity BT |
|---|---|---|---|---|---|---|
| 15 | 0.93 | 0.26 | 0.92 | 0.92 | 0.28 | 0.37 |

**Musicianship**

Q19. How engaged with music are you? Singing, playing, and even writing music counts here. Please choose the answer which describes you best:

- I am not engaged in music at all
- I am self-taught and play music privately, but I have never played, sung or shown my music to others
- I have taken lessons in music, but I have never played, sung or shown my music to others
- I have played or sung, or my music has been played in public concerts in my home town, but I have not been paid for this
- I have played or sung, or my music has been played in public concerts in my home town, and I have been paid for this
- I am professionally active as a musician
- I am professionally active as a musician and have been reviewed/featured in the national or international media and/or have received an award for my musical activities

**Confidence Scale (PEI)**

Questions*:

1. I often feel unsure of myself even in situations I have successfully dealt with in the past.
2. I lack some important capabilities that may keep me from being successful.
3. Much of the time I don't feel as competent as many of the people around me.
4. I have fewer doubts about my abilities than most people.
5. When things are going poorly, I am usually confident that I can successfully deal with them.
6. I have more confidence in myself than most people I know.
7. If I were more confident about myself, my life would be better.

Choices: Strongly Agree/ Mainly Agree/ Mainly Disagree/ Strongly Disagree

*This is the general subscale, assessing one's confidence to perform competently in general.
Shrauger, J. S., & Schohn, M. (1995). Self-confidence in college students: Conceptualization, measurement, and behavioral implications. *Assessment*, *2*(3), 255-278.

More information: Stankov, L., Kleitman, S., & Jackson, S. A. (2015). Measures of the trait of confidence. In *Measures of personality and social psychological constructs* (pp. 158-189). Academic Press.

**Demographics**

Sex: Please indicate your sex.

- Male
- Female
- Prefer not to answer

Age: Please indicate your Age in years.

- Input text

Education: Please indicate the highest level of schooling you have completed:

- Less than high school education
- High school (or equivalent)
- Some college or Associate's degree
- Bachelor's or equivalent degree
- Master's or equivalent degree
- Doctorate or equivalent degree

### Supplementary Figures

##### Supplementary Figure 1. Study design and Analyses pipeline.

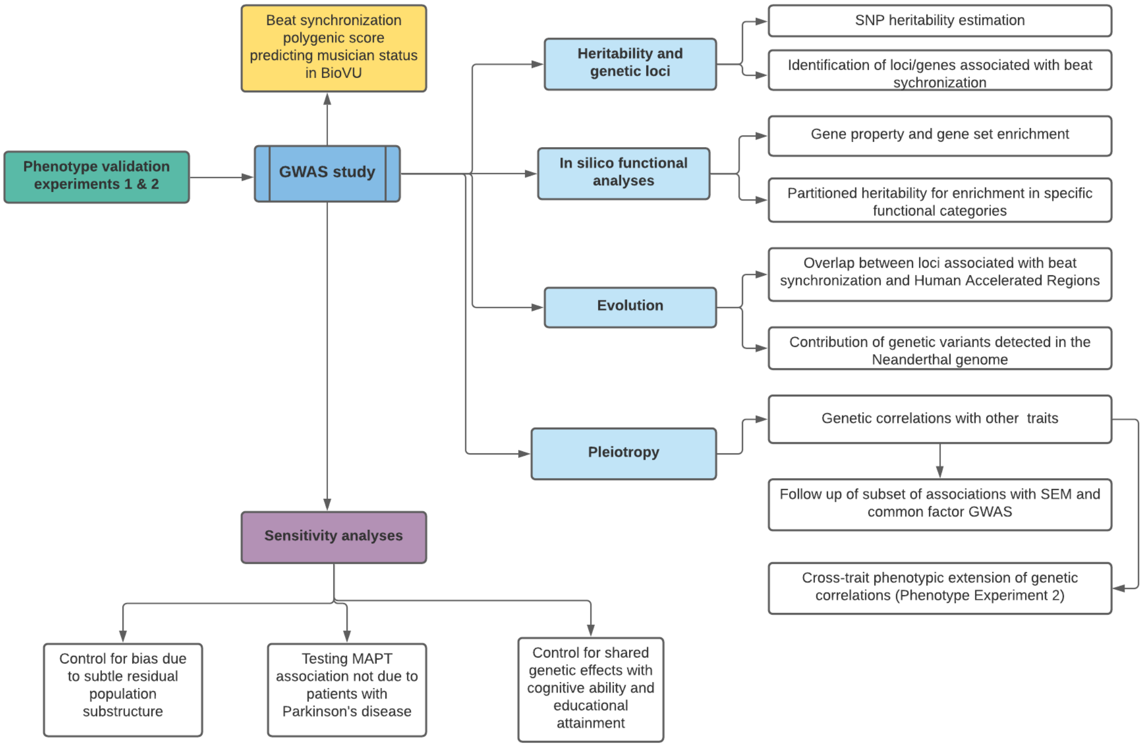

**Supplementary Figure 1.** We performed two phenotypic experiments to validate the self-report beat synchronization item (i.e., the single item 'Can you clap in time with a musical beat?' that was used in the genetic study), in relation to (task-based) measured rhythm perception and beat production and other musicality questionnaires. We investigated the genetic architecture of beat synchronization by performing a GWAS study of this self-report phenotype in 606,825 individuals. We estimated the SNP heritability, and identified loci/genes associated with beat synchronization. To validate the polygenic model of beat synchronization in relation to musicality in a separate sample, we constructed polygenic scores of beat synchronization within Vanderbilt's biobank, and tested whether they are associated with musician status. We then performed a series of in silico analyses including gene set enrichment analyses and gene property analyses to examine potential biological functions associated with beat synchronization. To determine whether the heritability of beat synchronization is enriched for specific functional categories, we used stratified LDSC, and LDSC-Specifically Enriched Genes (LDSC-SEG) to partition heritability. Given evolutionary hypotheses about the origins of rhythm, we analyzed the overlap between loci associated with beat synchronization and Human Accelerated Regions, and examined the contribution of genetic variants detected in the Neanderthal genome (present in modern Eurasians due to interbreeding) to the heritability of the beat synchronization phenotype. We also examined pleiotropy by testing genetic correlations with a curated list of traits, following up on certain associations with Structural Equation Modeling and common factor GWAS, and also tested cross-trait phenotypic replications of genetic correlations using data from the Phenotype Experiment 2. Finally, we conducted four types of sensitivity analyses to rule out potential the following potential biases and demonstrated therein that: 1) the GWAS beat synchronization results are not due to shared genetic effects with educational attainment or cognitive ability, 2) the GWAS genetic correlation results are not driven by subtle residual population substructure and 3) the MAPT association with beat synchronization is not driven by the presence of patients with Parkinson’s in the sample.

**Supplementary Figure 2. Task instructions for internet-based rhythm perception task in A. Phenotype validation Experiment #1 and B. Phenotype validation Experiment #2.**
**A.** Participants were instructed that in each trial, they would listen to the series of three rhythms (the first two were always identical, and the third could be the same or different), and they had to indicate if the third rhythm was the same or different.

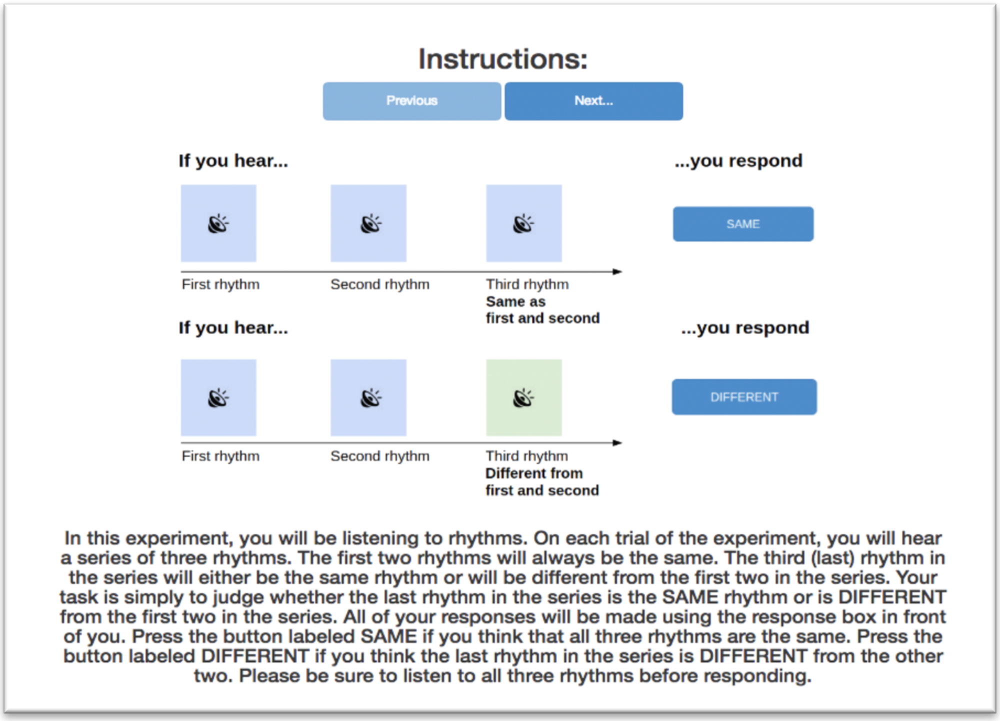

**B.** Participants were instructed that in each trial, they would listen to a music clip and their goal was to tap in time with the beat until the music ends. Participants were also informed that at the beginning of each clip they would hear a metronome to help them find the beat of the music. The tapping task consisted of 8 trials (4 music excerpts of 30 seconds each, and each excerpt occurring twice).

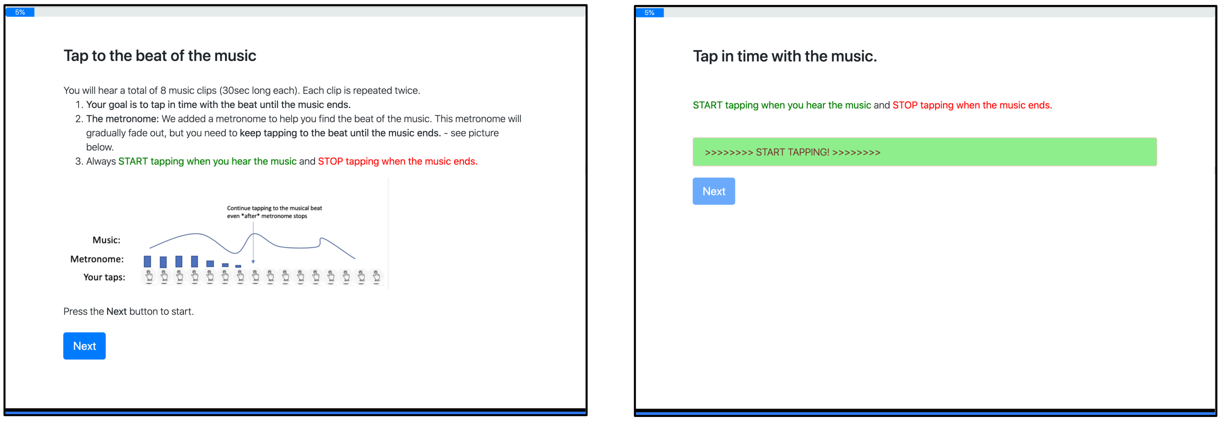

**Supplementary Figure 3.** As a supplemental analysis, we computed vector lengths as a tapping accuracy variable from the beat synchronization task in phenotype experiment #2. Key analyses to examine correlations with beat self-report demonstrated similar results to analyses in which SD of the Asynchrony had been used (note that higher vector length is indicative of more accurate tapping performance). From top left: association between vector length and the GWAS target question (McFadden's $R^2$ = 0.21, p<0.001); association between vector length and a similar self-report question asked on a Likert scale (r=0.37, p<0.001); association between vector length and a multi-item rhythm questionnaire (r=0.37, p<0.001); and association between Musical Sophistication (Gold-MSI) and vector length (r=0.31, p<0.01). See Supplementary Notes for details.

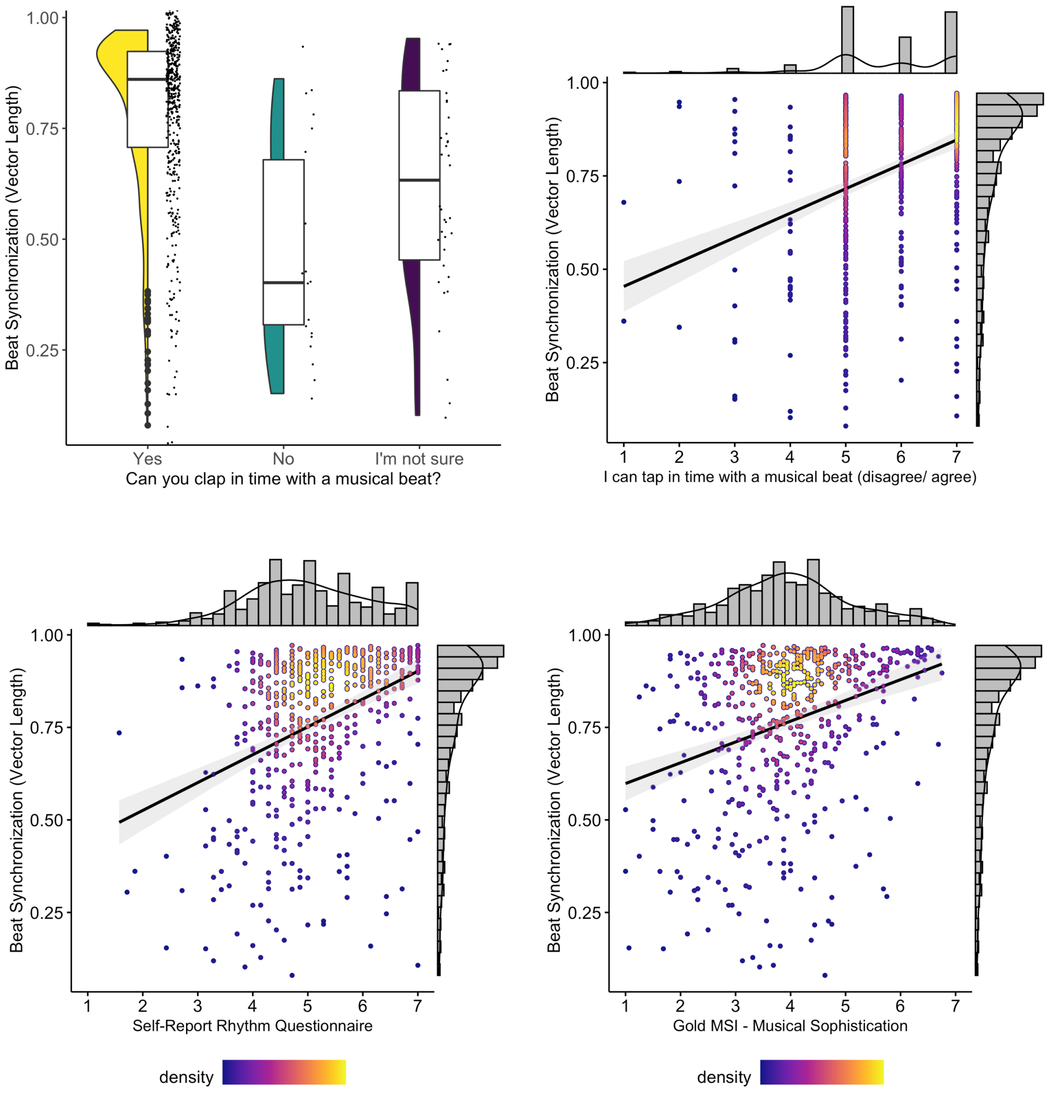

**Supplementary Figure 4. Q-Q plot of beat synchronization GWAS results.** The Quantile-Quantile (QQ) plot shows observed vs. expected -$\log_{10}$p-values. The inflation observed in the QQ plot is likely due to polygenicity of the trait rather than to population stratification, given that the lambda and intercept indexes are within the expected range, and that when adjusting for population substructure (see Supplemental Notes and Supplementary Table 17, the betas of the GWAS remained virtually identical).

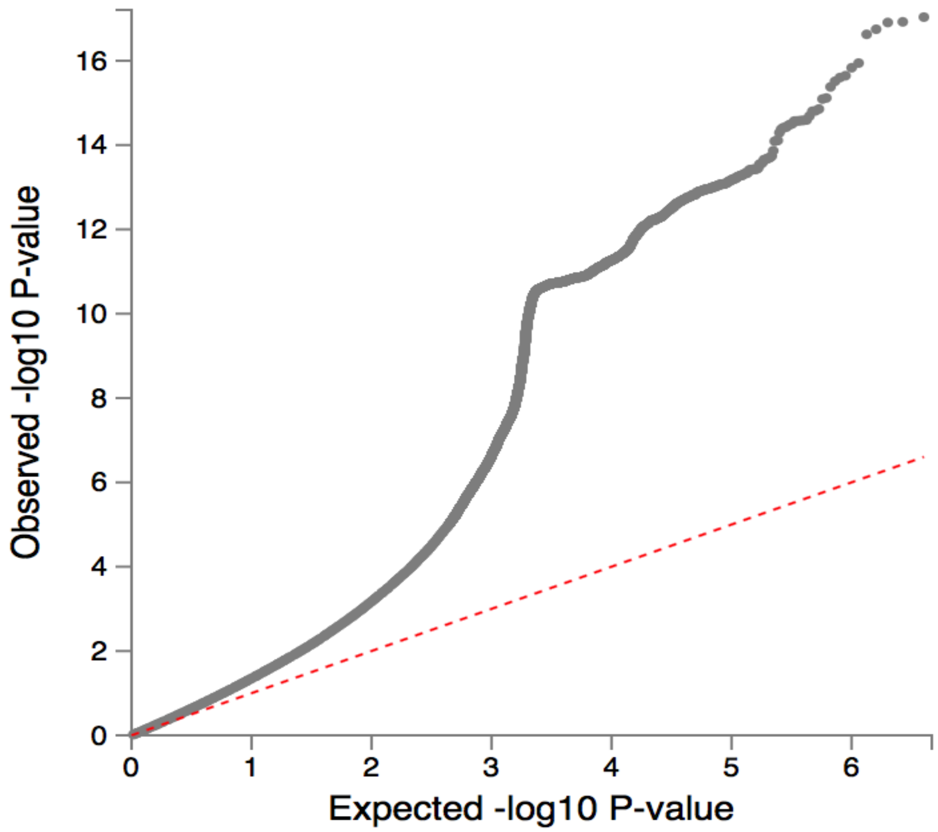

**Supplementary Figure 5. Results of multi-tissue enrichment of beat synchronization GWAS signal in enhancers.** Enhancer regions are defined as peaks of histone H3 lysine 4 monomethylation (H3K4me1). Partitioned heritability analysis was performed in LDSC-SEG (Finucane et al., 2018, Nature Genetics).

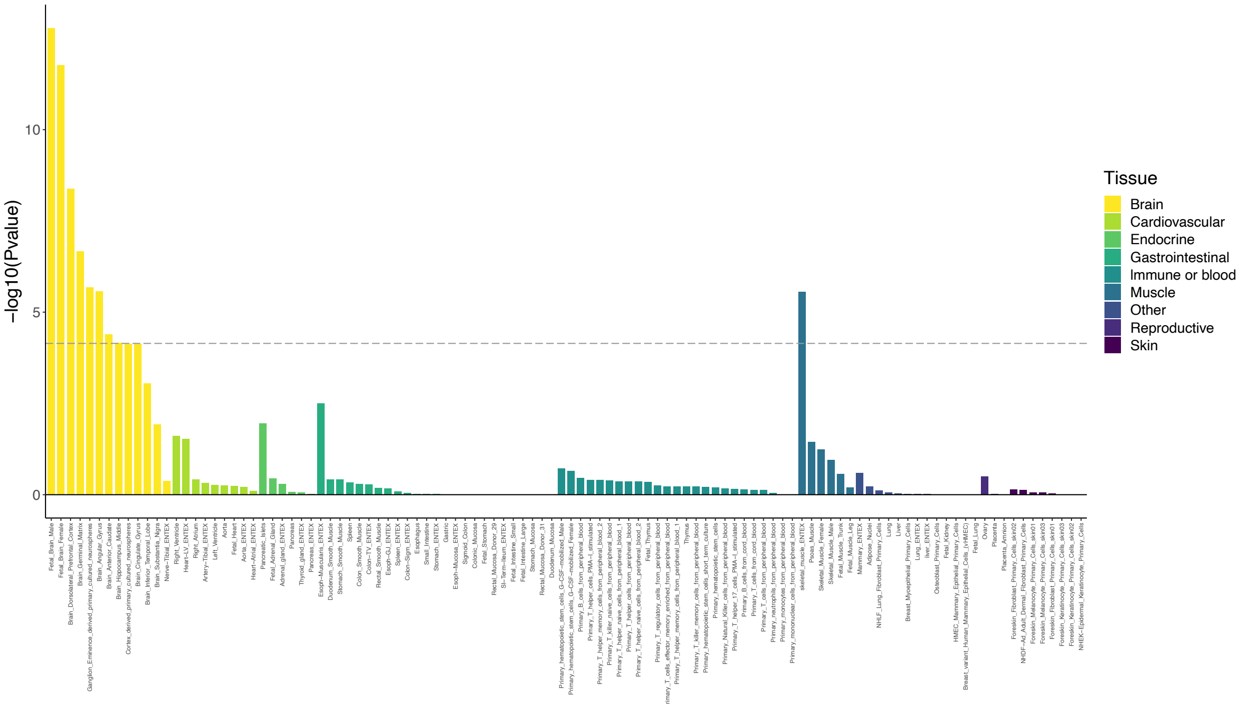

**Supplementary Figure 6. Enrichment of beat synchronization GWAS signal in active chromatin regions.** Active chromatin is defined as peaks of DNase hypersensitivity, and histone H3 acetylation at lysine 9 (H3K9ac) and lysine 27 (H3K27ac). Enrichment was calculated using LDSC-SEG partitioning heritability analysis (Finucane et al., 2018). The dashed line represents the Bonferroni corrected significance threshold.

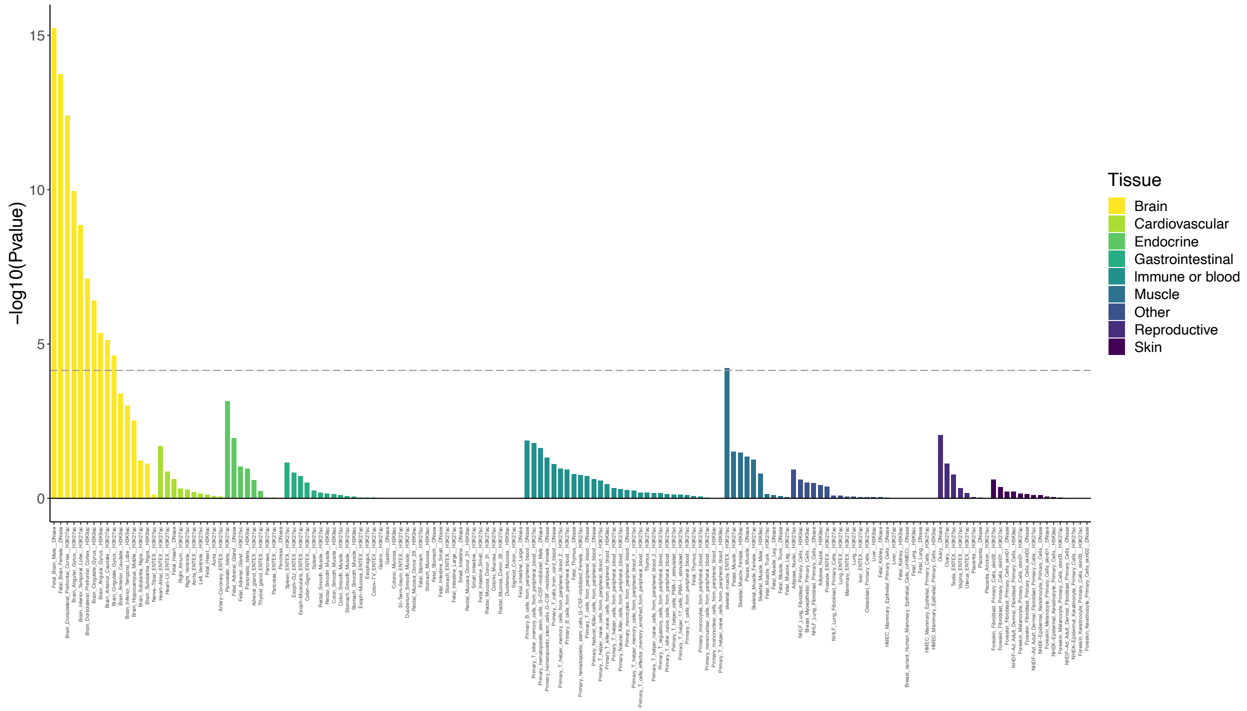

**Supplementary Figure 7: Enrichment of beat synchronization GWAS signal in multi-tissue gene expression set,** using LDSC-SEG. Results show enrichment in several types of brain tissue, conditional on other annotations. The dashed line represents the Bonferroni corrected significance threshold.

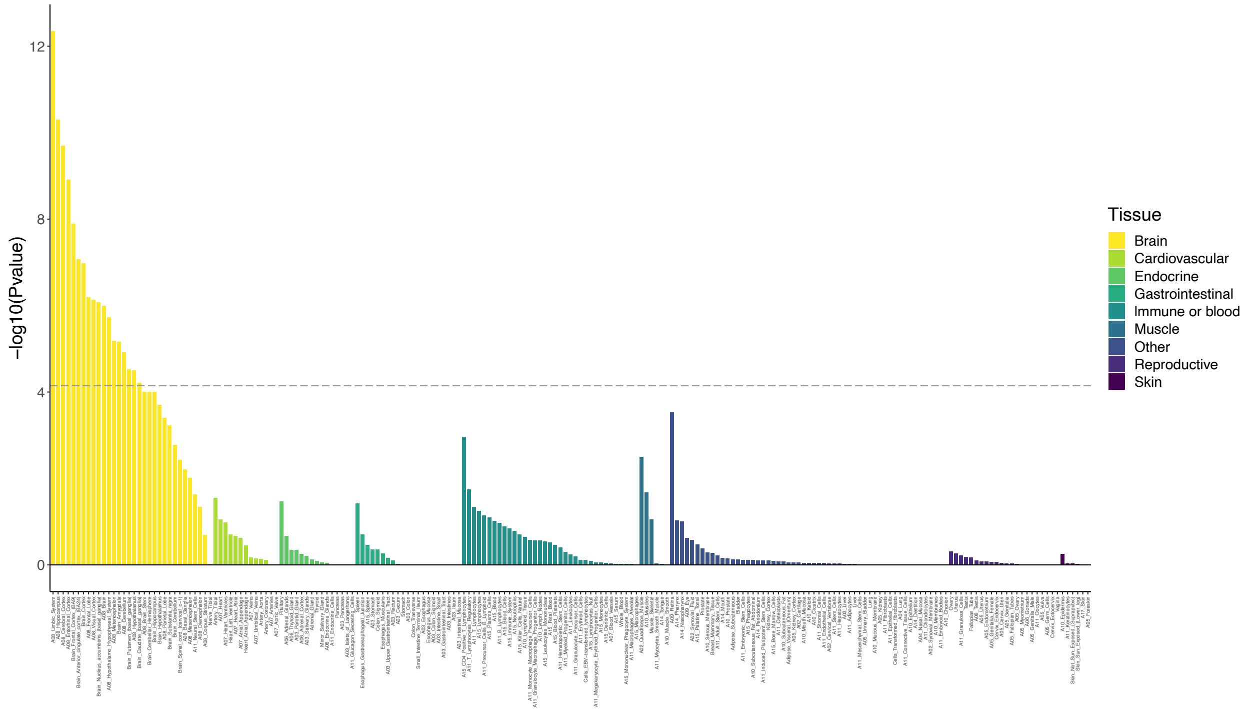

**Supplementary Figure 8. Manhattan plot of Multivariate Rhythm-Related Traits GWAS (based on the common factor from Genomic SEM analyses).** Genome-wide significance level is indicated on red dotted line.

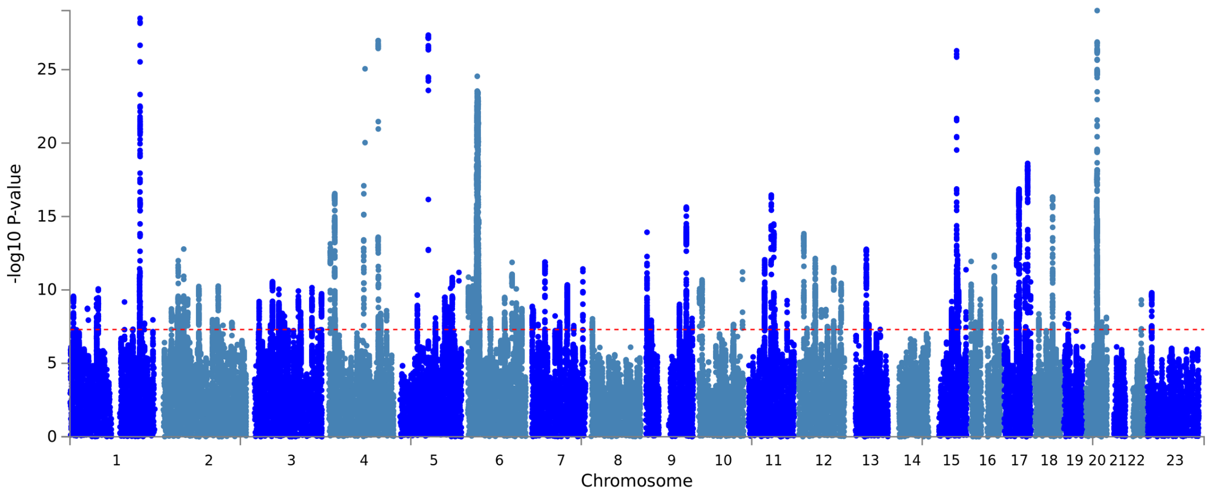

**Supplementary Figure 9. The common factor rhythm-related traits multivariate GWAS is enriched for cerebellum and pituitary tissue expression.** Results of MAGMA gene-property analysis based on gene expression levels from GTEx of 54 tissues. Associations were significantly enriched in brain-expressed genes compared to other tissues (-log-10 p-values are on the y-axis, with tissue type on the x-axis). Dotted line shows p-value threshold for significant enrichment after Bonferroni correction.

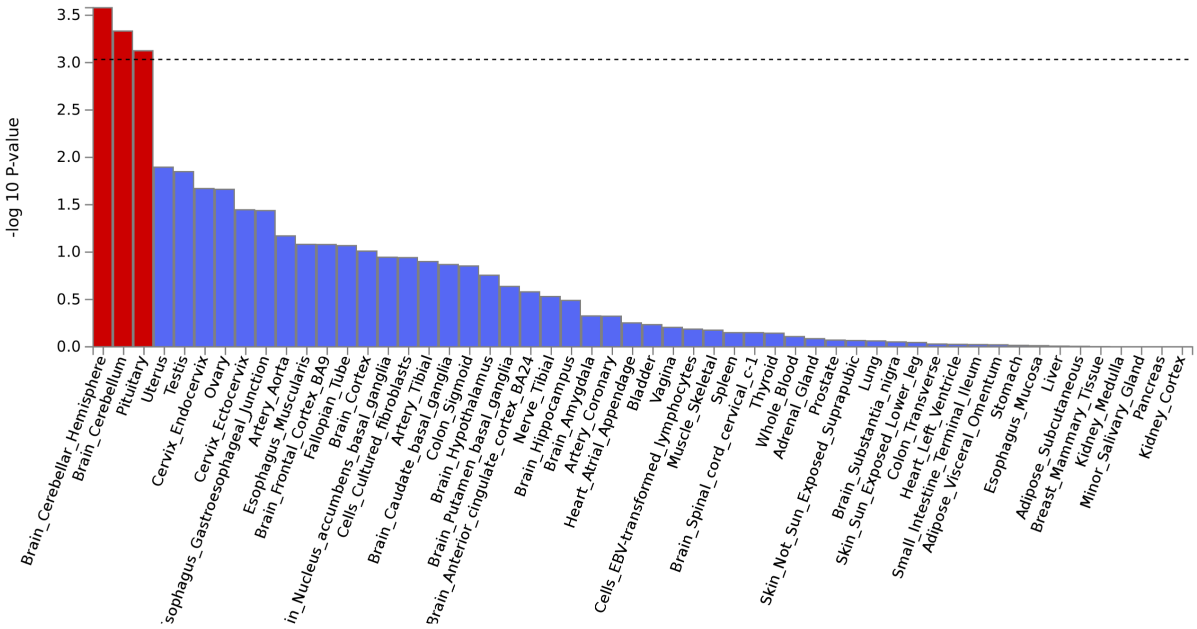

**Supplementary Figure 10. Miami plot showing the beat synchronization GWAS results unadjusted (top) and adjusted (bottom) for A. general cognitive ability and B. educational attainment**.

**A. Beat synchronization GWAS adjusted for general cognitive ability.** We used mtCOJO to condition the beat synchronization GWAS summary statistics on GWAS of general cognitive ability (de la Fuente et al[25]). The results remain largely unchanged; 62 of the original 69 genomic loci still surpass the criteria for genome-wide significance ($p<5x10^{-8}$) after adjusting for general cognitive ability (see Supplementary Table 14). For illustration purposes we only present 500,000 SNPs with p<0.1

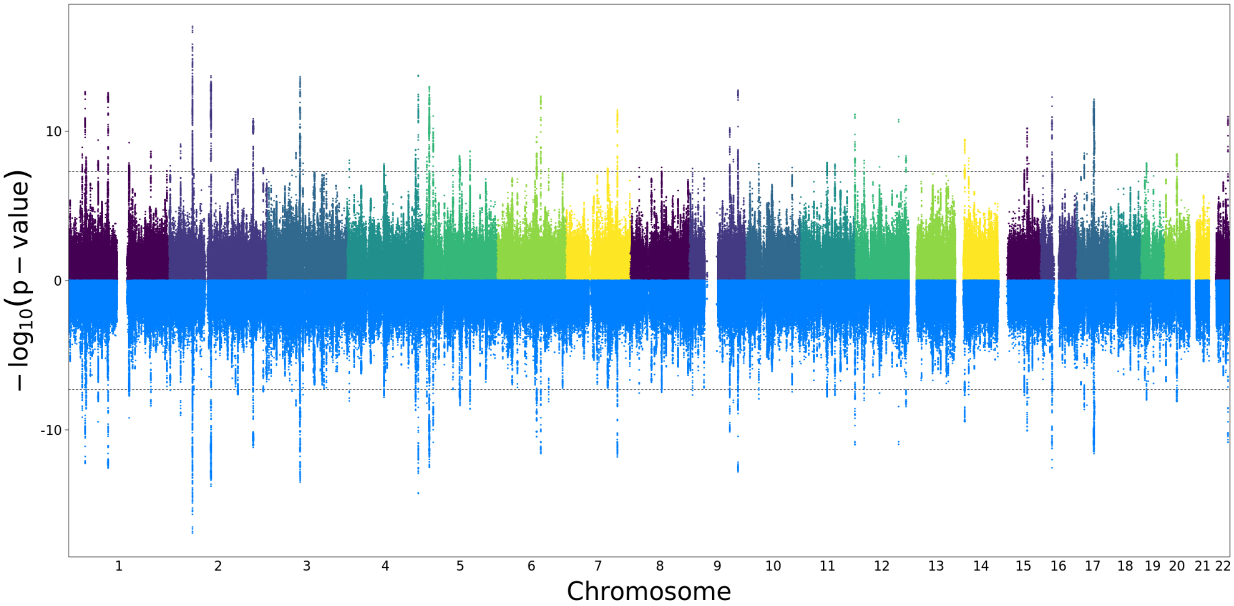

**B. Beat synchronization GWAS adjusted for educational attainment.** We used mtCOJO to condition the beat synchronization GWAS summary statistics on GWAS of educational attainment (Lee et al.[68]). The results remain largely unchanged; 65 of the original 69 genomic loci still surpass the criteria for genome-wide significance ($p<5x10^{-8}$) after adjusting for educational attainment (see Supplementary Table 15). For illustration purposes we only present 500,000 SNPs with p<0.1

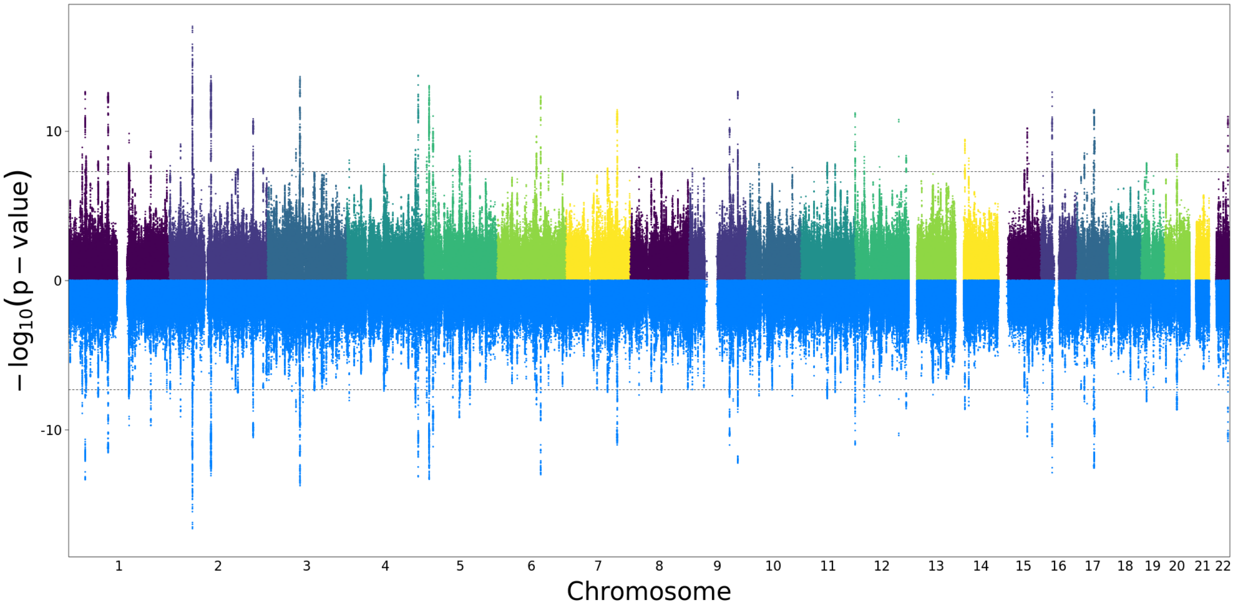
