## Supplementary material for "Genome-wide association study of musical beat synchronization demonstrates high polygenicity": Regional association plots

Regional association plots are presented below for each of the 69 genomic loci associated with rhythm that surpassed the threshold for genome-wide significance. Plots were generated for each locus using FUMA (setting: GWAS summary statistics), from the rhythm summary statistics (which had undergone two rounds of LD pruning, first at $r^2$=0.6 [independent SNPs], and another round of pruning at $r^2$ = 0.1, kb=250). In each plot, the top independent SNP for each locus is depicted as “Top lead SNP”, with a navy circle.

rs848293

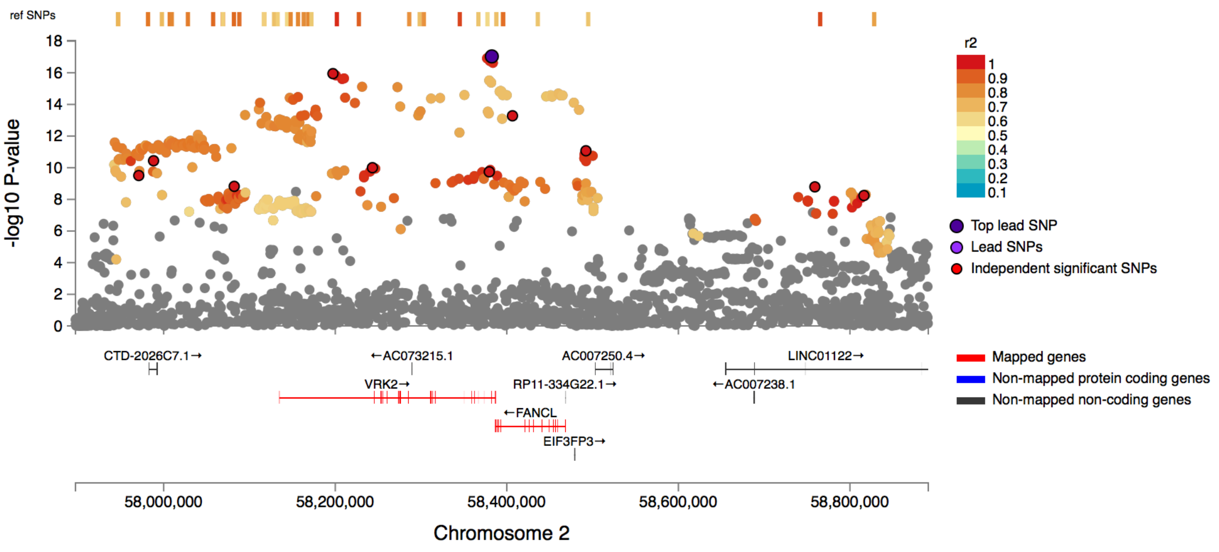


rs62340585

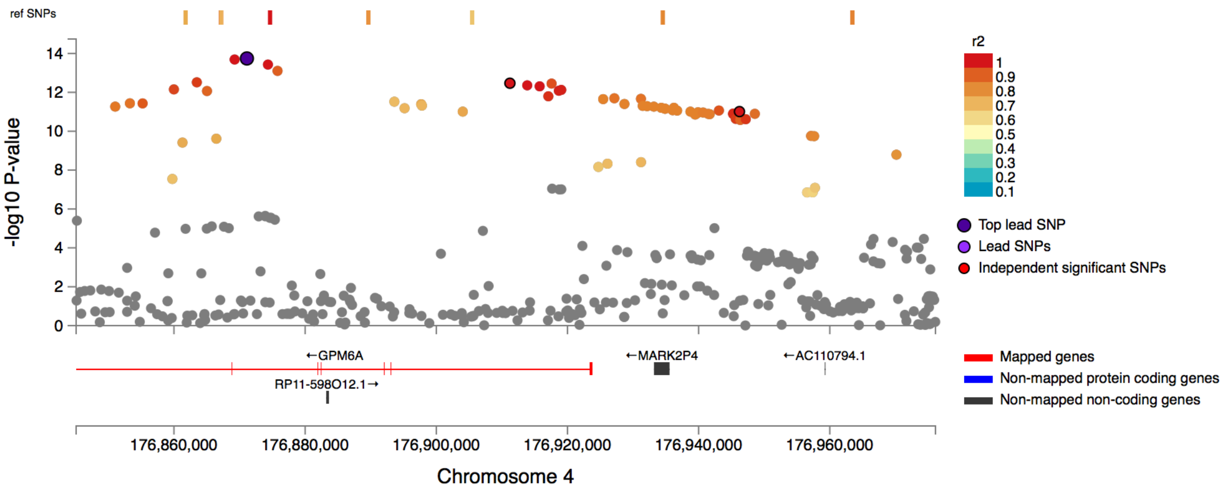

rs10168817

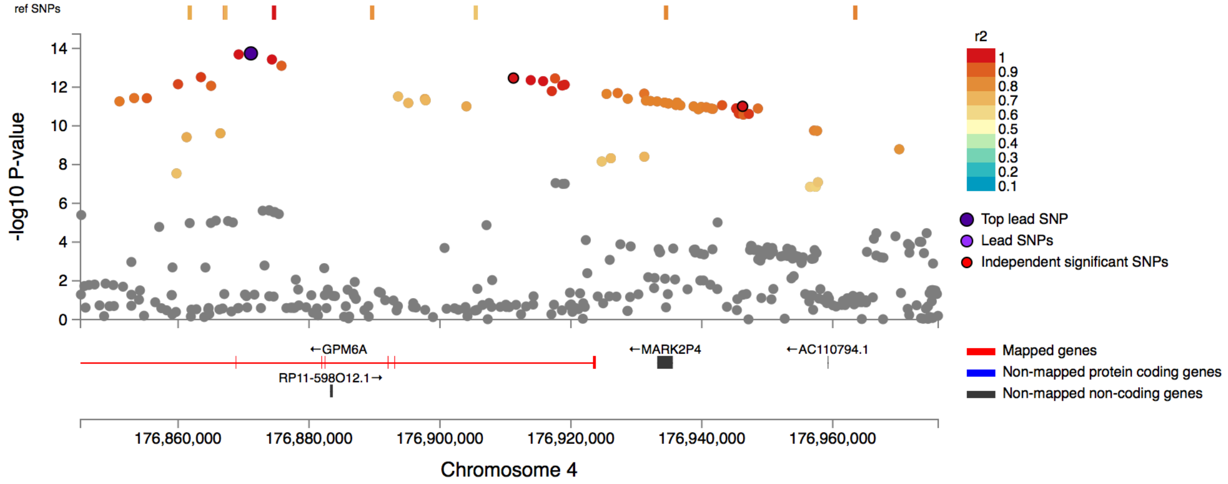


rs10779987

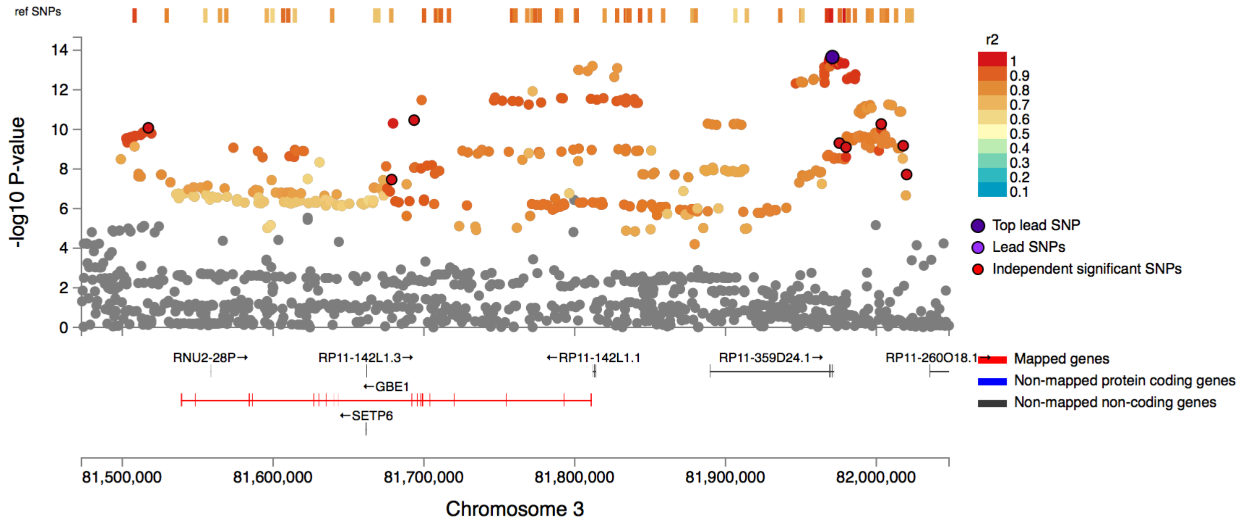

rs28392605

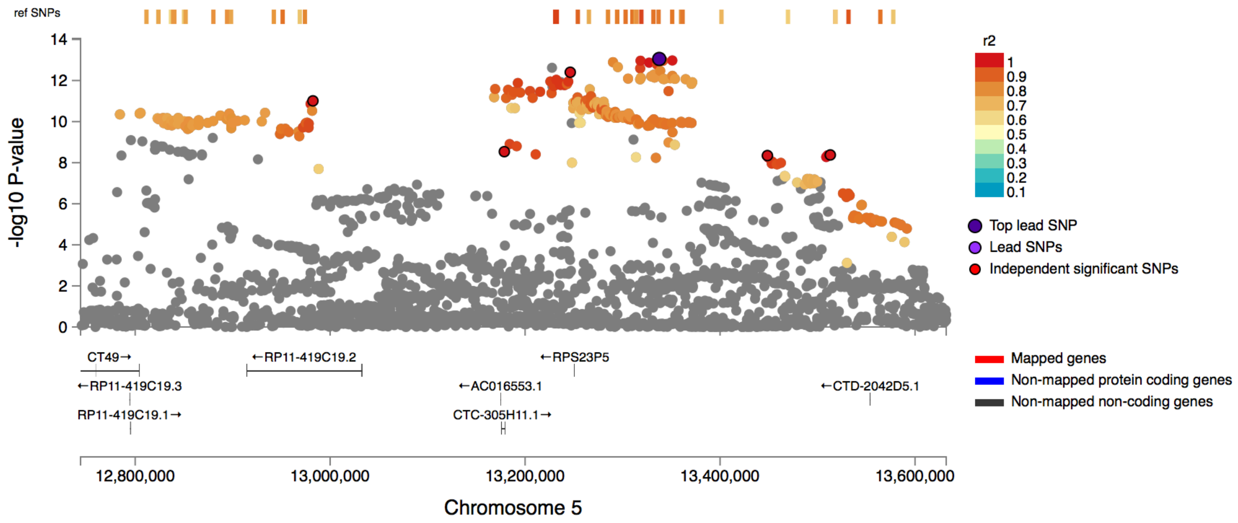


rs1832909

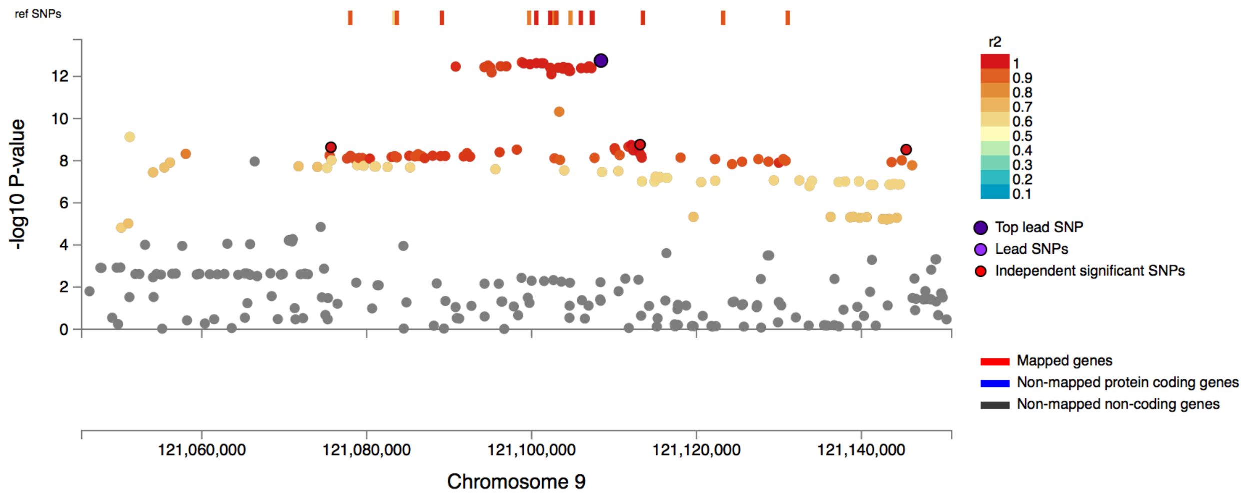

# rs34762587

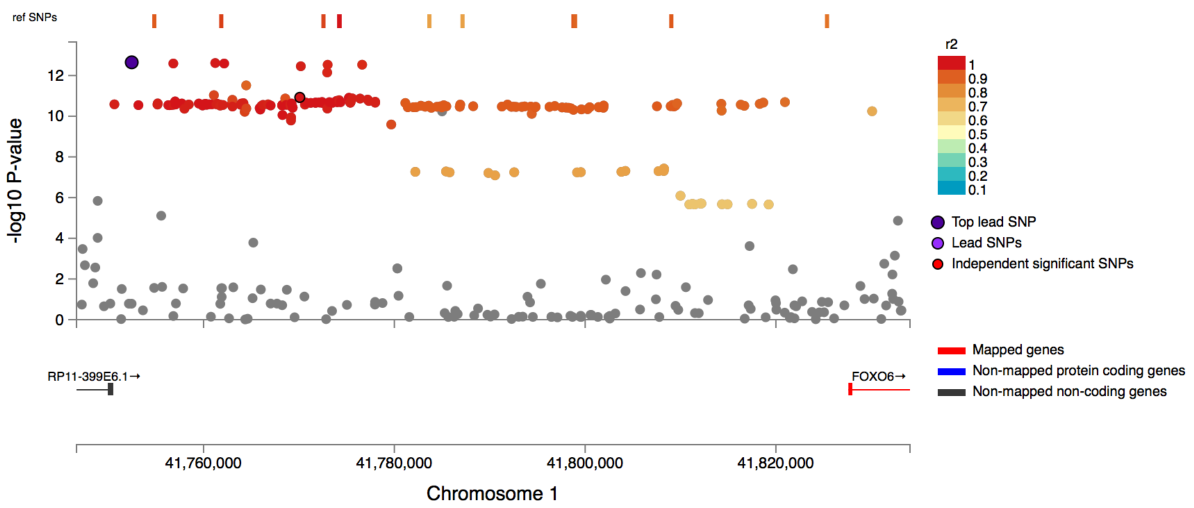


# rs7542

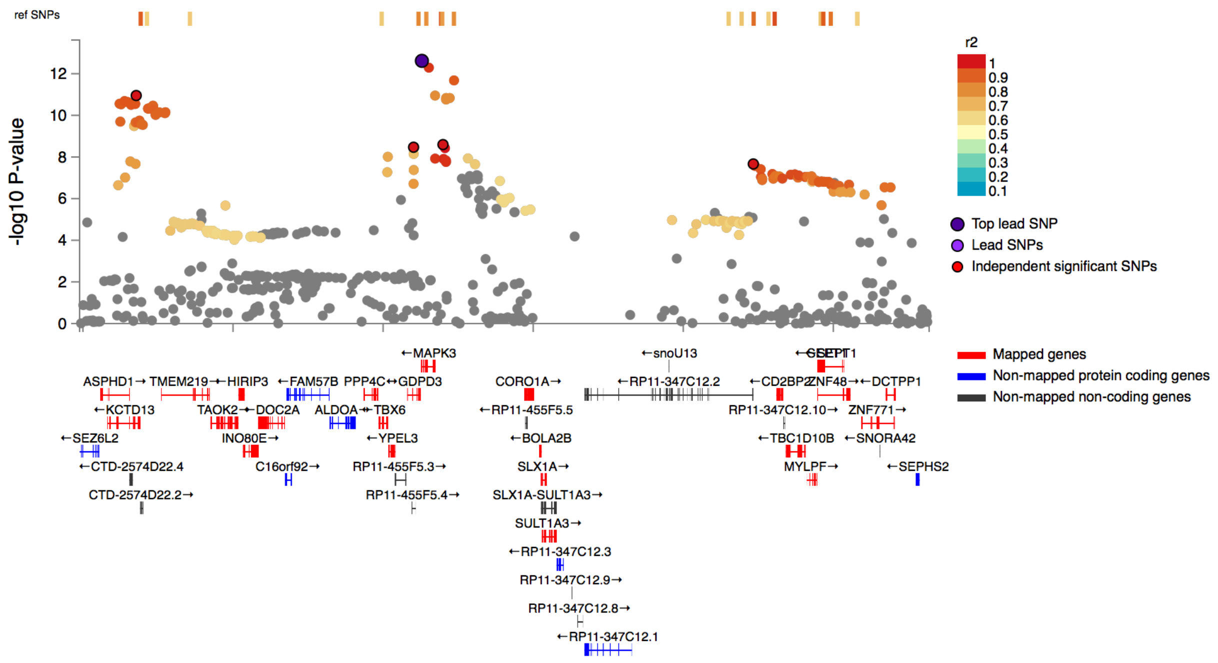

## rs10875125

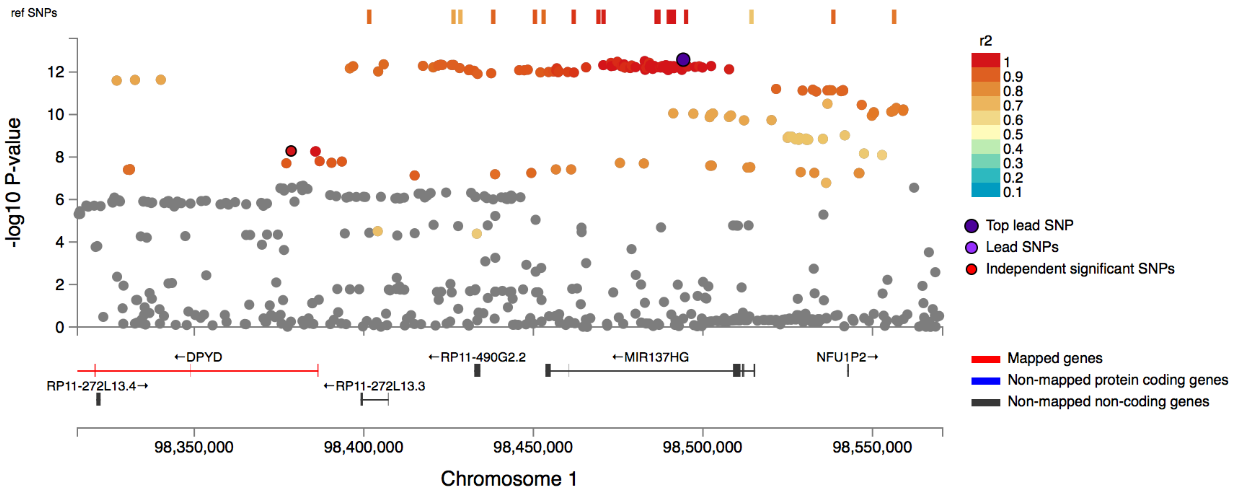


## rs9400241

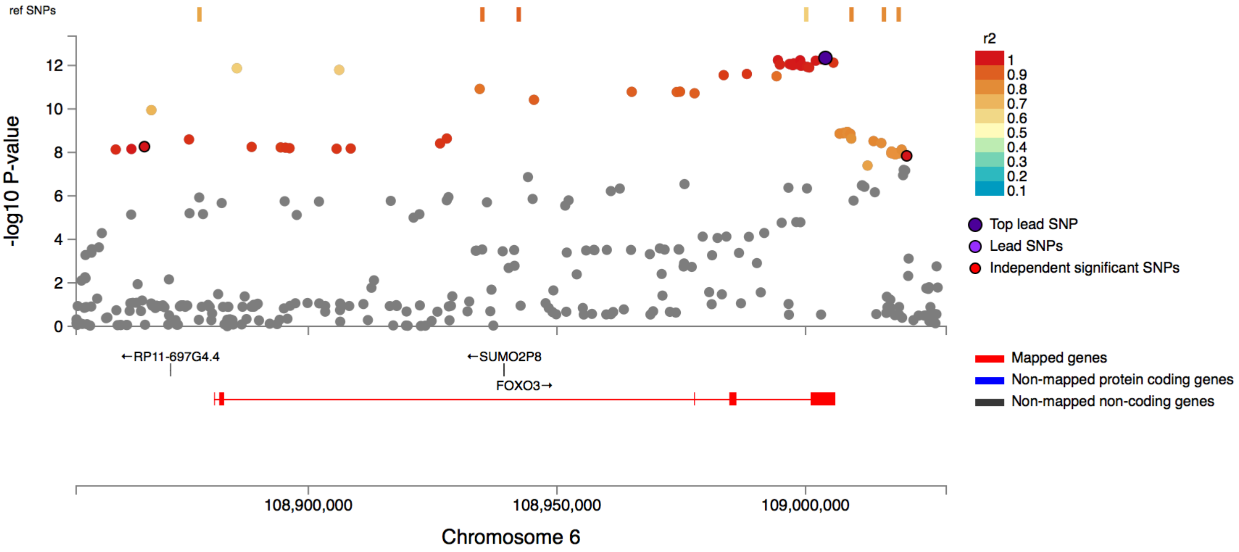

rs4792891

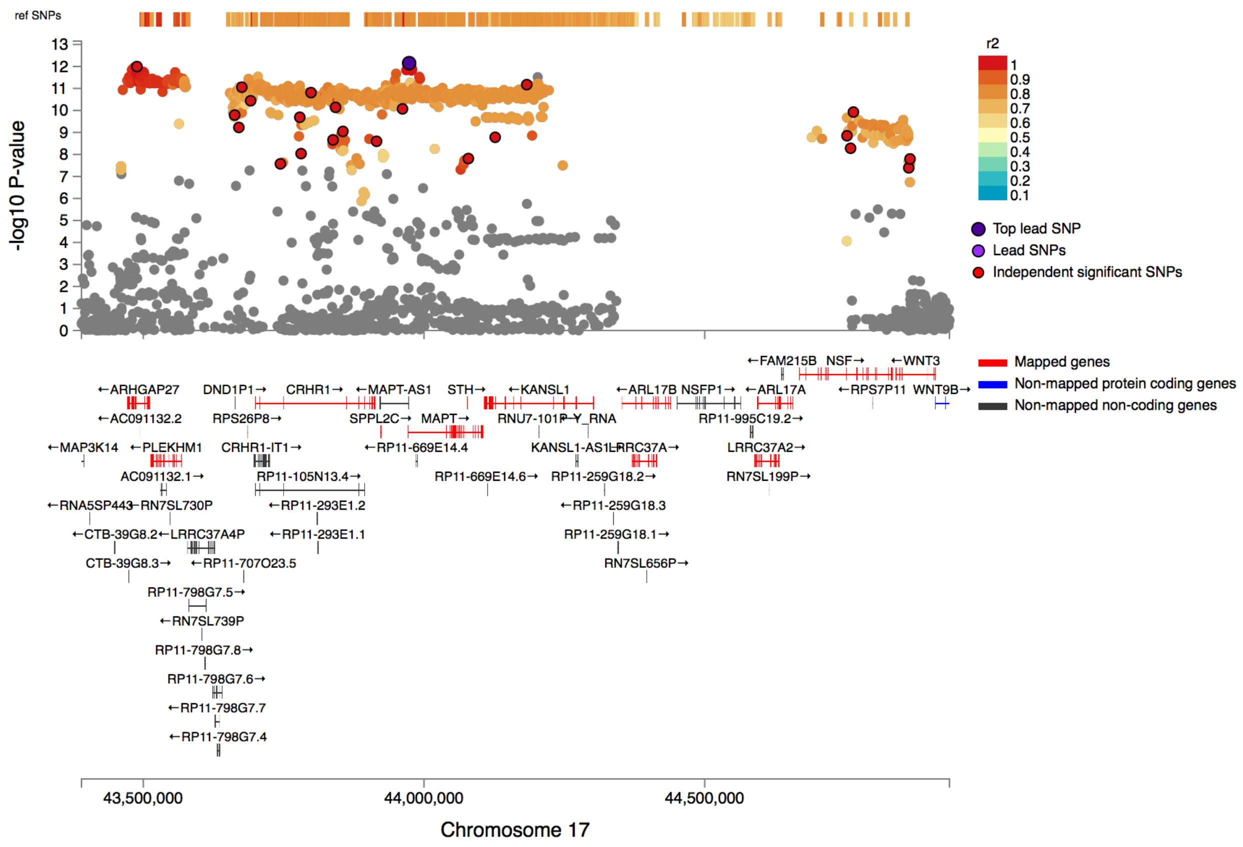
ref SNPs
-log10 P-value
r2
Top lead SNP
Lead SNPs
Independent significant SNPs
Mapped genes
Non-mapped protein coding genes
Non-mapped non-coding genes
43,500,000
44,000,000
44,500,000
Chromosome 17

rs1468701

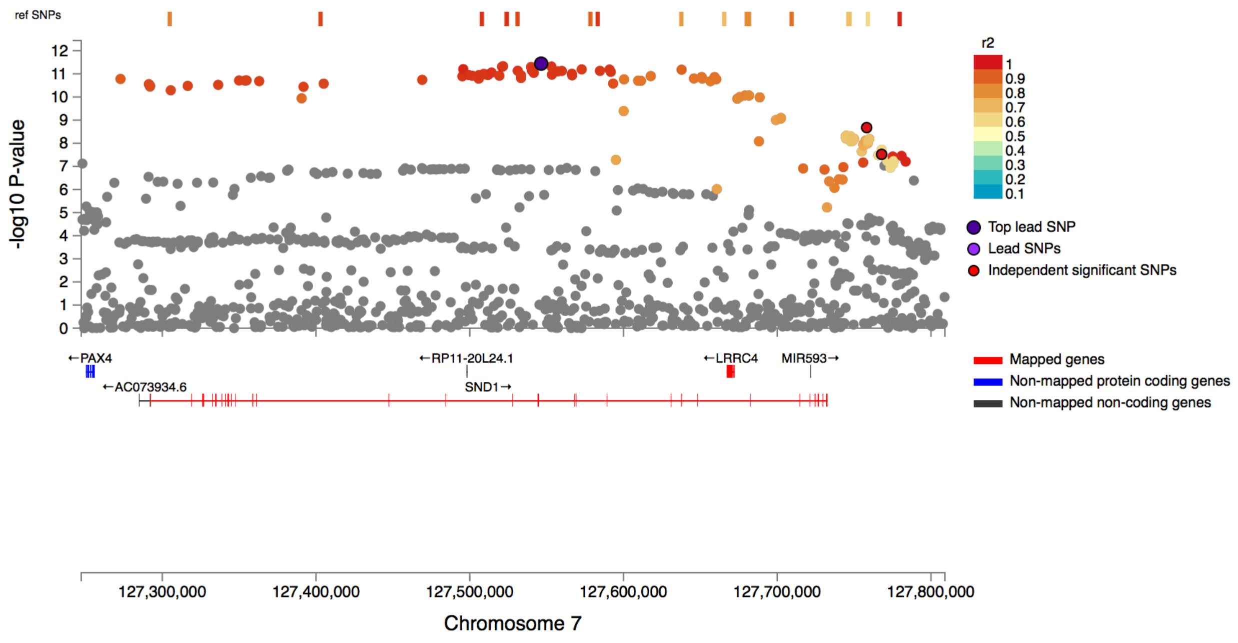


rs10848650

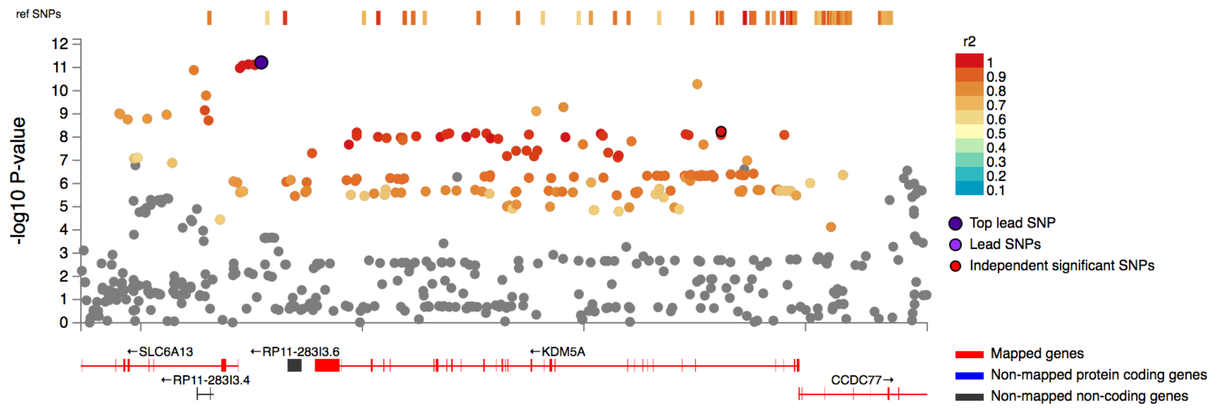

rs2635634

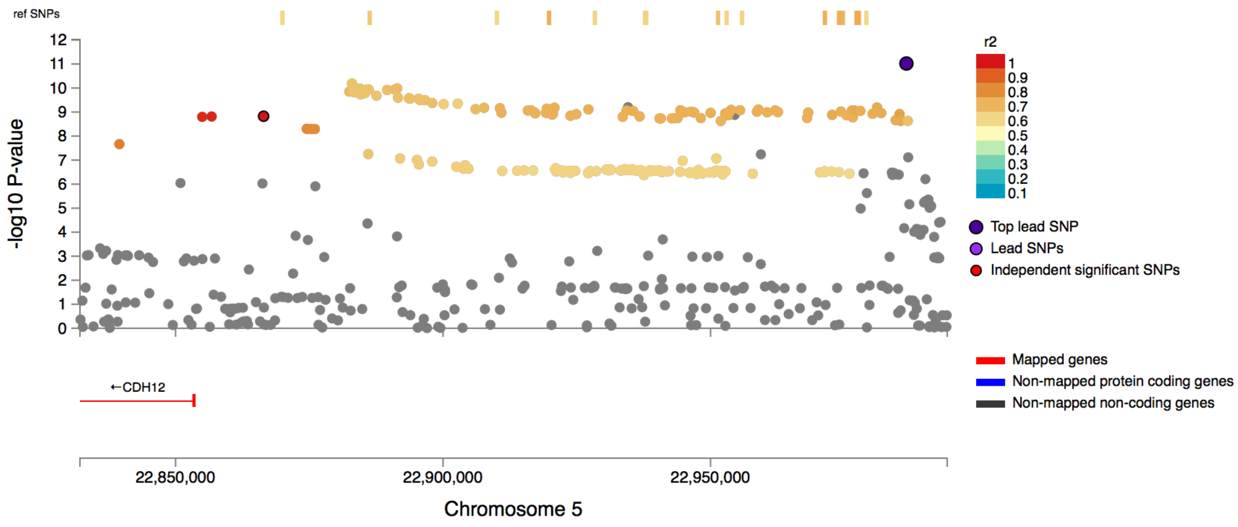


rs9626920

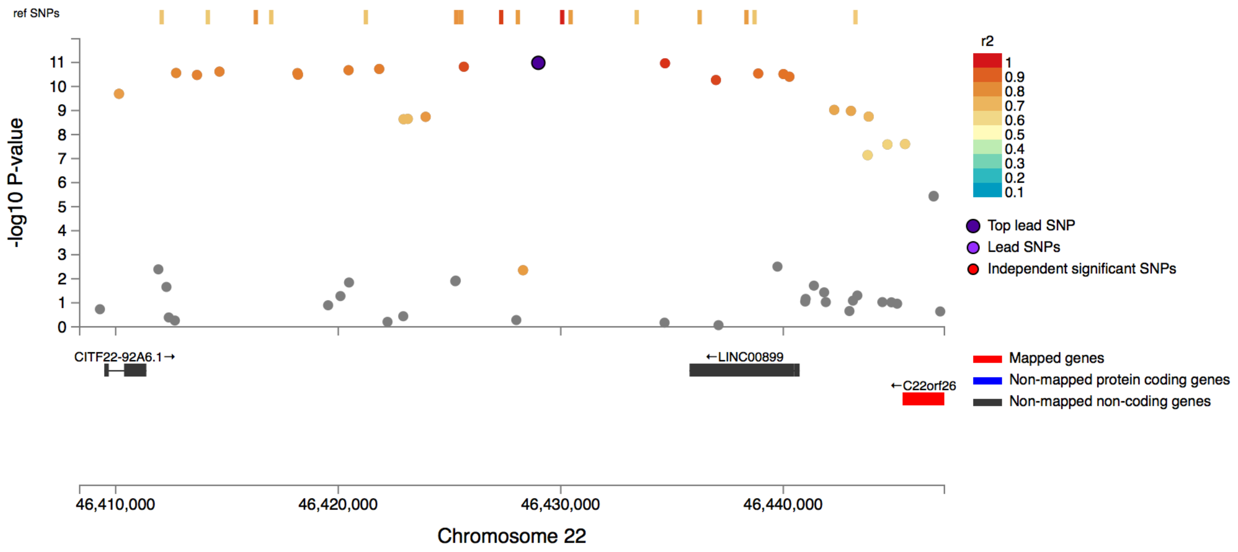

rs764299

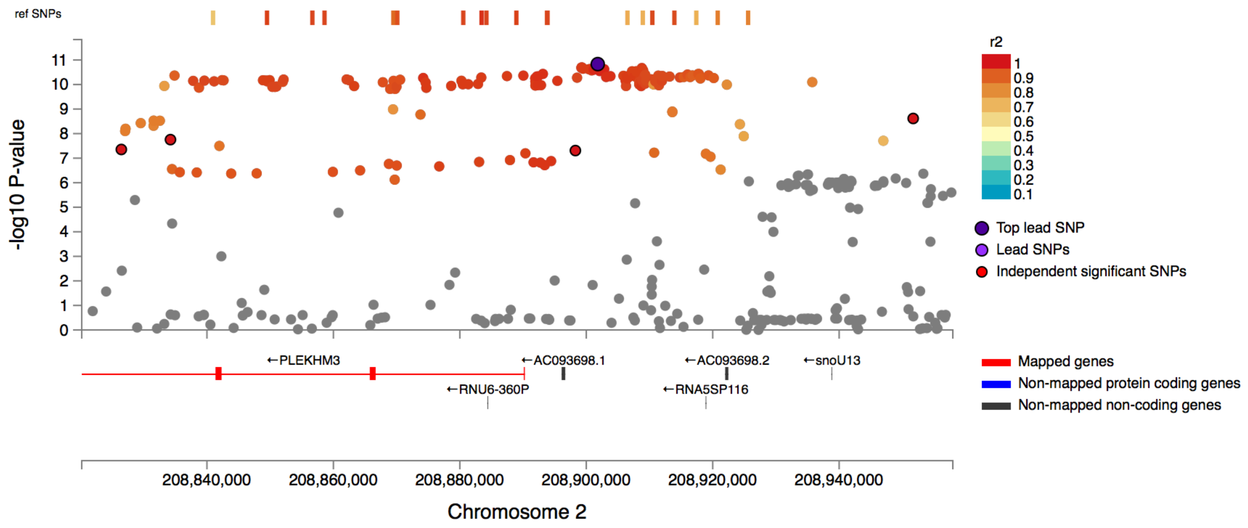


rs10984506

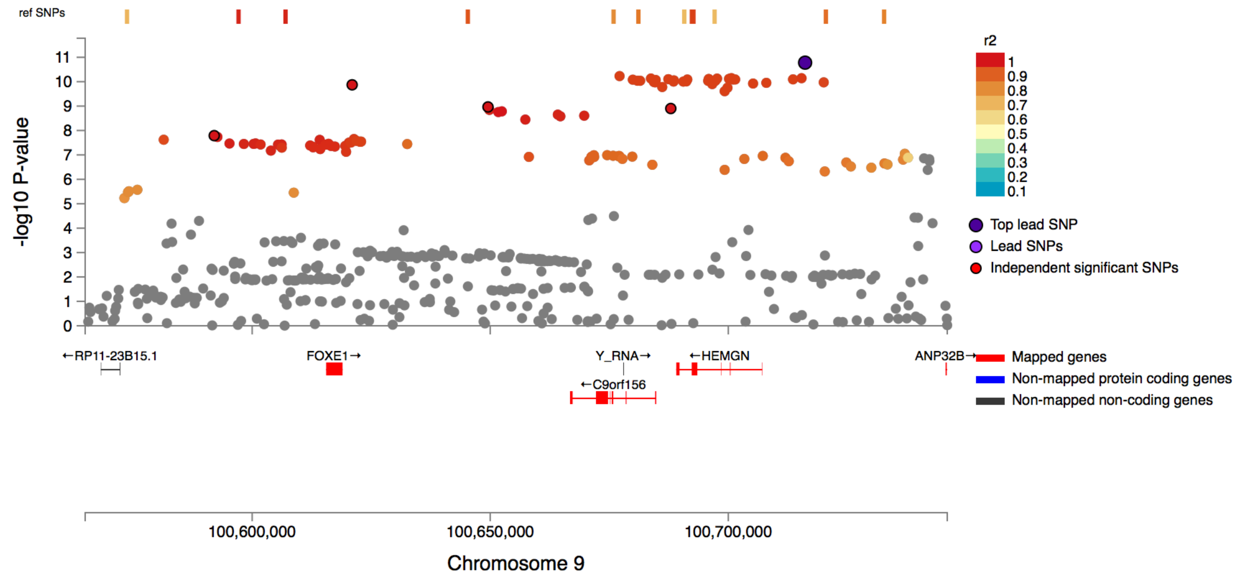

rs1426371

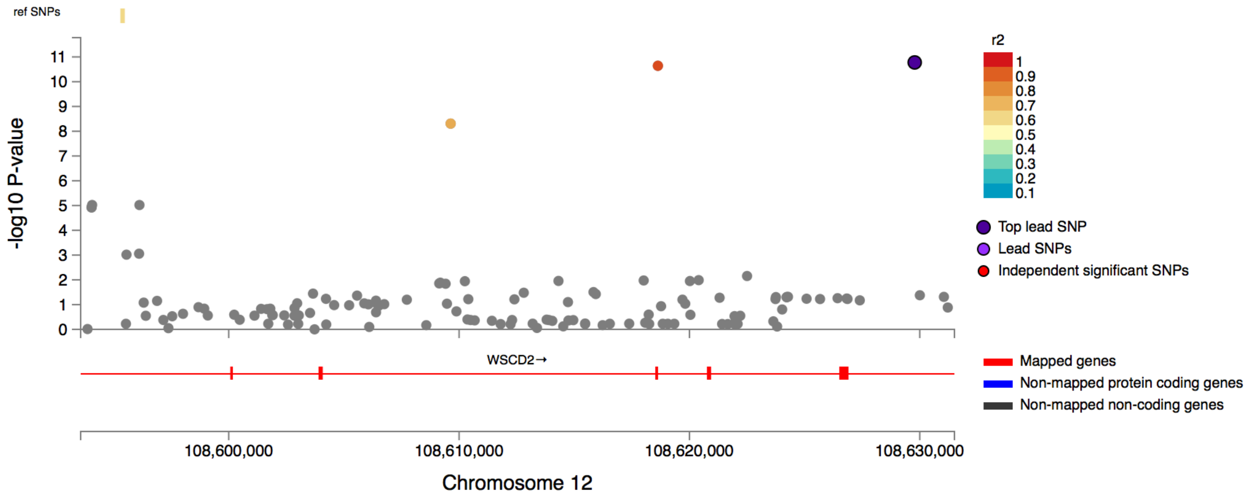


rs12913592

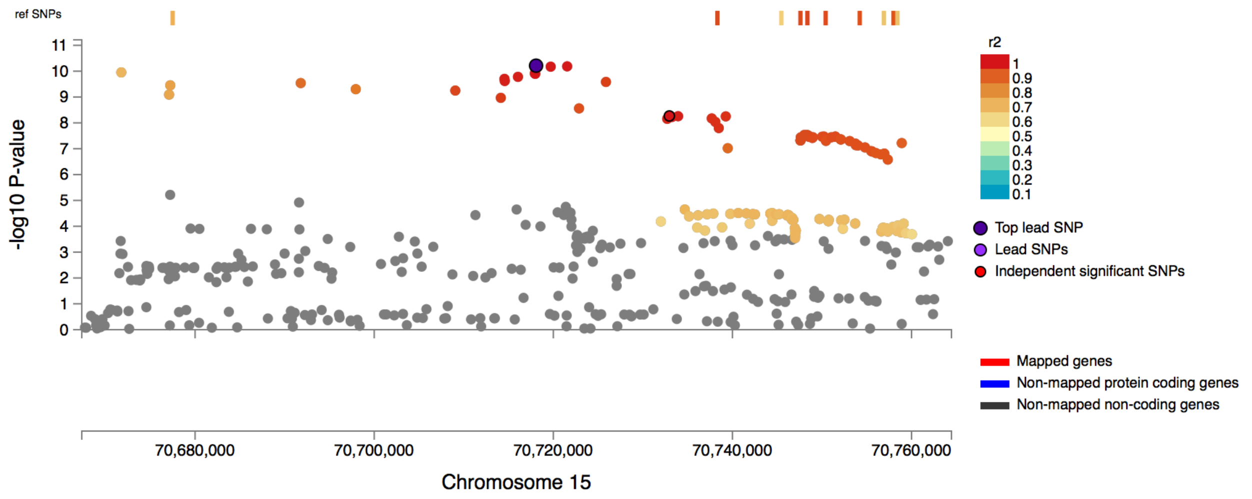

## rs72700870

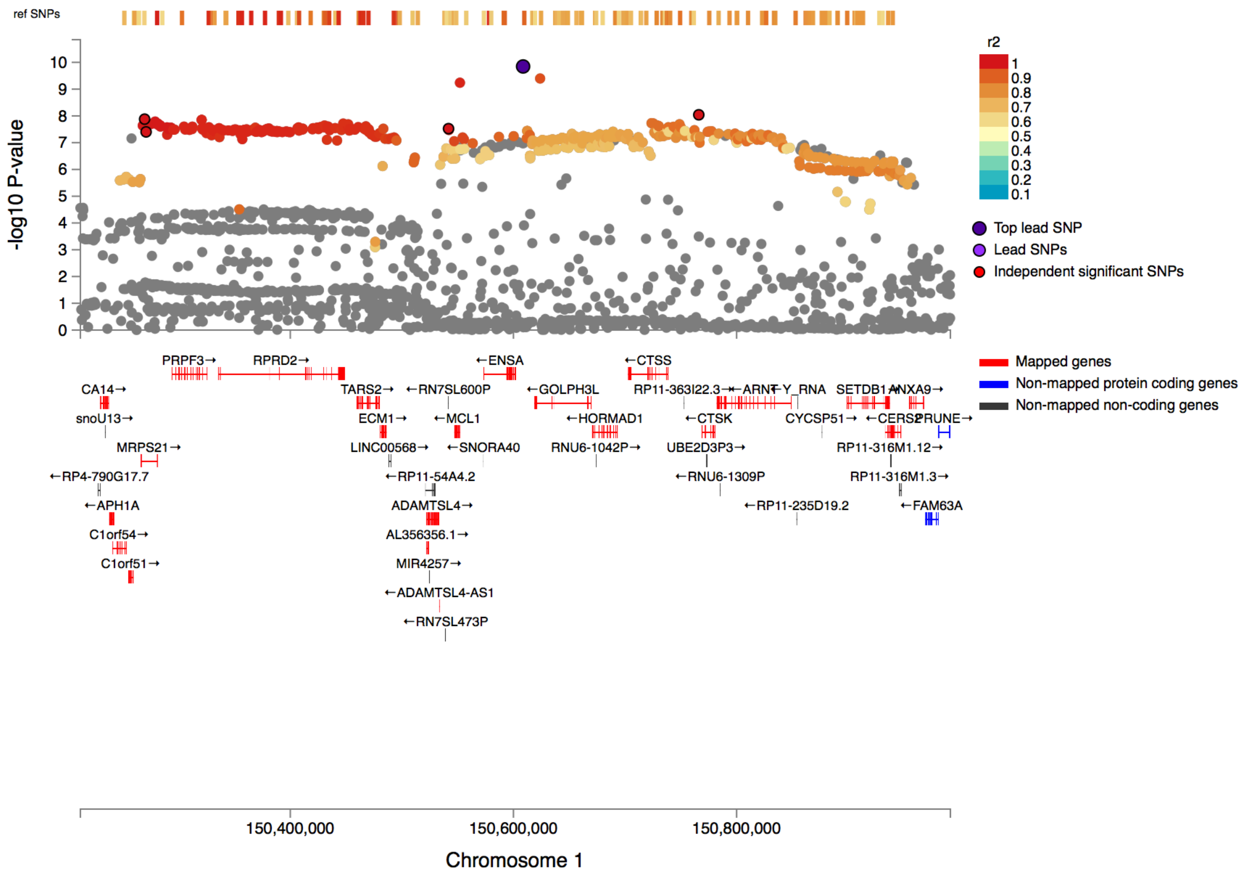


## rs9388171

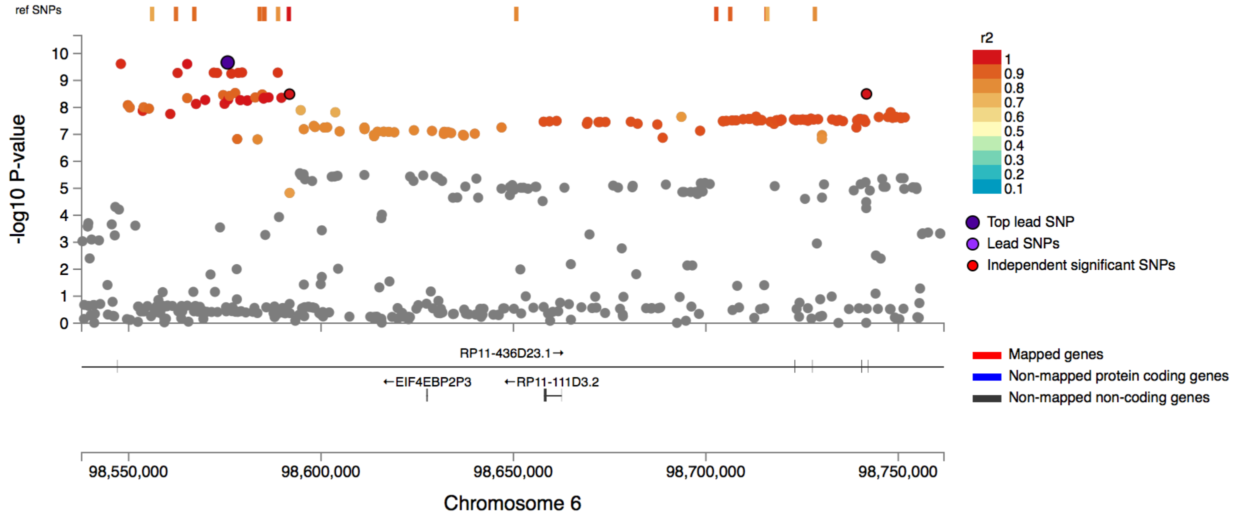

## rs6572878

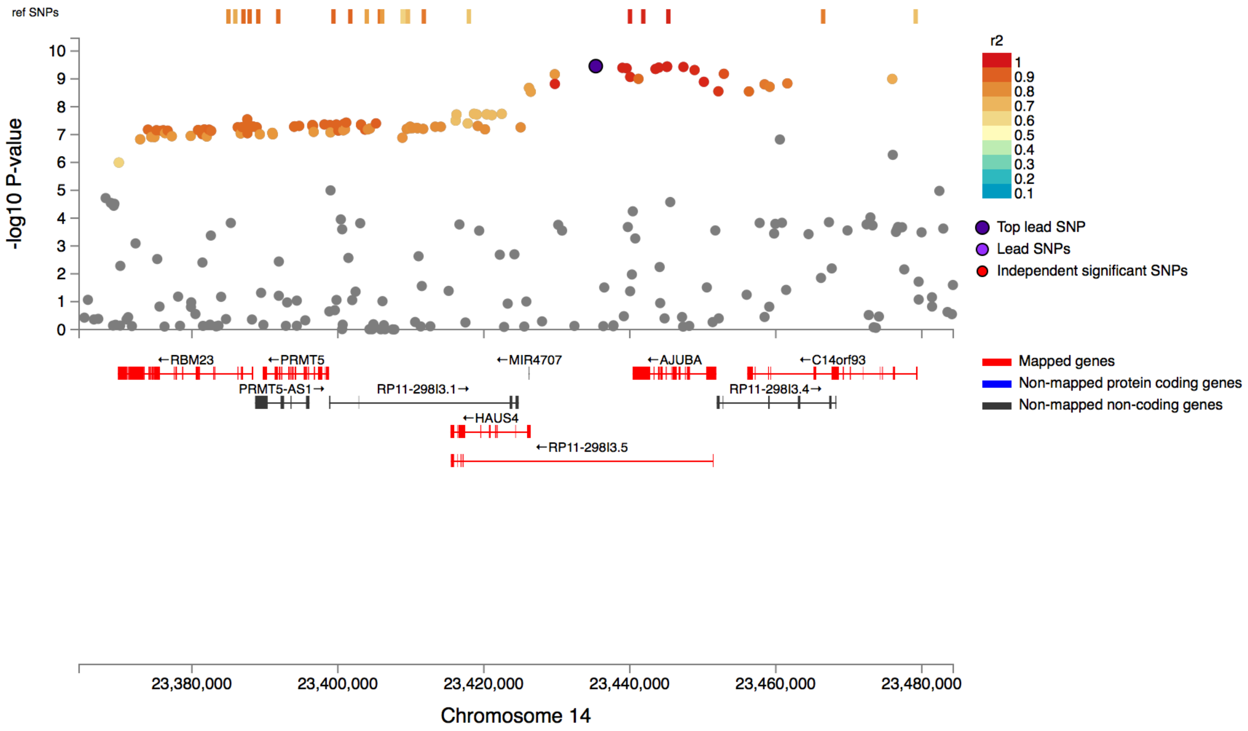


## rs11210206

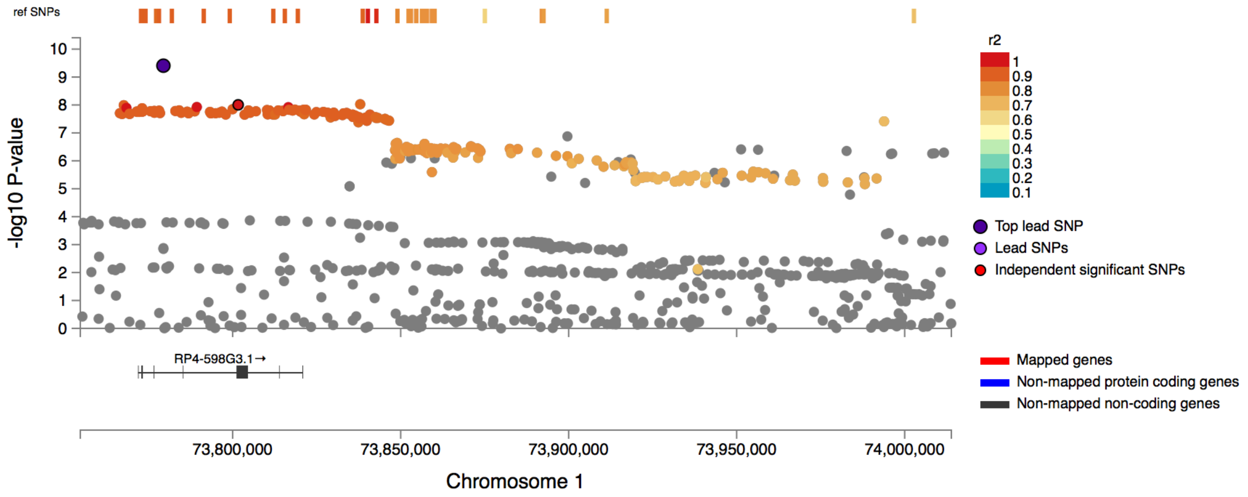

# rs72633496

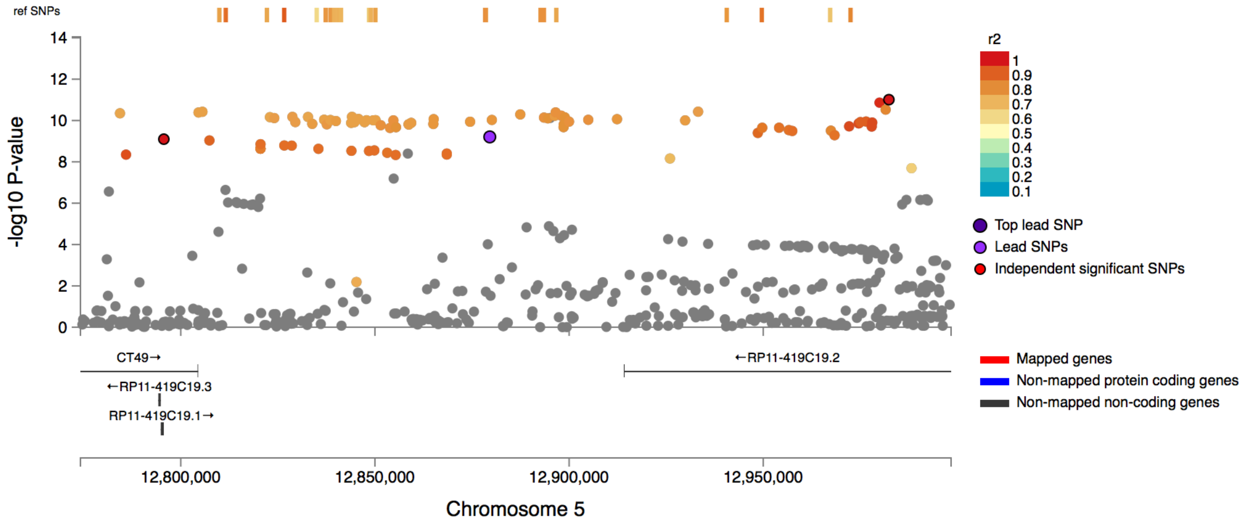


# rs7586405

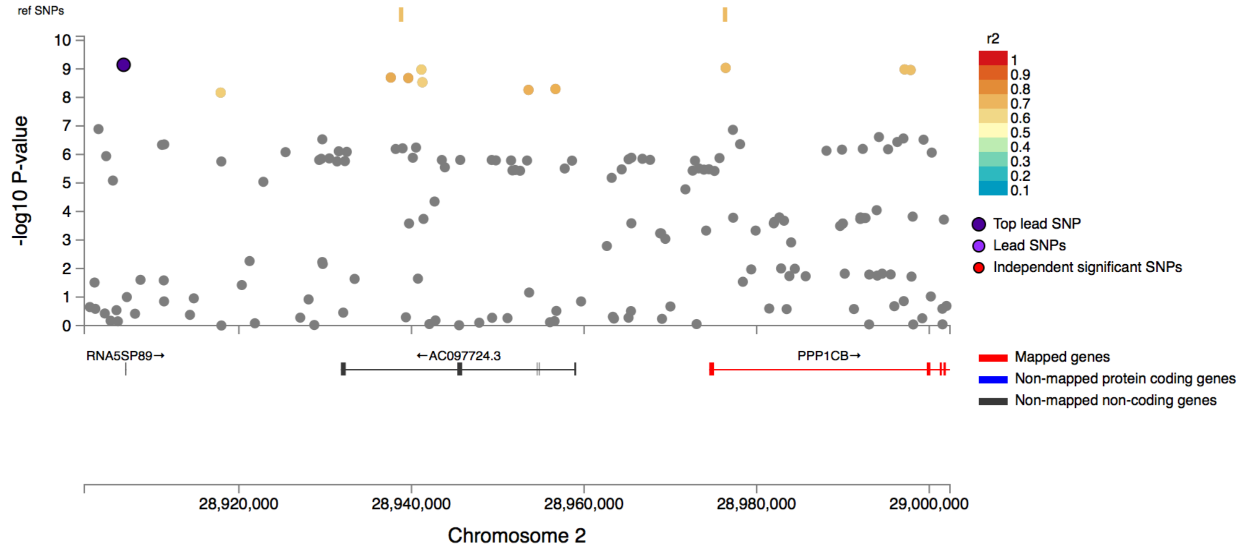

rs3024293

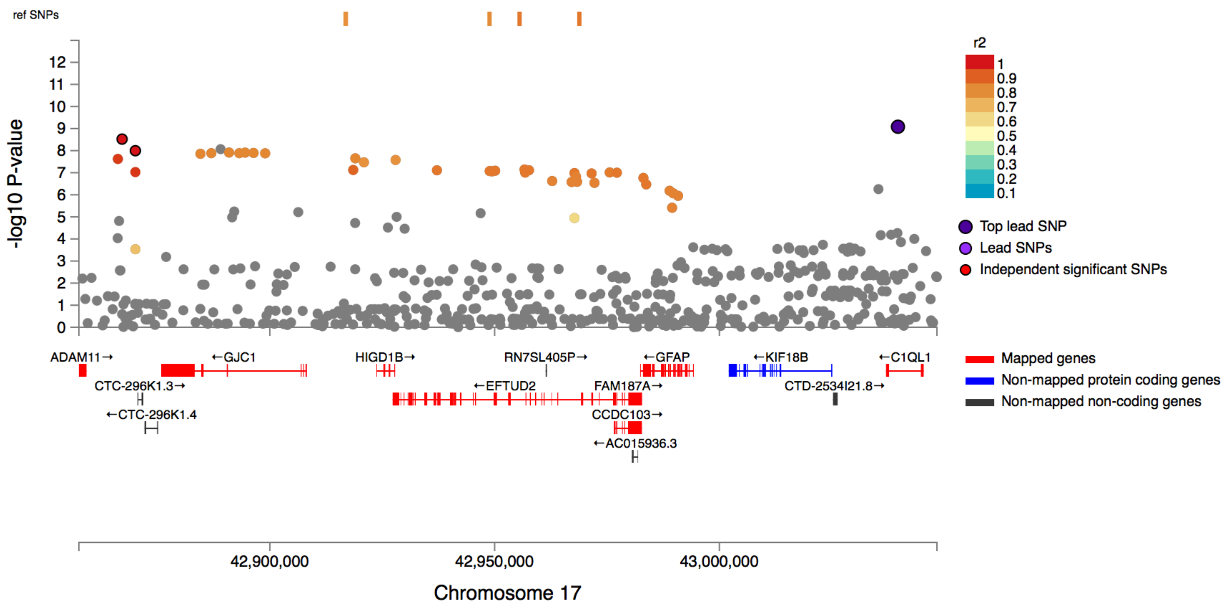


rs2061843

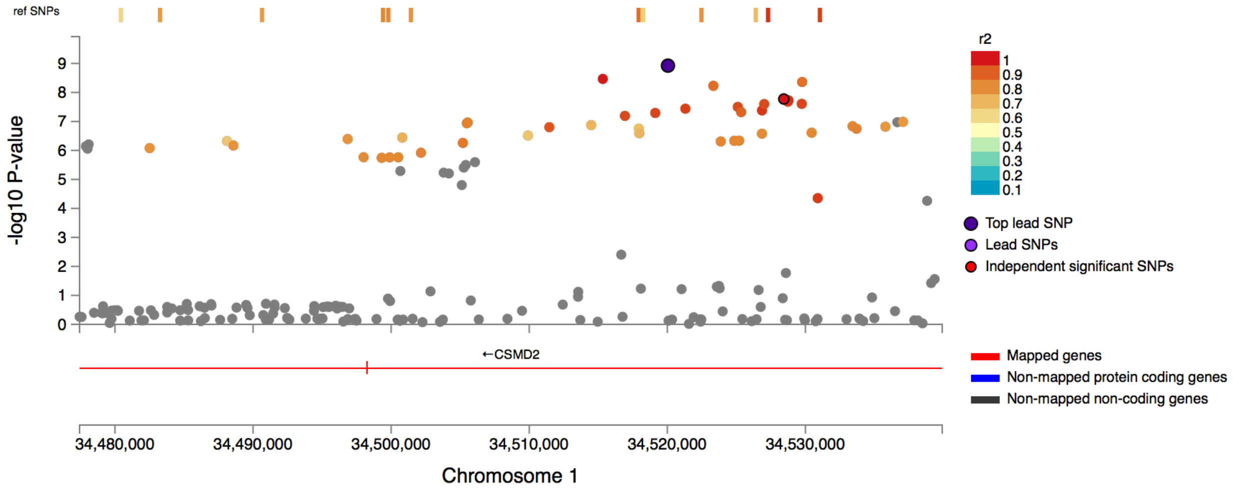

rs1349028

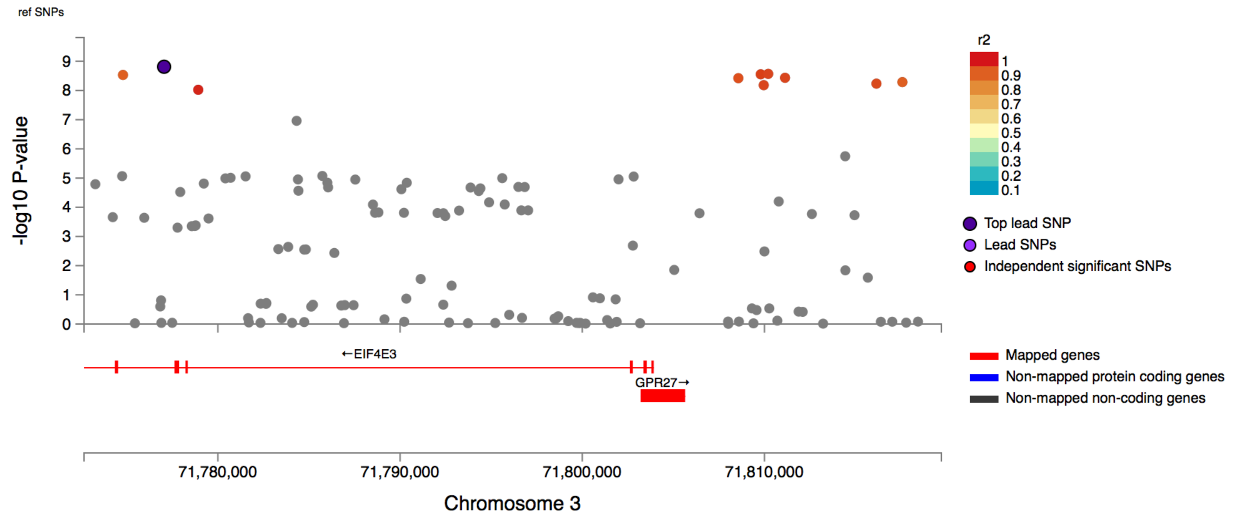


rs4443239

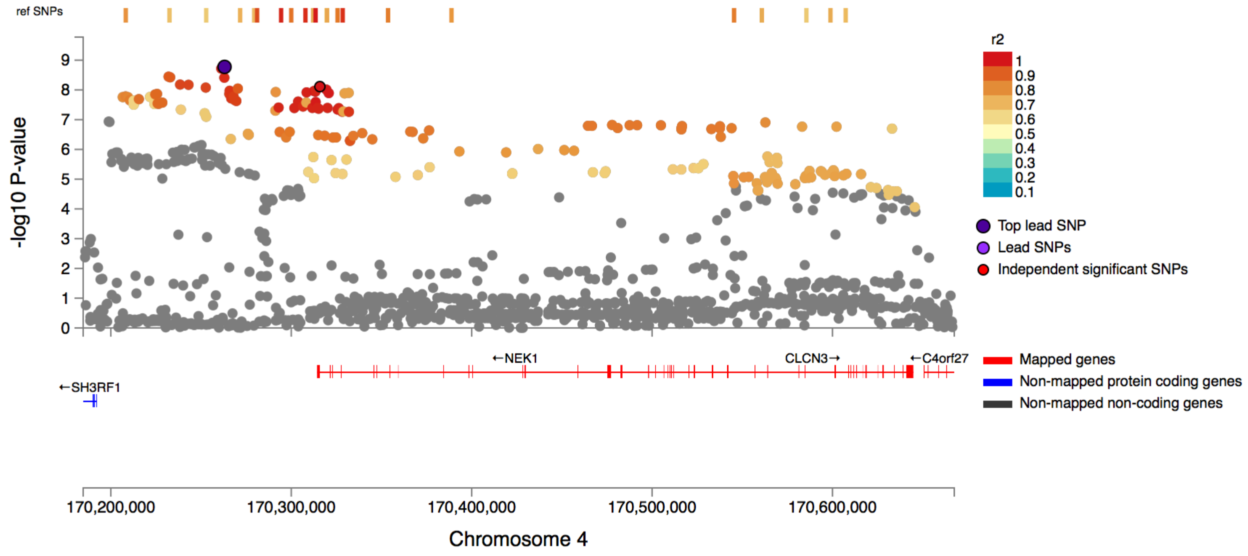

rs1901739

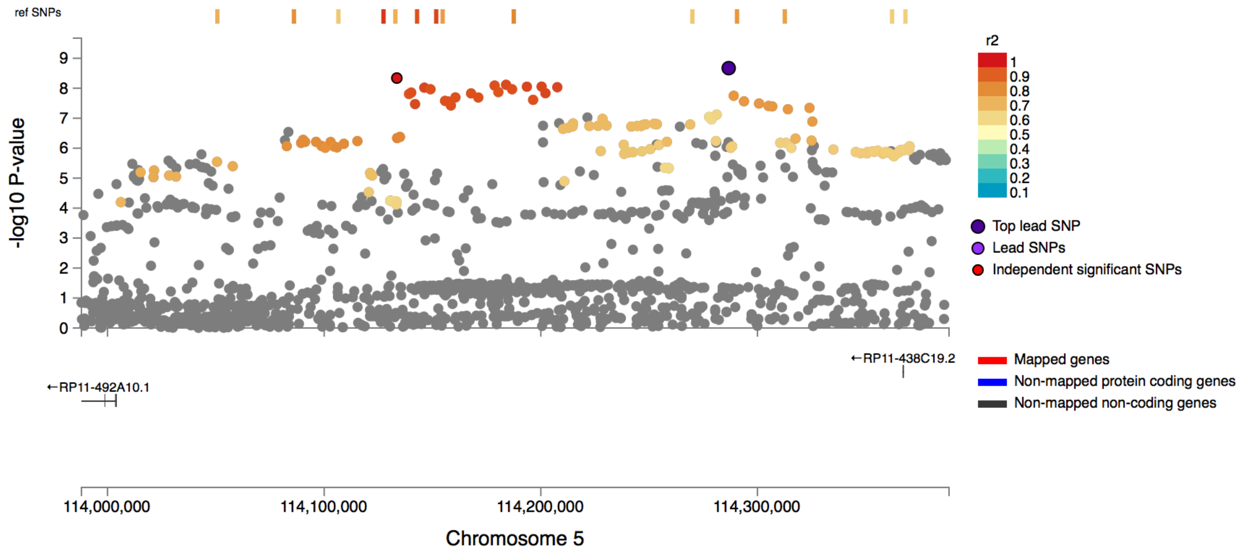


rs55678522

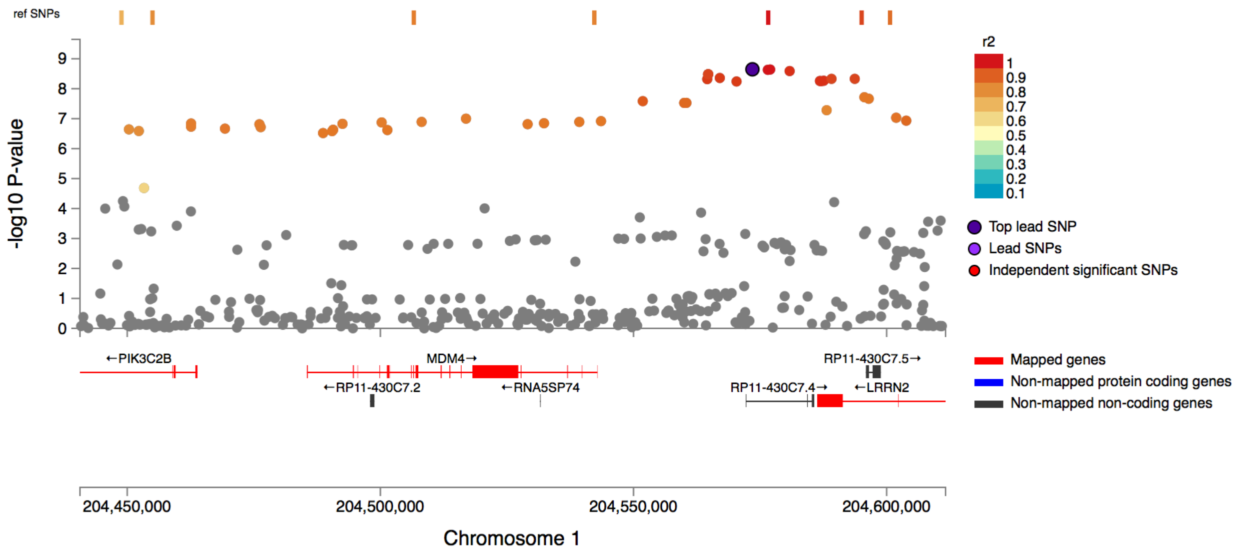

rs8079923

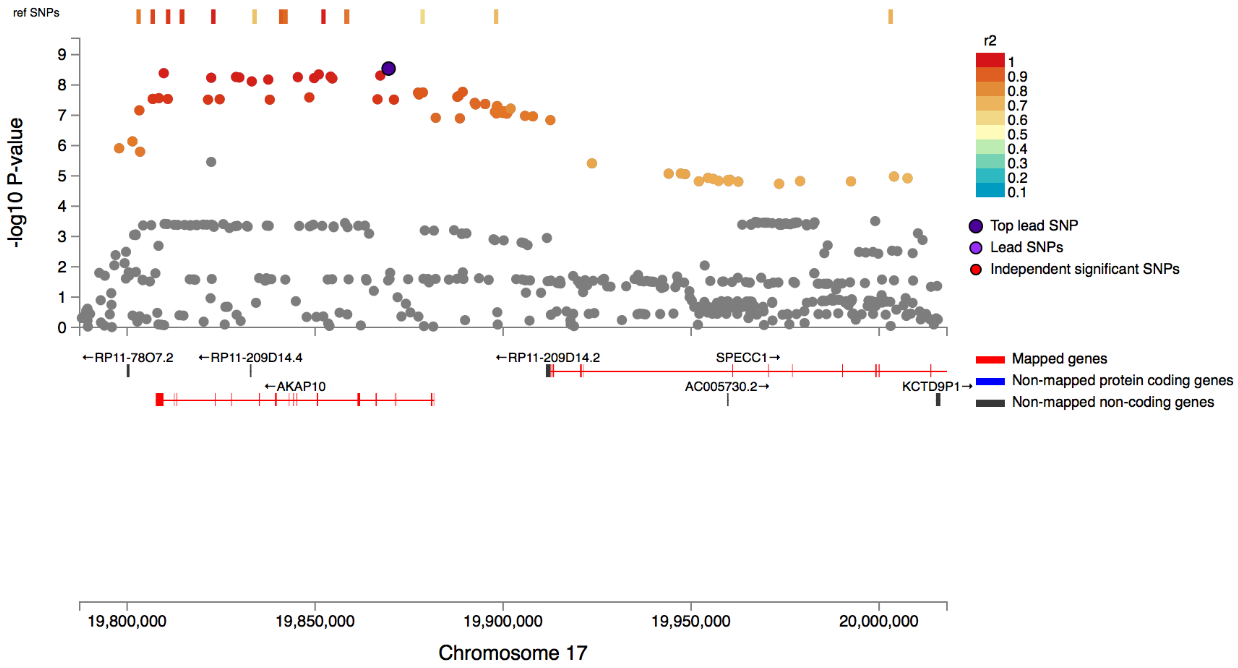


rs7501911

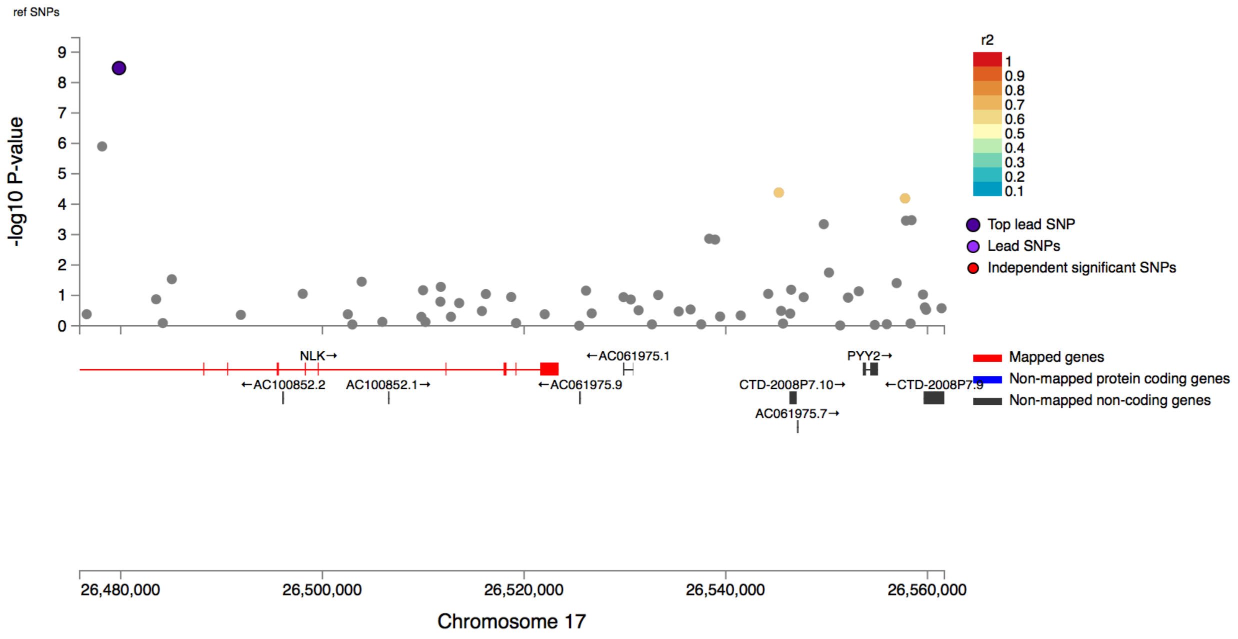

rs6087848

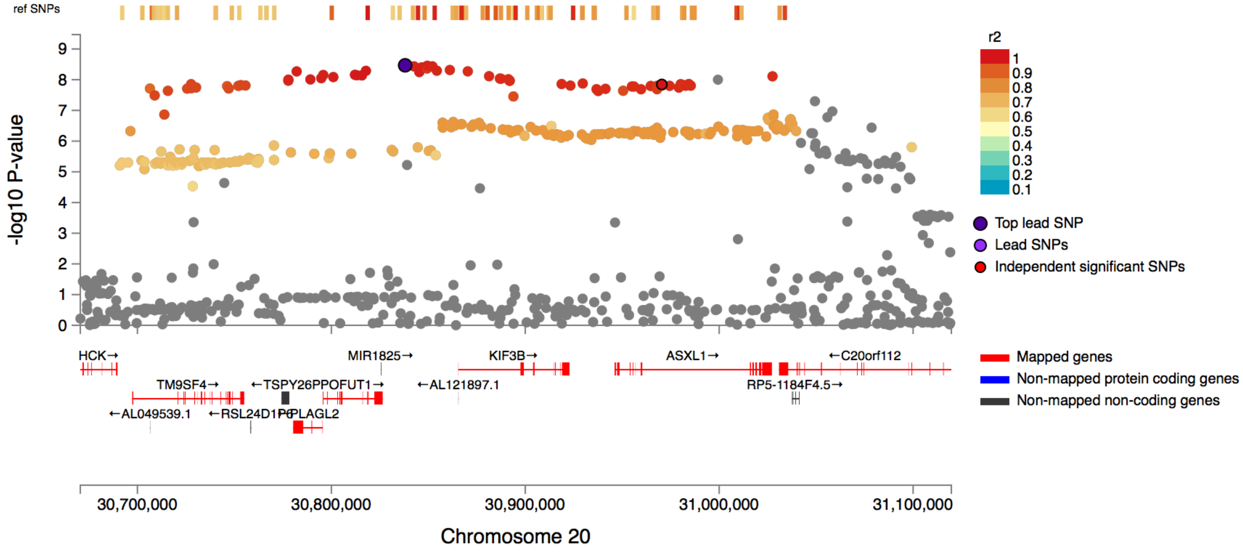


rs10744255

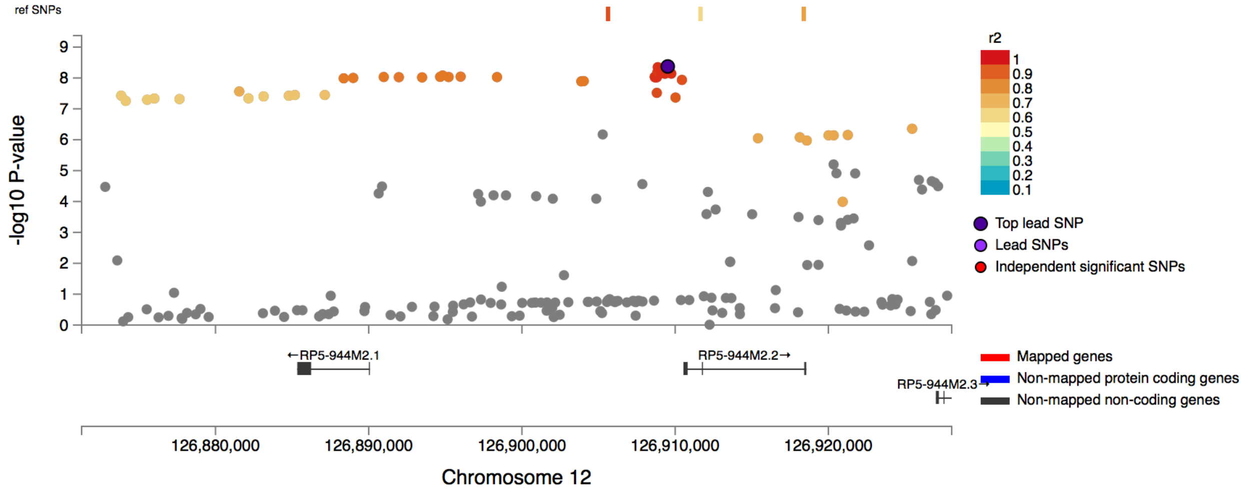

rs13163173

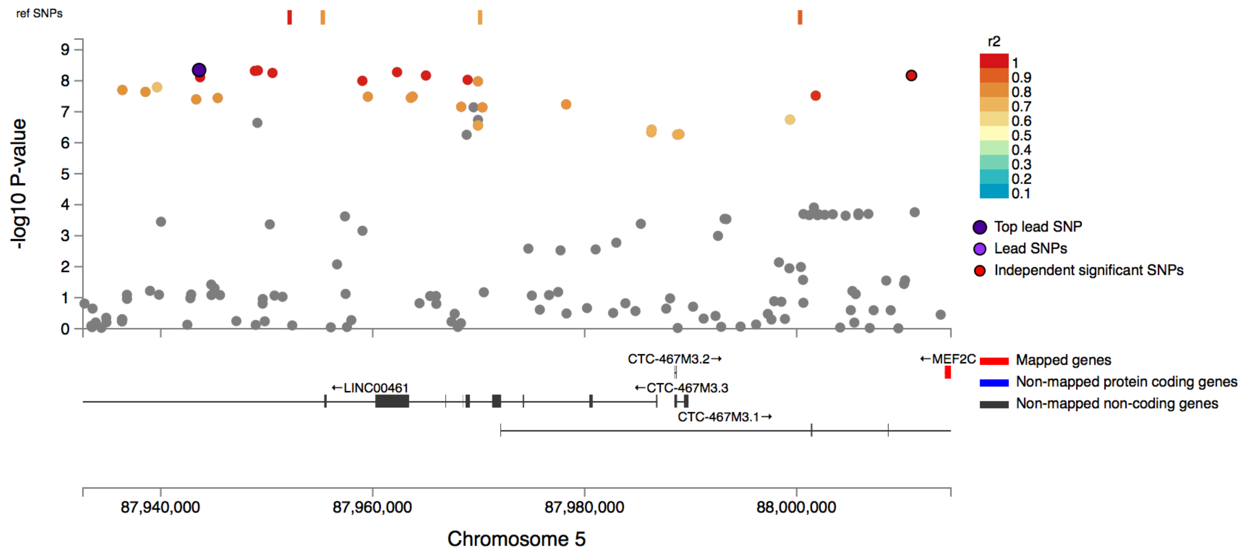


rs2819333

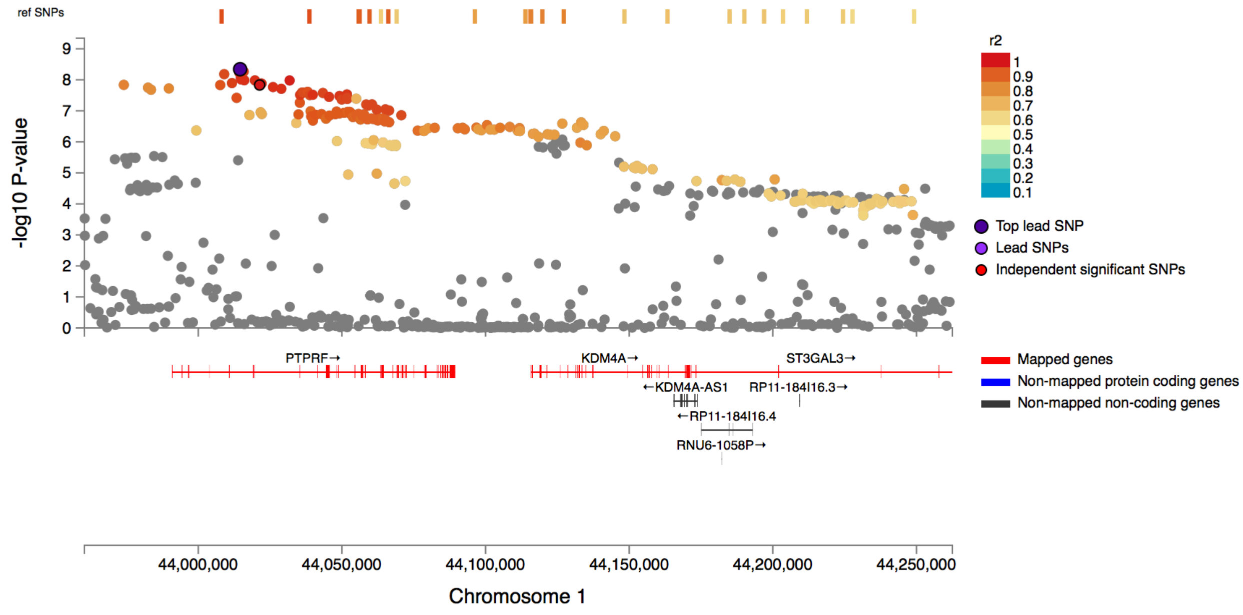

rs2453873

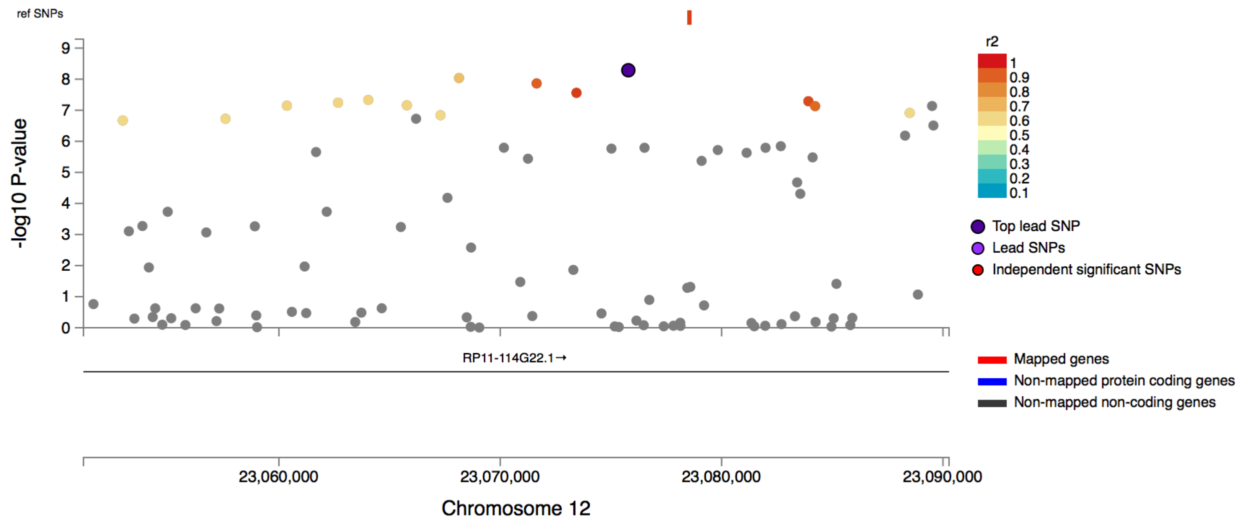


rs67264739

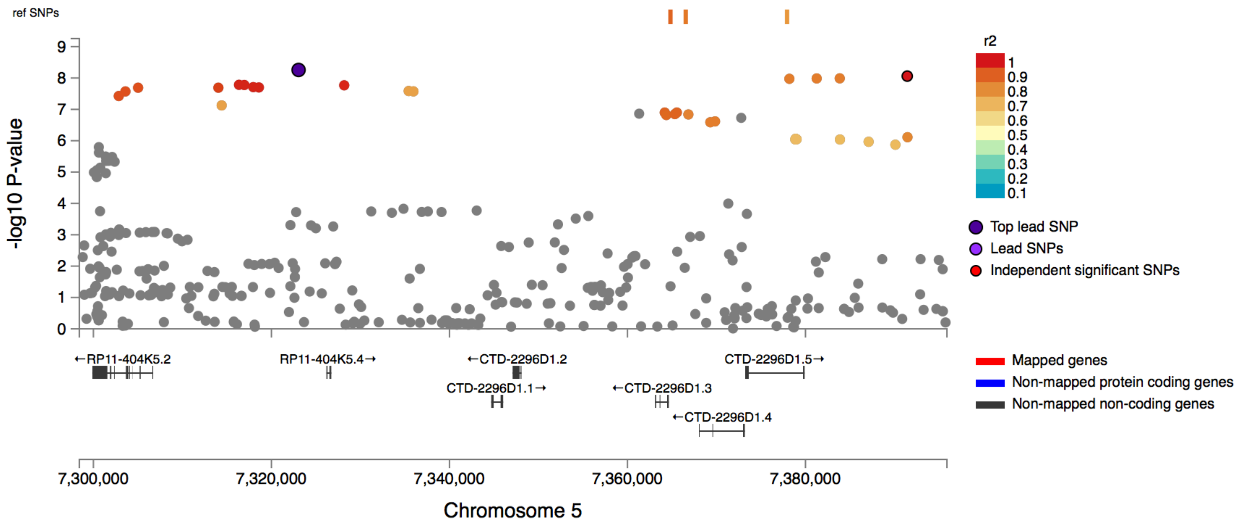

rs4898322

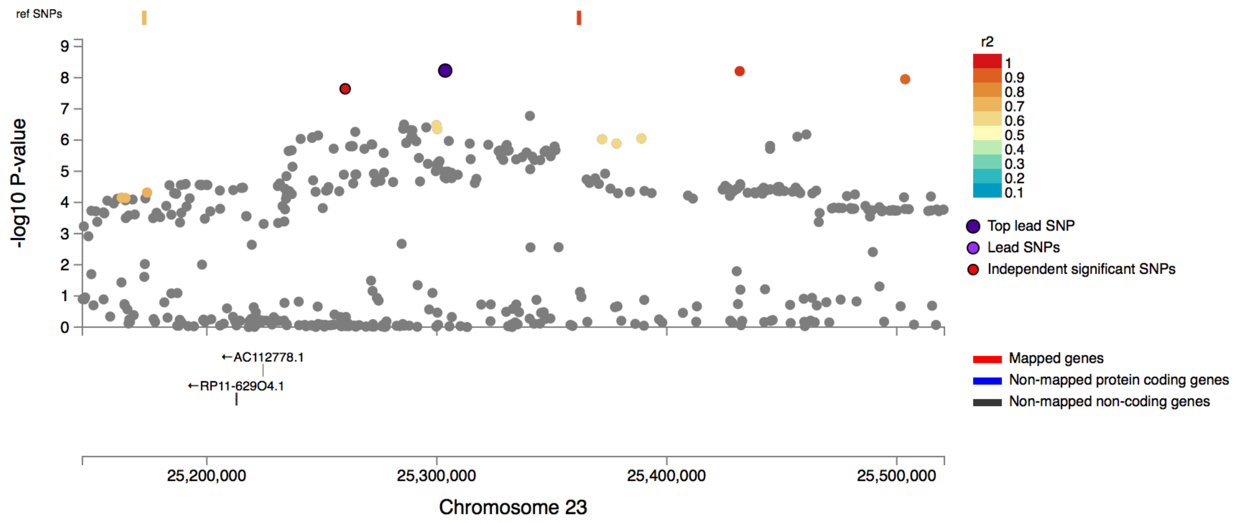


rs2284901

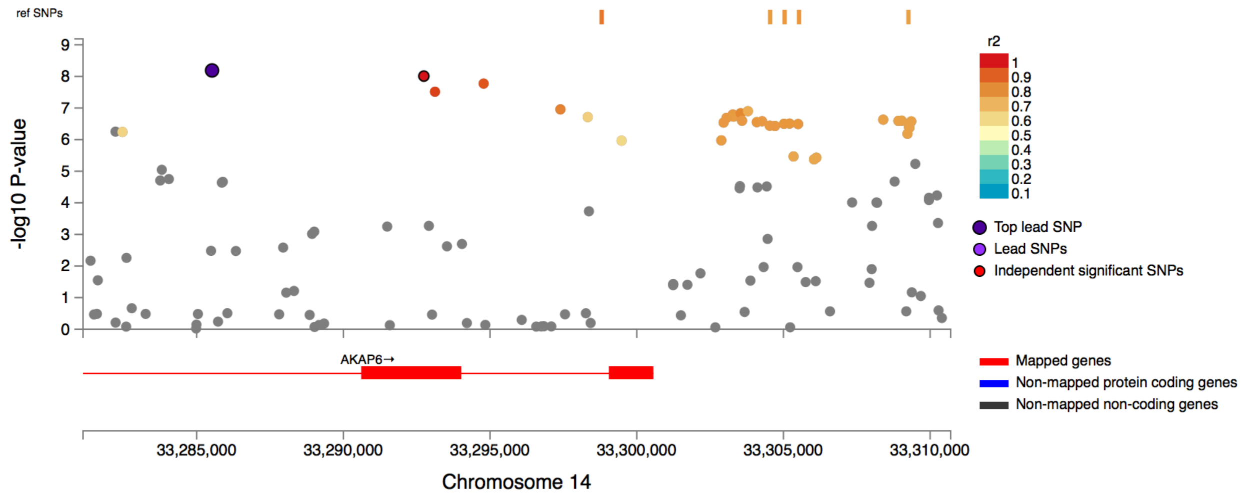

rs1596431

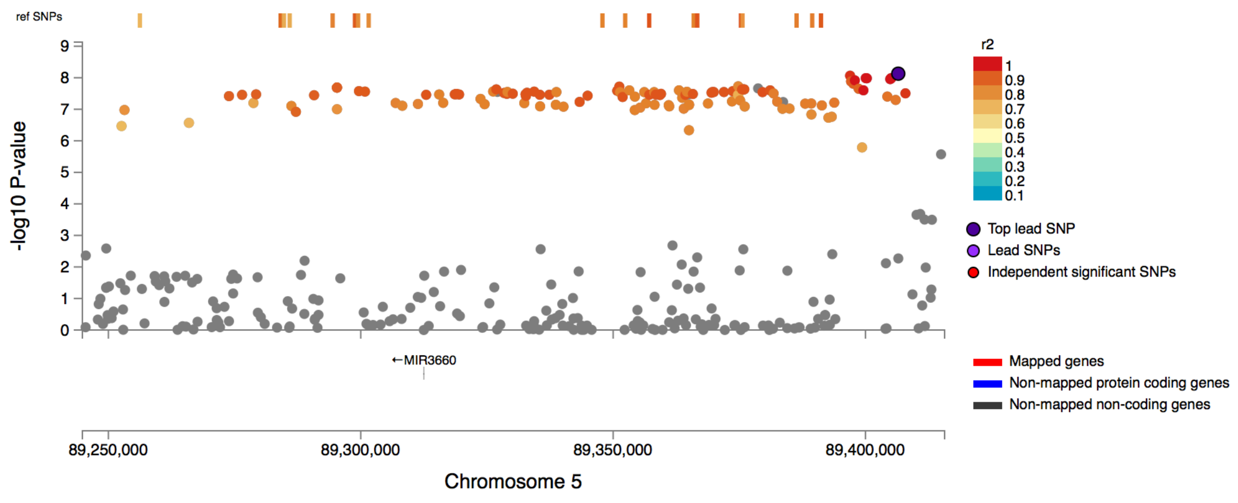


rs10978661

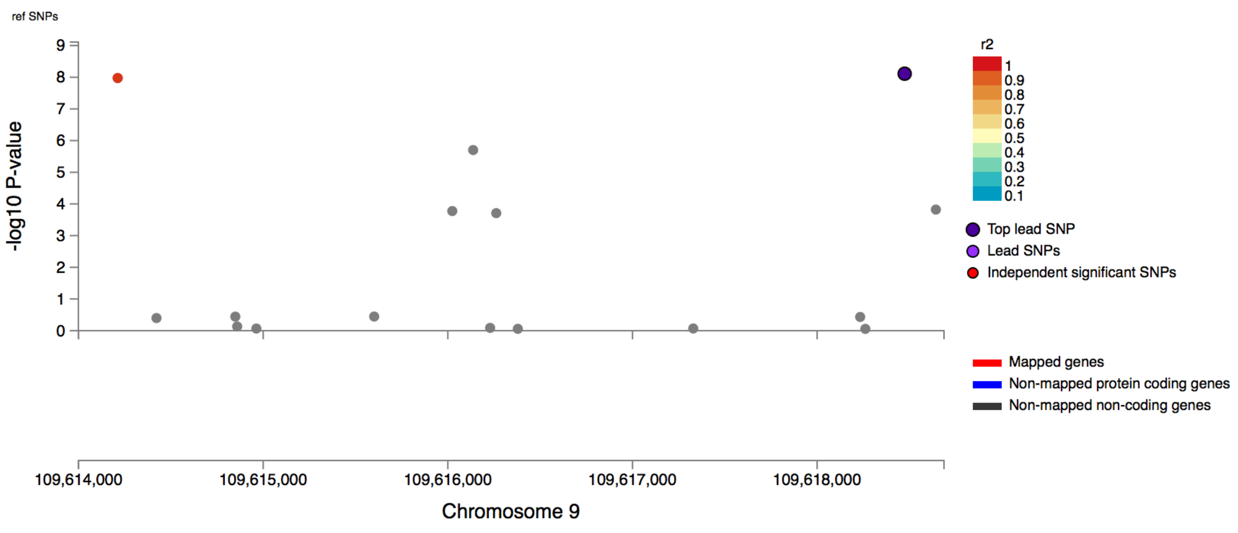

rs4263335

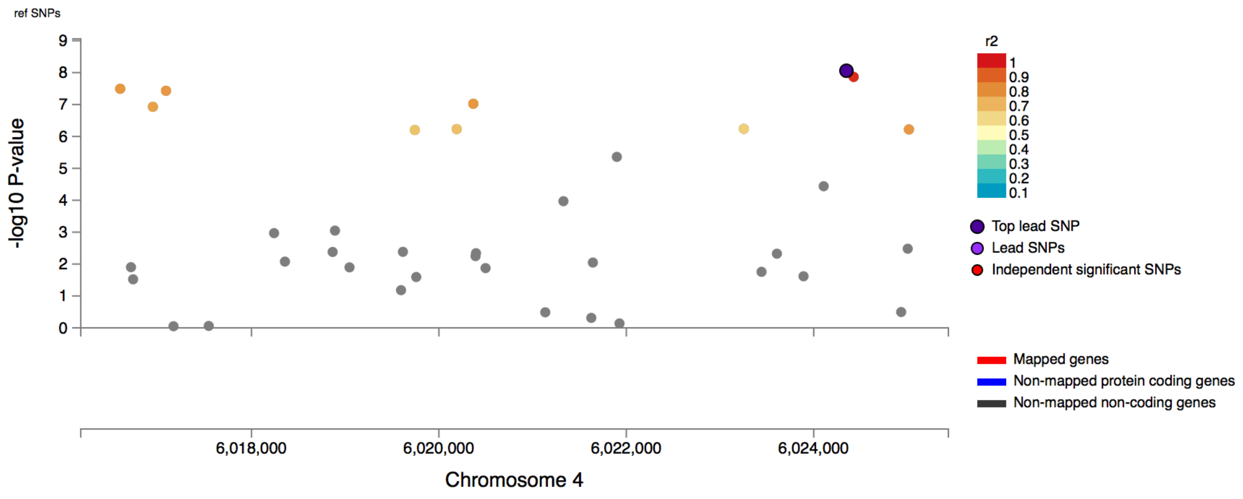


rs7939759

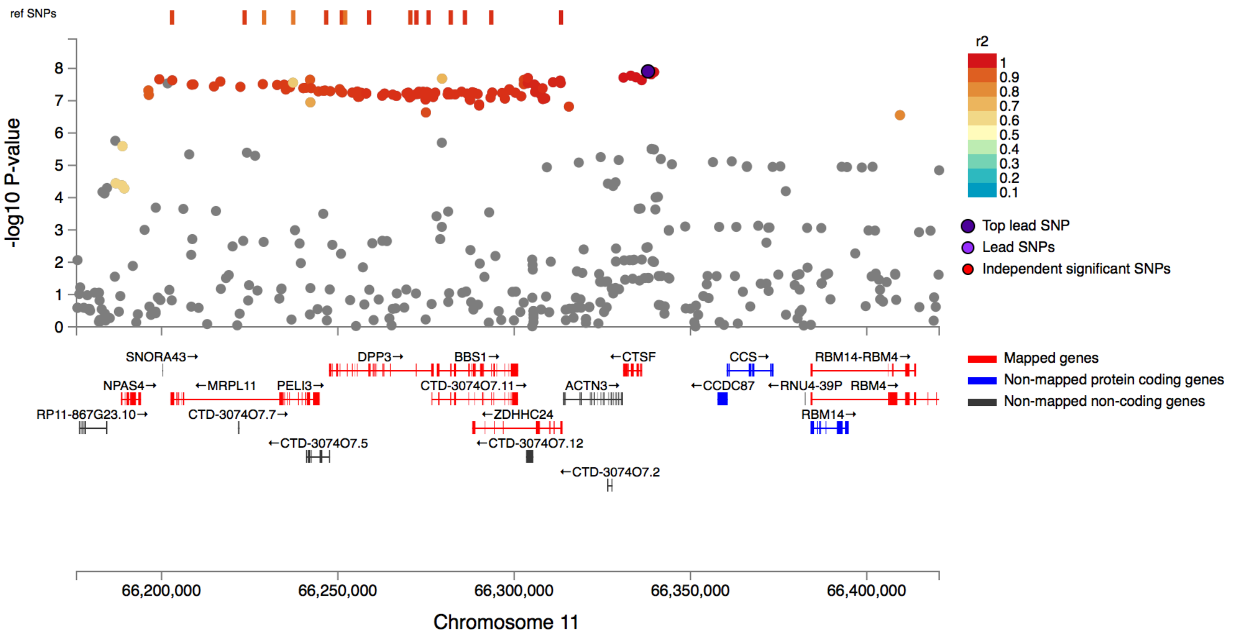

rs9710427

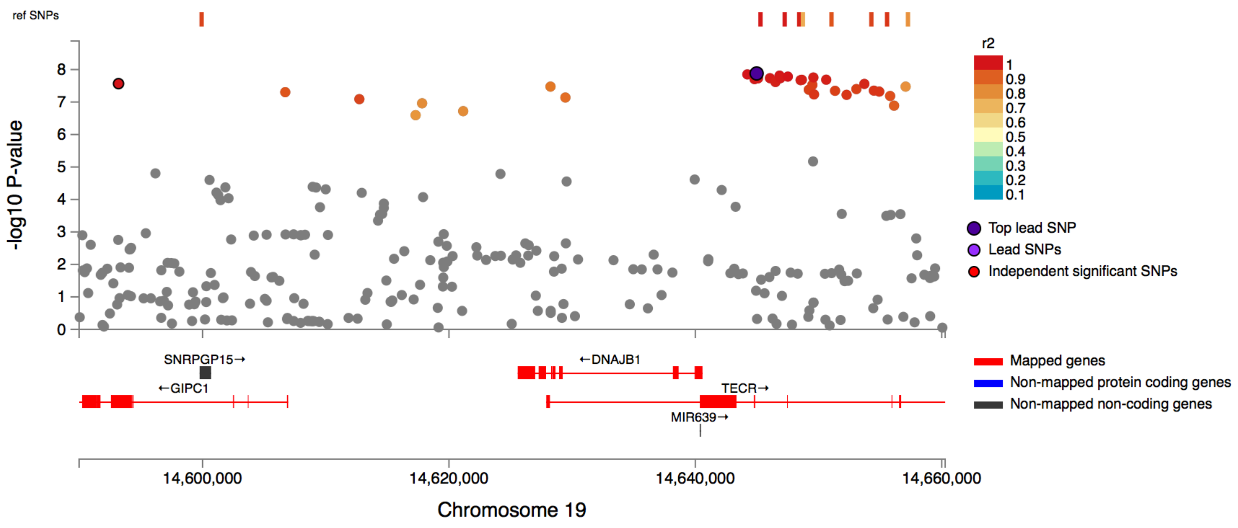


rs12638746

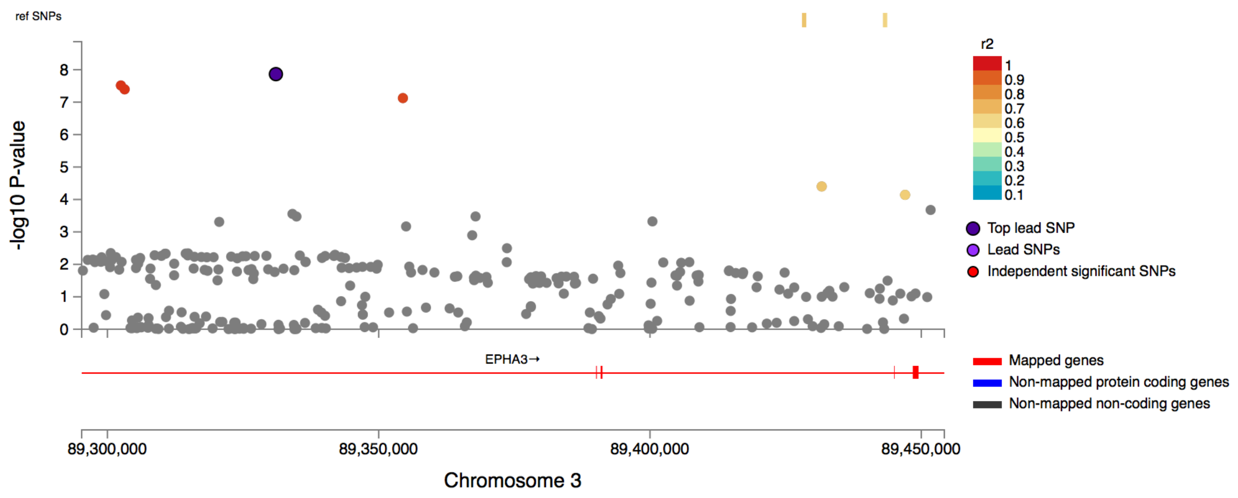

## rs12909047

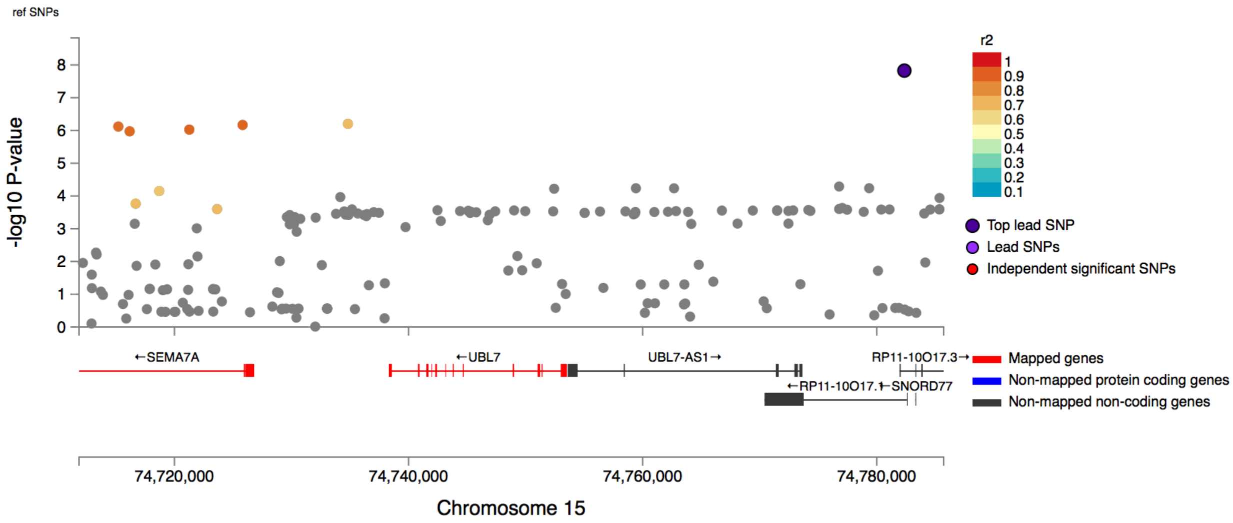


## rs2505344

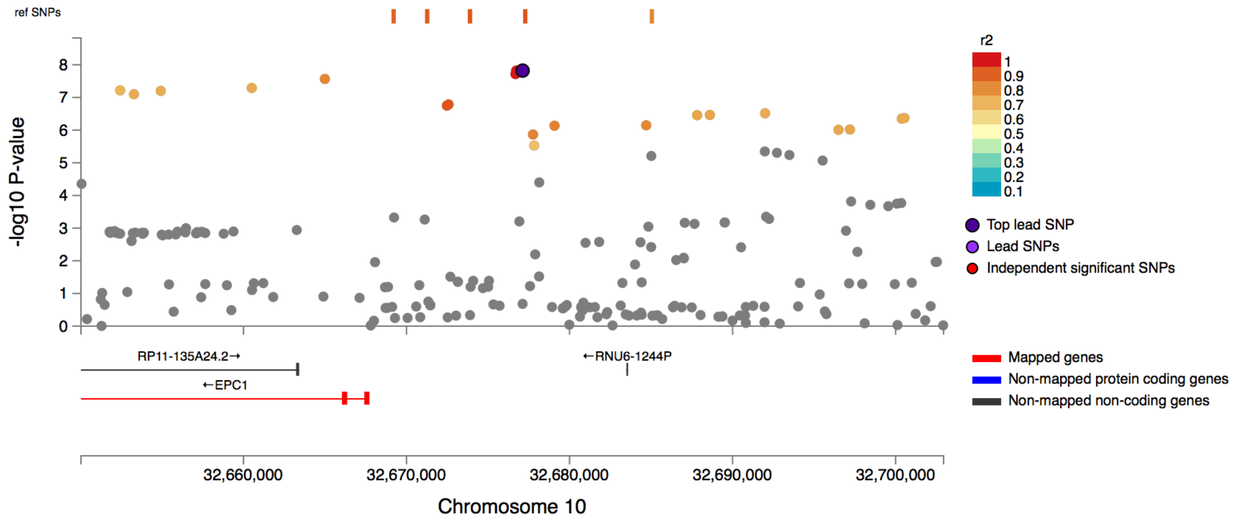

rs67816799

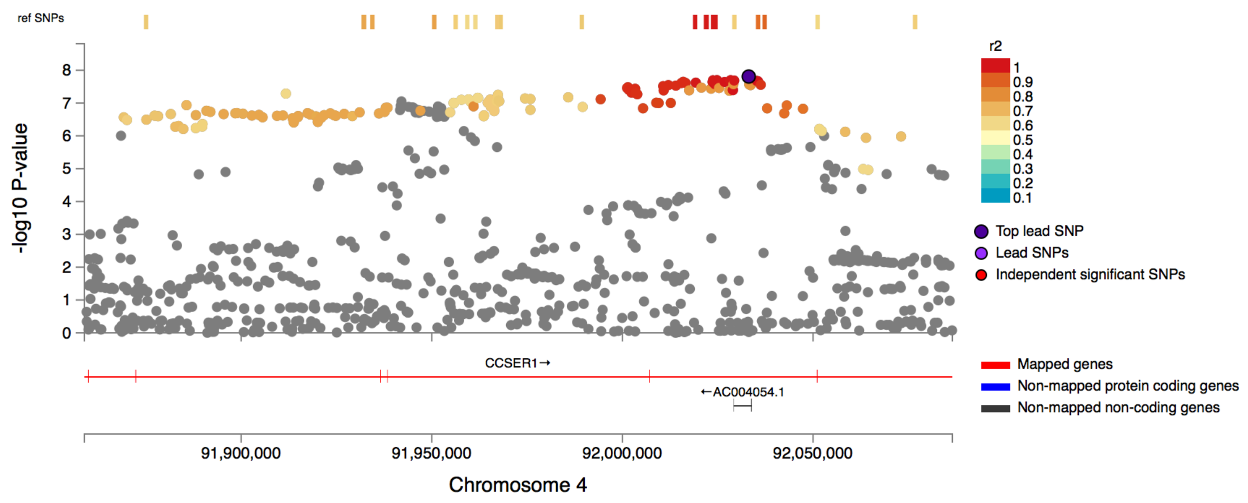


rs10932201

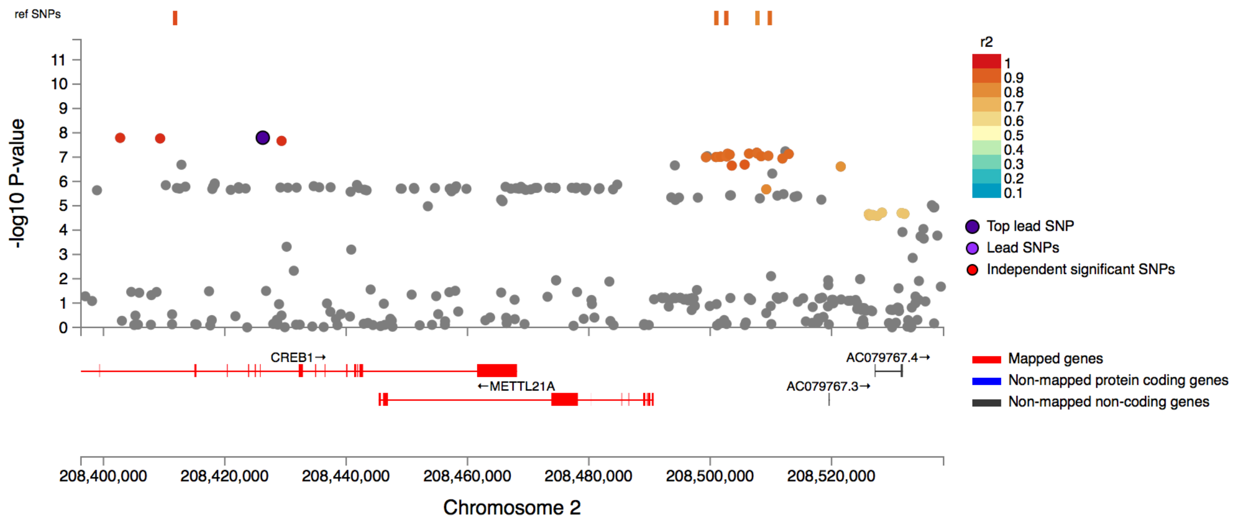

rs526904

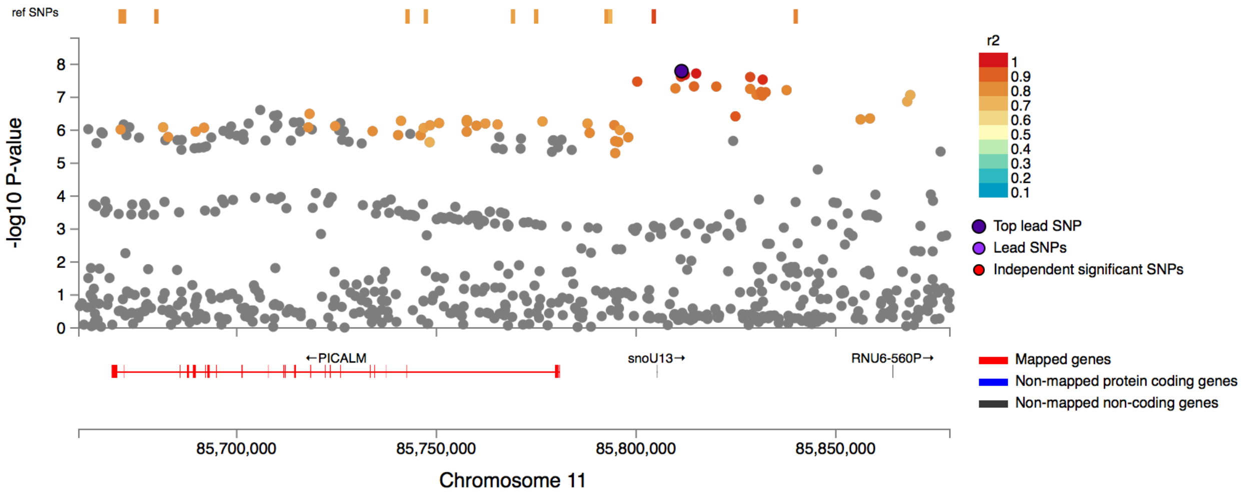


rs764935655

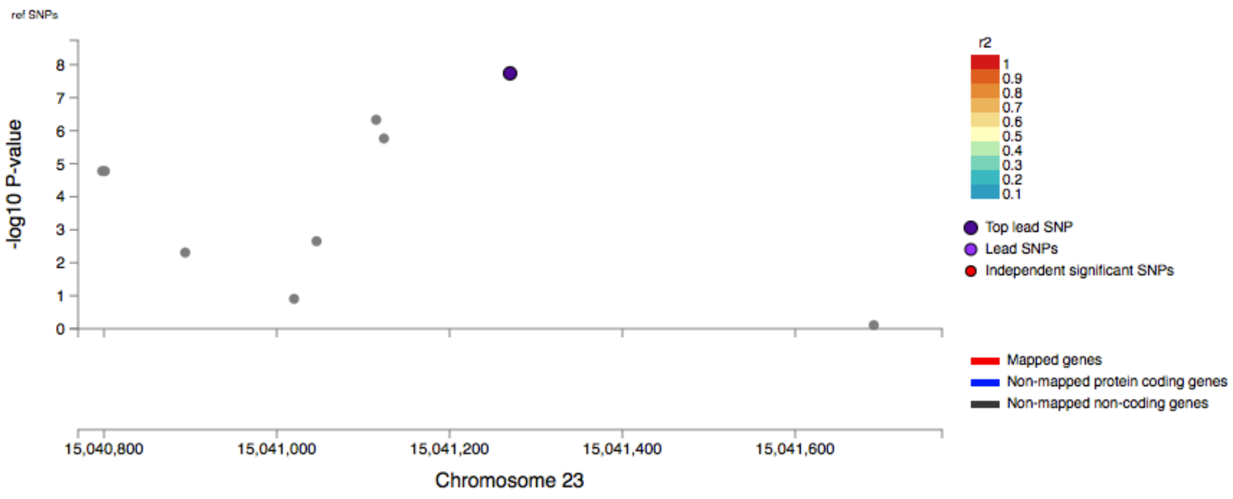

## rs6548147

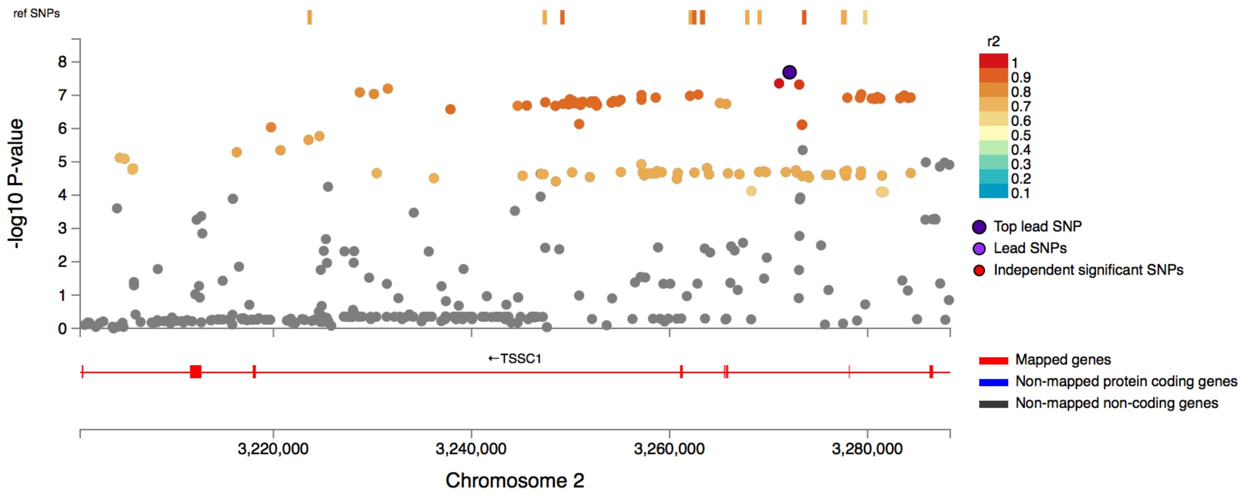


## rs10877461

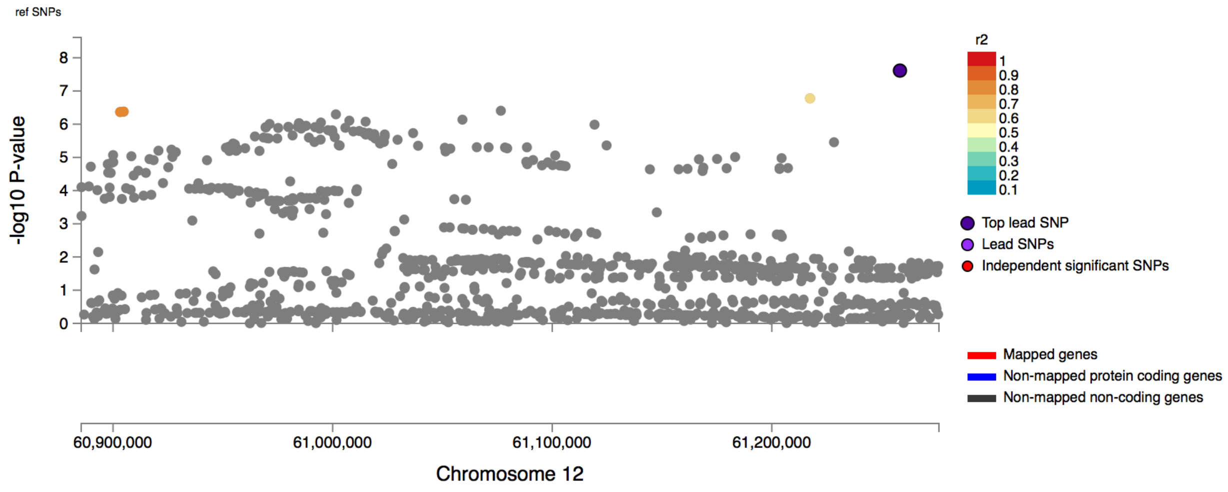

rs11996434

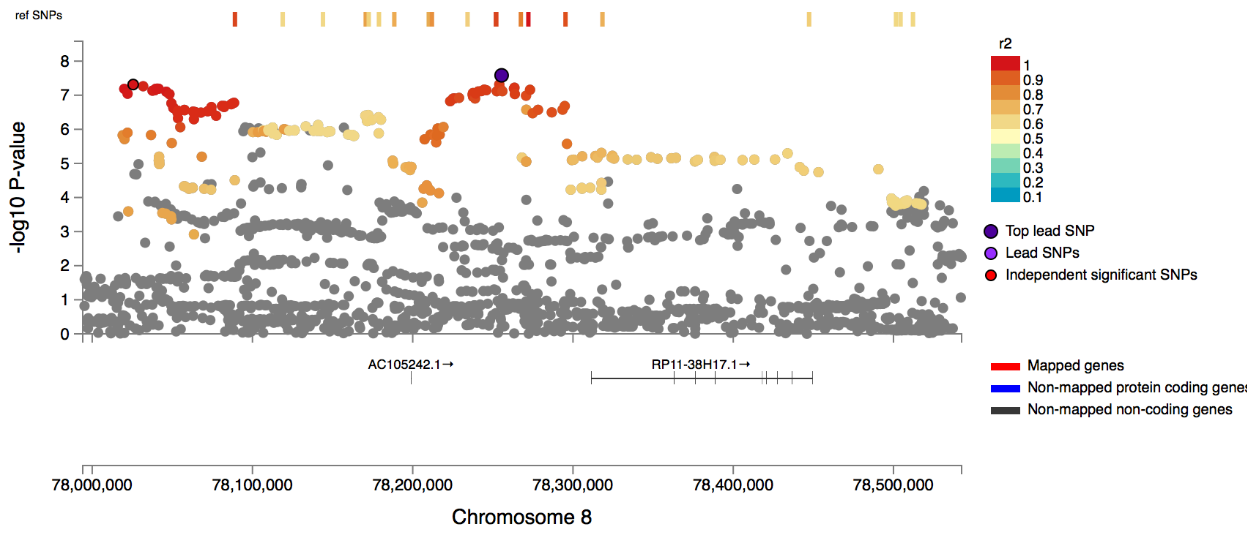


rs10885458

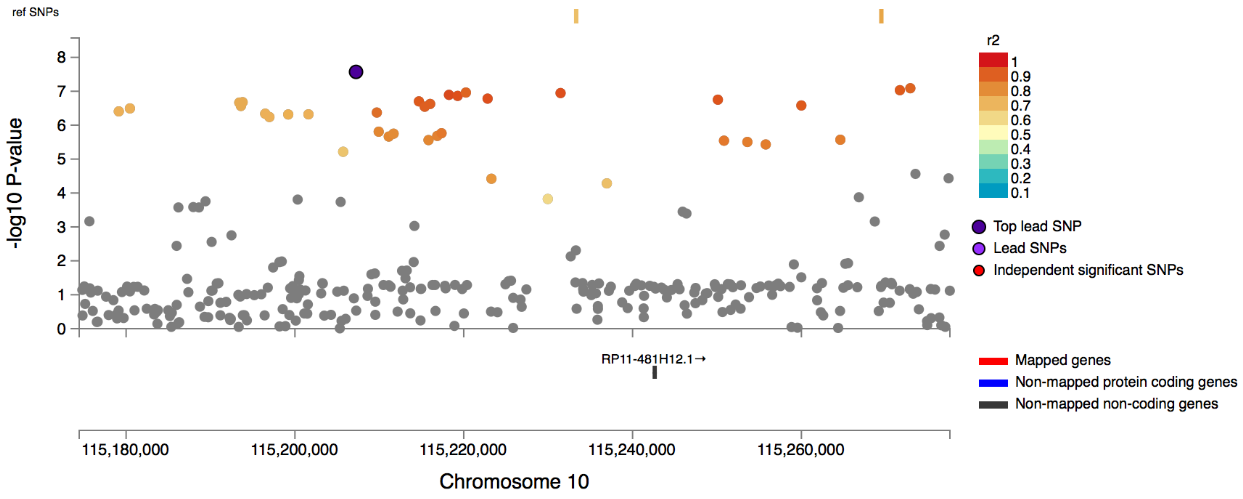

rs1996148

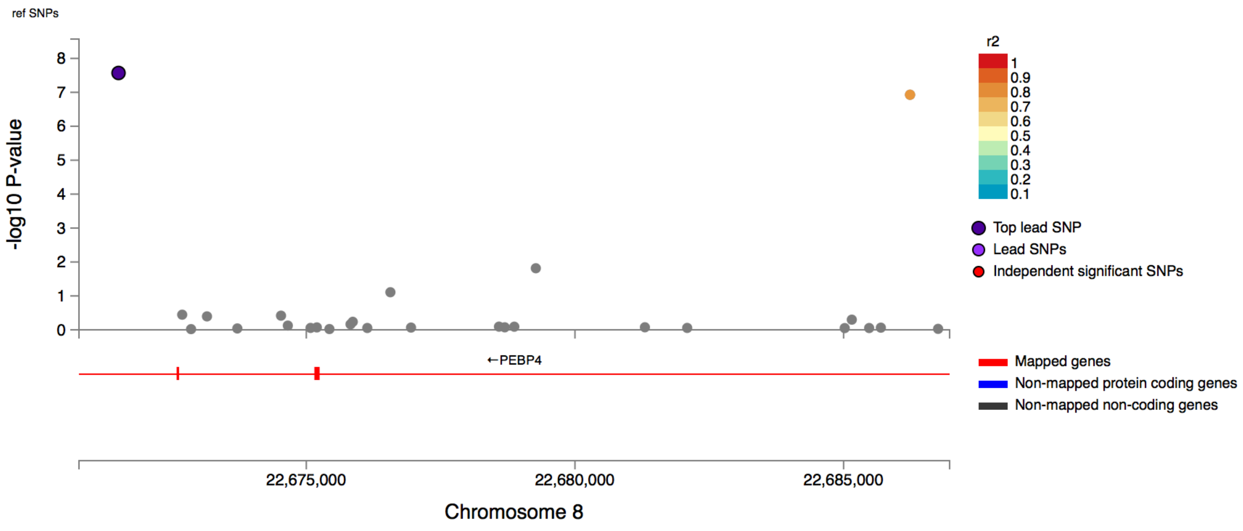


rs191373913

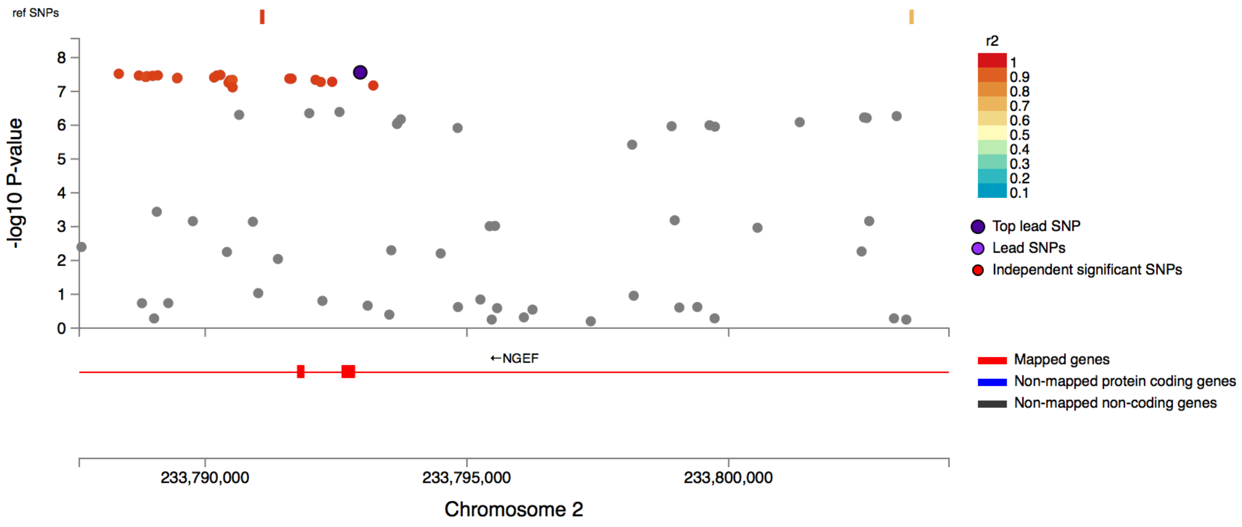

rs12056186

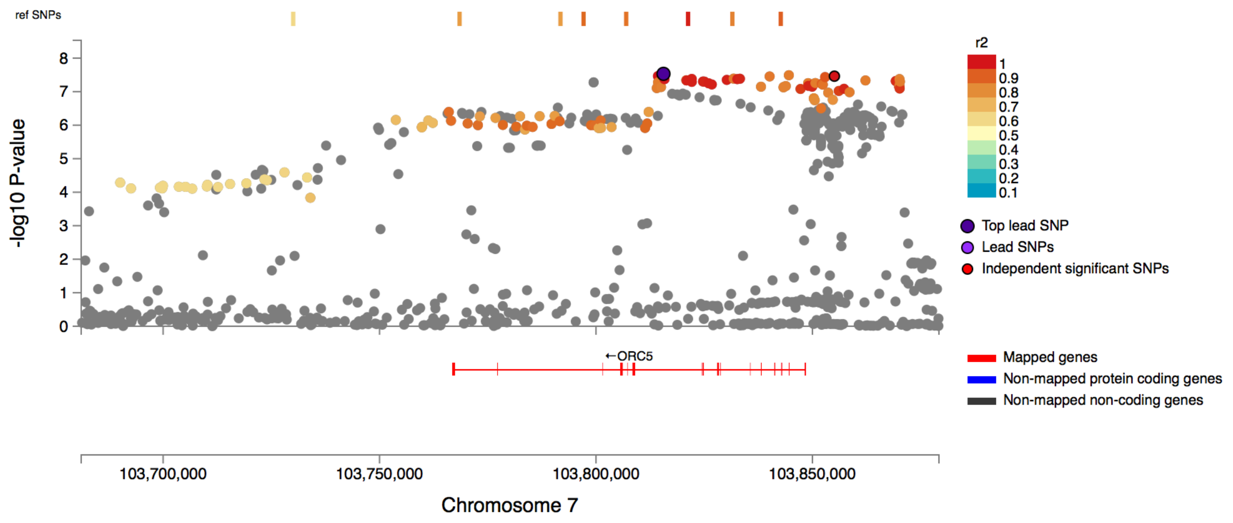


rs7856850

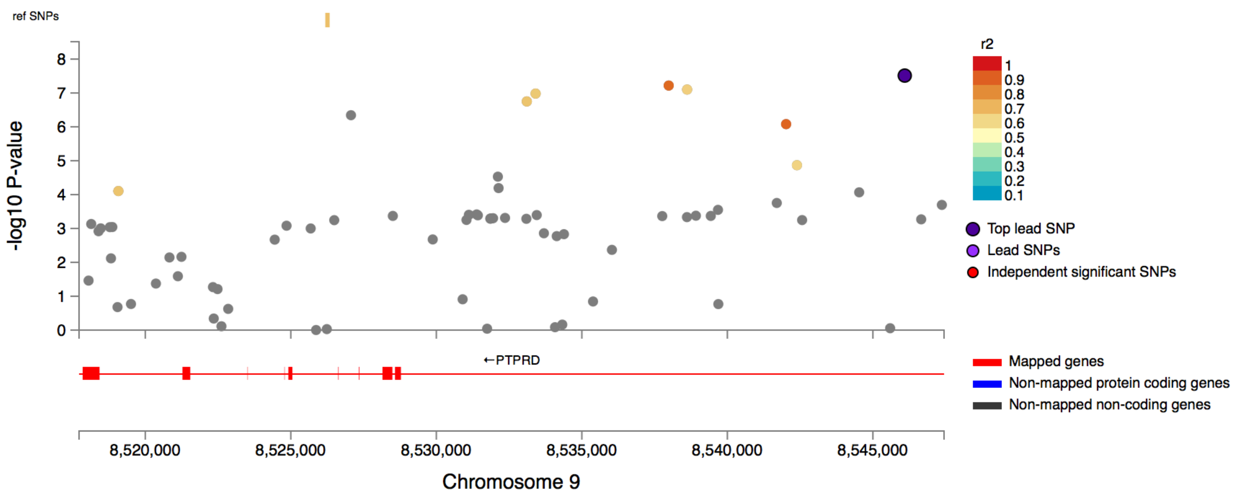

rs13197257

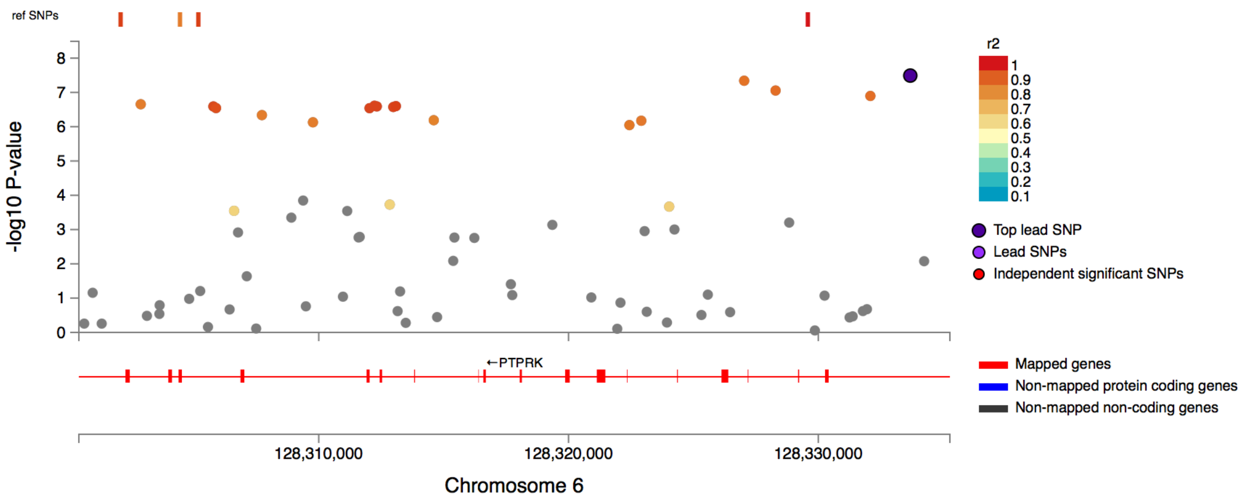


rs10497355

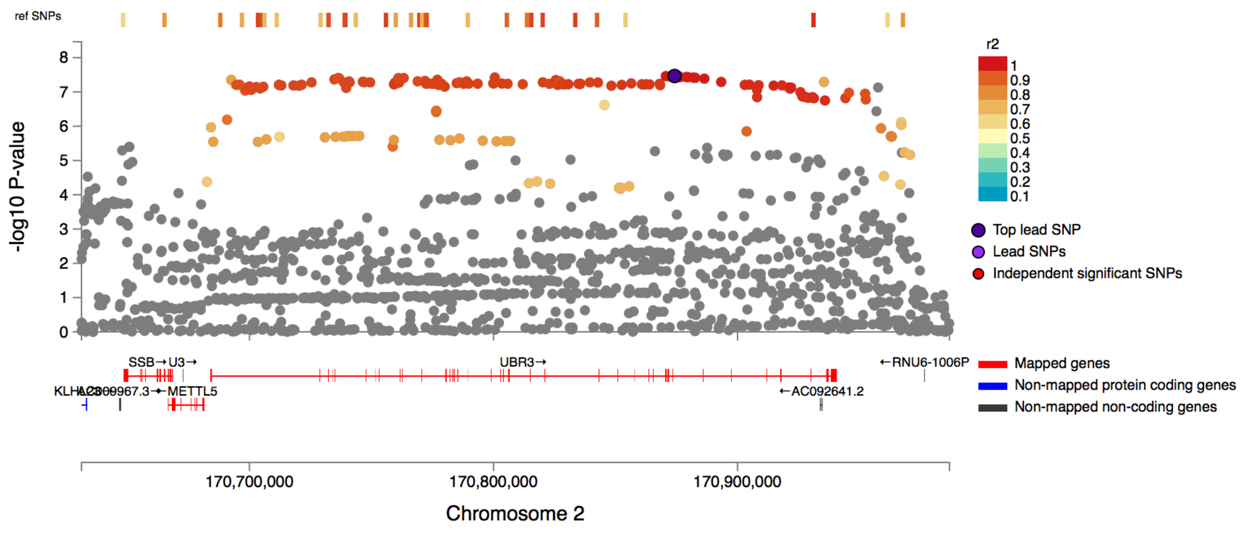

rs11692449

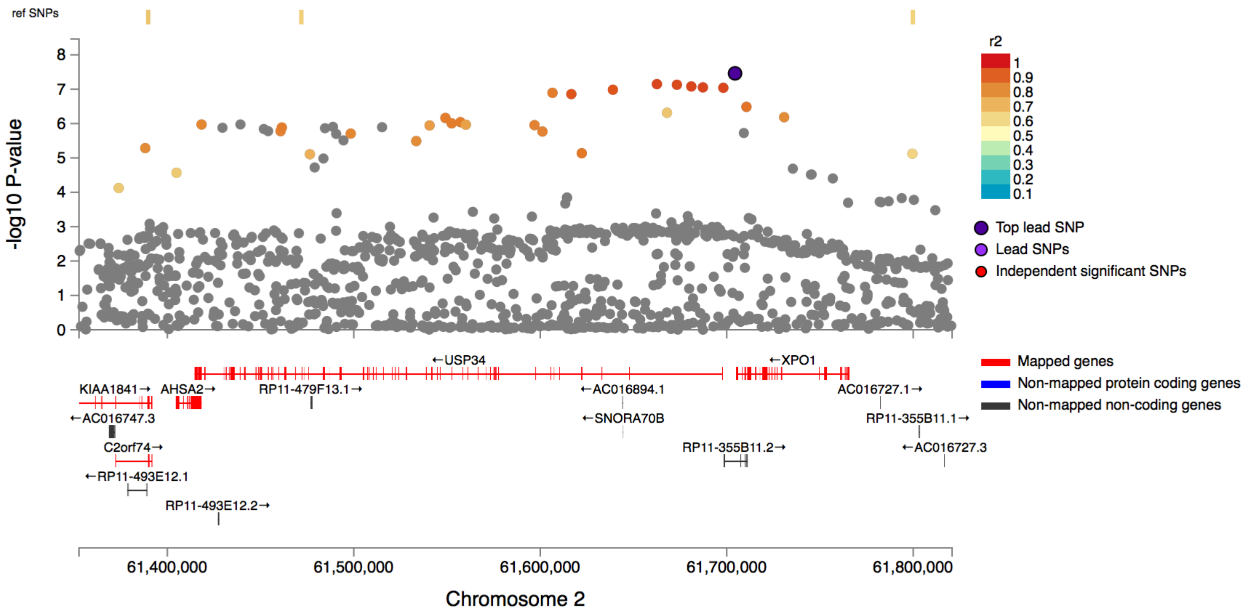


rs4704043

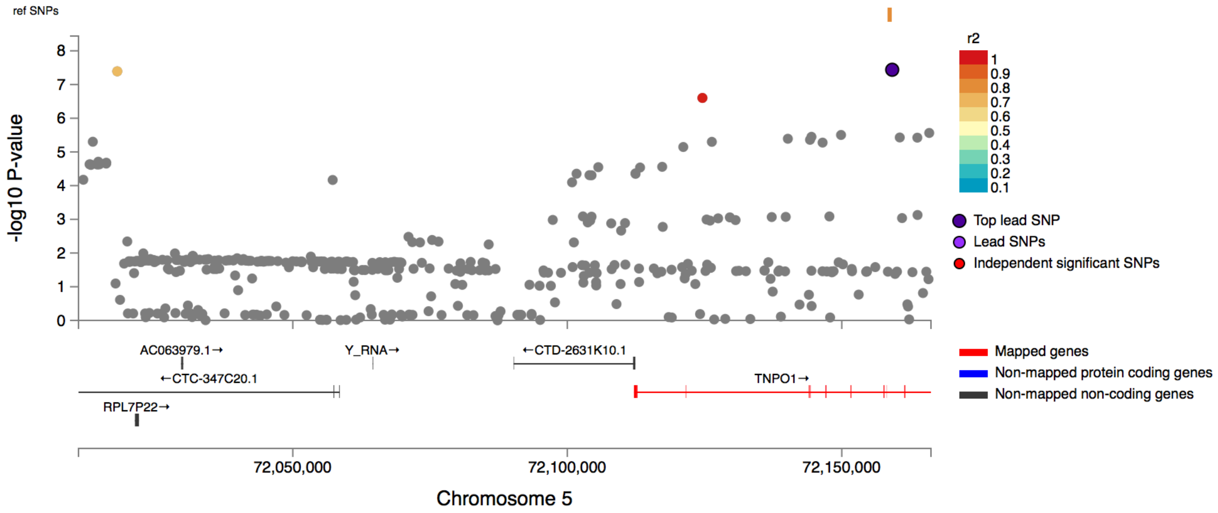

rs43182

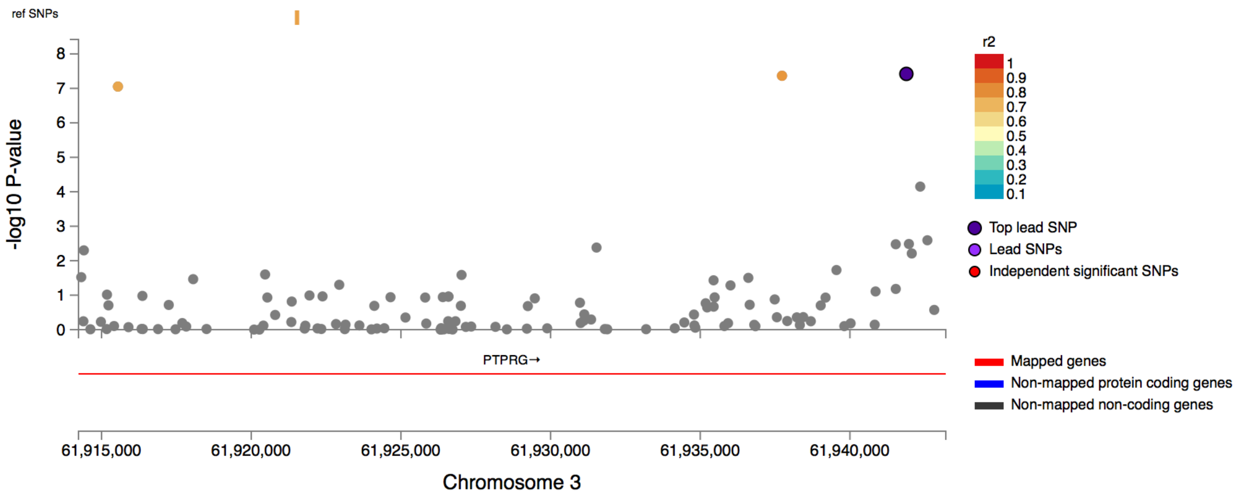


rs62014217

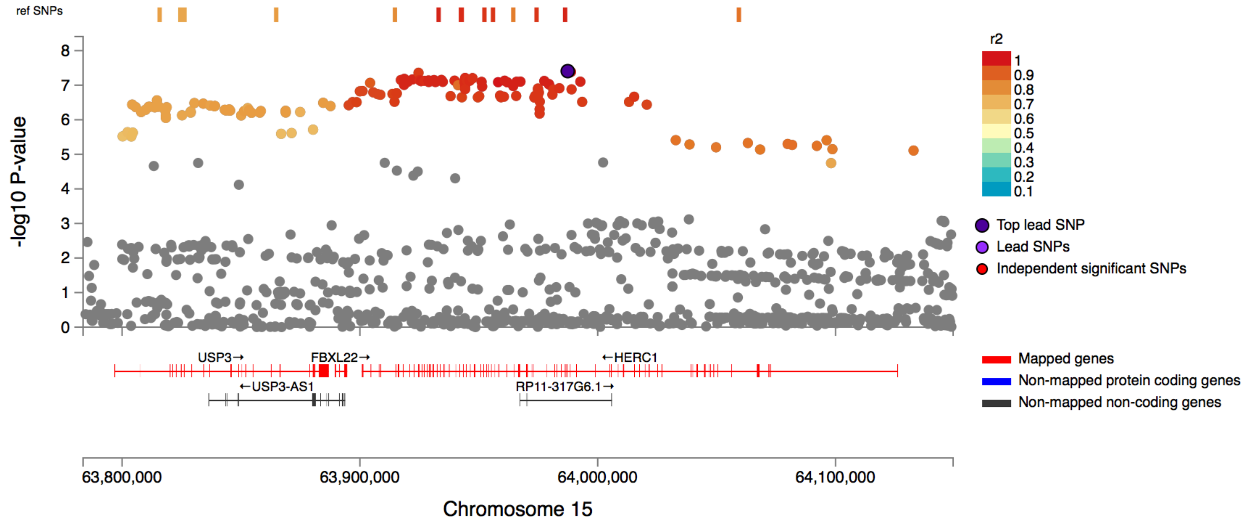

rs476141

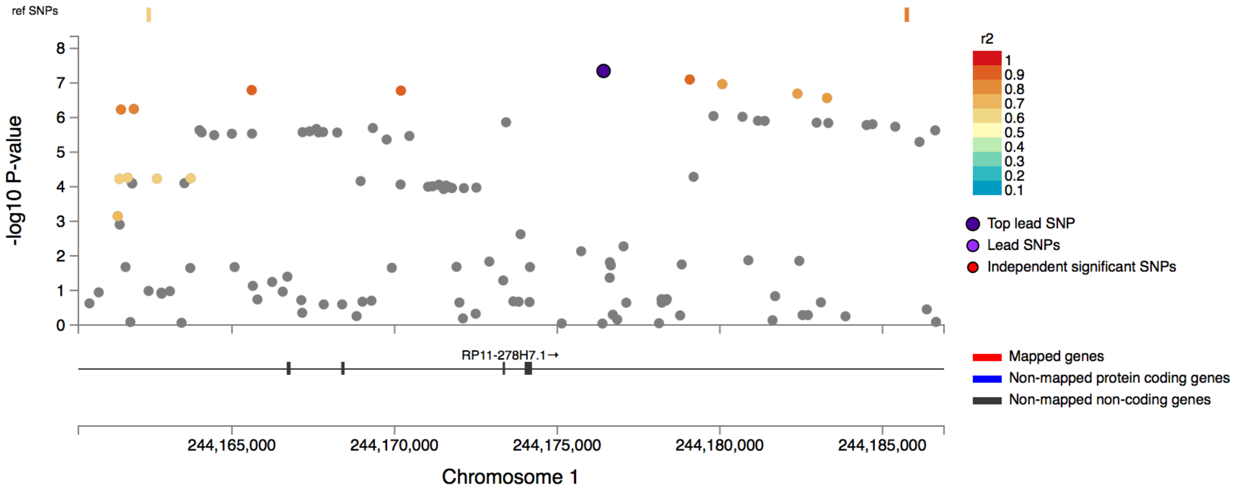


rs2849543

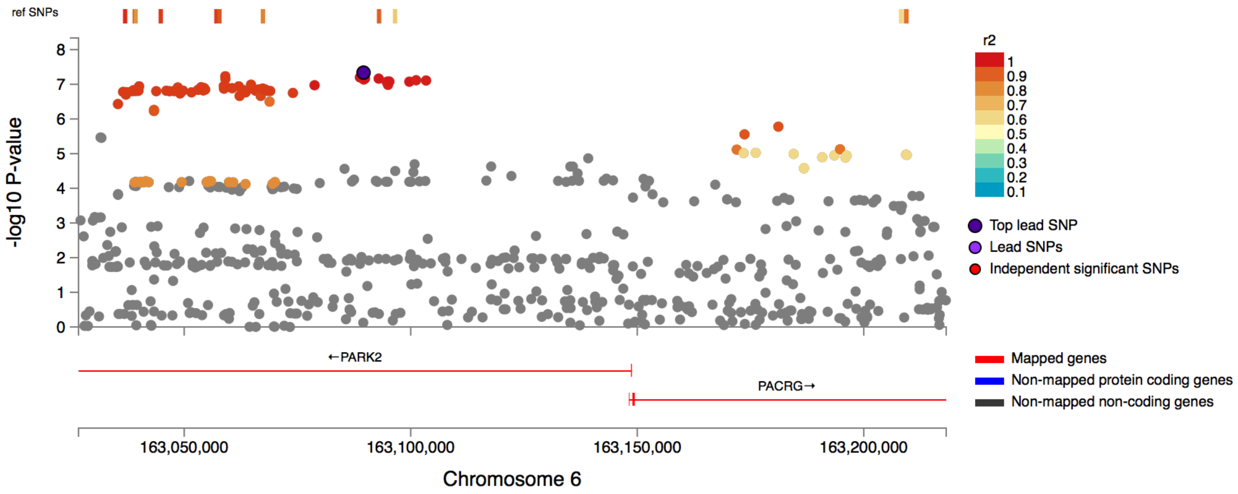

rs571760466

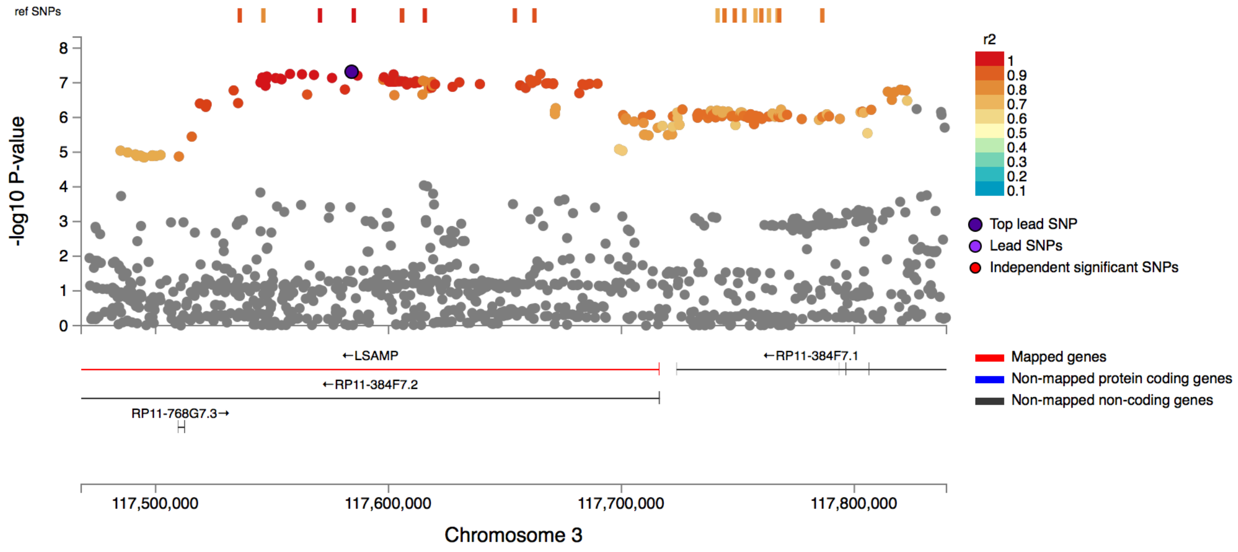
